# Fine decomposition of rodent behavior via unsupervised segmentation and clustering of inertial signals

**DOI:** 10.1101/2024.09.20.613901

**Authors:** Romain Fayat, Marie Sarraudy, Clément Léna, Daniela Popa, Pierre Latouche, Guillaume P. Dugué

## Abstract

Decomposing behavior into elementary components remains a central challenge in computational neuroethology. The current standard in laboratory animals involves multi-view video tracking, which, while providing unparalleled access to full-body kinematics, imposes environmental constraints, is data-intensive, and has limited scalability. We present an alternative approach using inertial sensors, which capture high-resolution, environment-independent, and compact three-dimensional kinematic data, and are commonly integrated into rodent neurophysiological devices. Our analysis pipeline leverages unsupervised, computationally efficient change-point detection to break down inertial time series into variable-length, statistically homogeneous segments. These segments are then grouped into candidate behavioral motifs through high-dimensional, model-based probabilistic clustering. We demonstrate that this approach achieves detailed rodent behavioral mapping using head inertial data. Identified motifs, corroborated by video recordings, include orienting movements, grooming components, locomotion, and olfactory exploration. Higher-order behavioral structures can be accessed by applying a categorical hidden Markov model to the motif sequence. Additionally, our pipeline detects both overt and subtle motor changes in a mouse model of Parkinson’s disease and levodopa-induced dyskinesia, highlighting its value for behavioral phenotyping. This methodology offers the possibility of conducting high-resolution, observer-unbiased behavioral analysis at minimal computational cost from easily scalable and environmentally unconstrained recordings.

## Introduction

In animal studies, investigations into brain-behavior relationships have typically relied on constrained experimental environments with specialized instruments designed to monitor predefined actions, such as lever presses or nose pokes [1]. Advancing toward more flexible and ethologically relevant experimental designs requires innovative methods for continuously tracking animal activities over time and objectively parsing the underlying action sequences [2, 3]. Progress in this field has been largely driven by computer vision techniques [4, 5], particularly through the use of convolutional neural networks to detect user-defined anatomical landmarks, or keypoints, in images of behaving animals [6–8]. Applied to multi-view video recordings, keypoint detection can be used to compute three-dimensional pose estimates [9–14], which can then serve as input data for machine learning-based behavioral mapping frameworks [13, 15–18].

However, these approaches come with several limitations that impose significant constraints on how they can be effectively deployed. Keypoint detection is particularly vulnerable to occlusions, which occur when body parts are obscured by environmental elements, other individuals, or when animals adopt contorted postures. Additionally, the estimation of keypoint localization is inherently jittery, reducing the spatio-temporal resolution of this technique and complicating subsequent data analysis [12, 18]. In larger environments, the trade-off between image resolution and field-of-view leads to reduced spatial accuracy unless a complex, self-orienting video tracking system is employed [19]. Enhancing image resolution and increasing frame rate to minimize motion blur can produce sharper measurements but results in larger data volumes. Threedimensional pose estimation presents further challenges, including the need for meticulous camera calibration, a strategy to mitigate the effect of potential frame drops and the management of larger data loads, which are often incompatible with long-term recordings.

Wearable inertial measurement units (IMUs), which directly report the instantaneous three-dimensional angular velocity and acceleration of specific body parts, provide a compelling alternative for monitoring behavior. These sensors produce small data files compatible with longitudinal onboard data logging, accounting for their widespread adoption among field ethologists [20, 21], animal welfare scientists [22], biomechanists [23] and clinicians [24, 25]. Due to their high precision, exquisite temporal resolution, and compact, lightweight form factors, IMUs are also increasingly favored by physiologists for tracking rodent head [26–45], limb [46, 47], and body [48–52] movements. Being environment-agnostic, these devices are ideally suited for transitioning toward more naturalistic experimental settings involving large, complex or crowded setups. Reflecting their growing popularity in the field, IMUs are now routinely integrated into both commercial and open source head-mounted electrophysiological and imaging devices [53–55]. Despite these advantages, inertial data have rarely been used as a main source of information for automating behavioral analysis in experimental animal models [26, 43, 51], in contrast with IMU-based approaches in human activity recognition [56], animal health monitoring [57] and ethological research [58]. Yet, the high sensitivity, data efficiency and versatility of IMUs are well-suited for developing advanced analysis frameworks aimed at unraveling the architecture of naturalistic behavior, offering the enhanced resolution required by computational neuroethology [2] and deep behavioral phenotyping [59].

To unlock the potential of inertial recordings for mapping rodent behavior, we developed a fully unsupervised analysis pipeline called DISSeCT (Decomposing Inertial Sequences by Segmentation and Clustering in Tandem), which parses IMU signals into discrete activity motifs. Our pipeline employs a two-step process: first, transitions between statistically homogeneous segments are detected using an optimal, computationally efficient change-point detection method, taking full advantage of the low noise, and high sampling rate of IMU recordings. The resulting segments are then grouped into candidate behavioral motifs using high-dimensional Gaussian mixture modeling. We ran our pipeline on head inertial recordings obtained from freely moving rodents using both wired and wireless 6-axis digital IMUs, leveraging the fact that these animals rely heavily on head movements to explore their environment [31, 37, 60] and that many of their activities exhibit distinct head inertial signatures [26, 30, 43, 61]. We show that DISSeCT effectively decomposes inertial data into meaningful activity motifs, validated by concurrent video recordings, parsing the full repertoire of orienting movements, isolating rearing and locomotion episodes, distinguishing the main components of self-grooming, and highlighting episodes of olfactory exploration. Using a mouse model of Parkinson’s disease and levodopa- induced dyskinesia, we also illustrate how our pipeline can be used to detect and characterize behavioral anomalies.

## Results

### Pipeline architecture

Inertial measurements of head kinematics were conducted on four Long-Evans rats placed inside a 70 × 70 cm arena. A custom multi-angle video capture system, operating synchronously, was used to independently document the animals’ activity and compute three-dimensional pose estimation (see Methods). The headborne IMU contained a 3-axis accelerometer and gyroscope, providing head-referenced three-dimensional vector data for acceleration (denoted as **a**) and angular velocity (denoted as ω), respectively. An estimation of the gravitational component of acceleration (denoted as **a**_**G**_) was computed using a previously validated method that combines **a** and ω [42] (see Methods), fully accounting for head orientation relative to gravity (hereafter referred to as head tilt). Subtracting **a**_**G**_ from **a** provided an estimate of the non-gravitational component of acceleration (denoted as **a**_**nG**_), representing acceleration generated by the animal’s movements (Fig. 1A, left).

**Figure 1:**
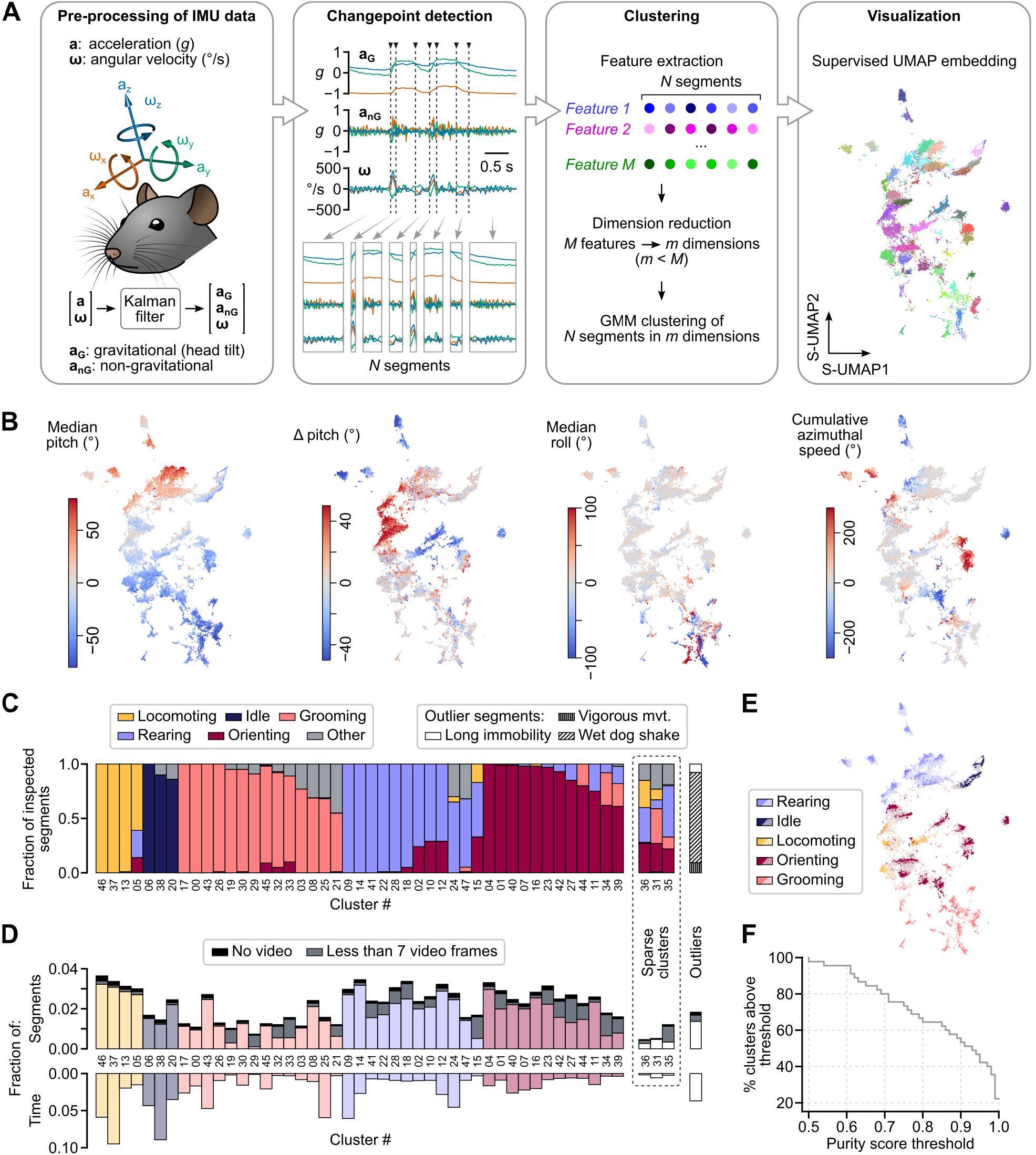
Decomposition of rat behavior using DISSeCT. **A**, Pipeline architecture. **B**, Example supervised UMAP segment embeddings with color-coded feature values. **C**, Stacked bar plot depicting the assignment of main behavioral categories to the top 100 most representative segments within each cluster (see Methods). Clusters are grouped by their dominant category and sorted by decreasing purity score. In the case of outliers (rightmost bar), all segments were examined. **D**, Top: fraction of the total number of segments represented by each cluster. For each cluster, the proportion of segments whose corresponding video snippet was either lacking or comprised less than 7 frames is highlighted in black and grey, respectively. Bottom: fraction of the total recording duration represented by each cluster (all segments included). **E**, Supervised UMAP embedding showing segments color-coded according to their cluster’s dominant main category (dark colors: inspected segments; light colors: other segments). **F**, Fraction of clusters with a purity score above a given threshold, plotted as a function of that threshold.

The initial stage of DISSeCT aims to break down the multivariate time series composed of **a**_**G**_ and ω into non-overlapping segments with stationary statistics, in accordance with the notion that behavior can be conceptualized as a succession of elementary motifs [62, 63]. To achieve this, we opted for kernel-based change-point detection (CPD) using a Gaussian Radial Basis Function kernel [64], implemented through an algorithm that ensures optimal segmentation with a linear computational cost [65, 66] (see Methods). Unlike other techniques that focus solely on detecting shifts in mean values [67] or changes in variances [68], this CPD variant identifies changes in the entire distribution of the multivariate time series [69]. This allows for the detection of a broader range of segment types, effectively generalizing most traditional CPD methods. In our specific use case, this method outperformed a more traditional linear-kernel-based CPD approach in accurately isolating wet dog shake events (Fig. S3 and S5). The algorithm was calibrated to produce a median segment duration of 300 to 400 ms, aligning with the timescale of previously identified mouse behavioral modules [62]. Using this methodology, our dataset, totalling 8 h and 38 min of IMU recordings sampled at 300 Hz, was parsed into 44 322 segments (median duration: 377 ms, average duration: 701 ms, *q*_25_ = 203 ms, *q*_75_ = 820 ms) in about 5 min on a conventional workstation (see Table S3).

The second stage of DISSeCT aims to conduct a cluster analysis of segments, treating each segment as an individual observation. It begins by extracting a catalog of 240 features that characterize segment properties. These features comprise multiple statistics on **a**_**G**_, **a**_**nG**_ and ω, as well as time-frequency attributes of **a**_**nG**_ and ω along with directional information computed from **a**_**G**_ and ω (see Supplementary Methods). One option, as explored in previous studies [13, 15–17, 70], would have been to implement the clustering step by partitioning a two-dimensional representation of the feature space, albeit at the expense of losing potentially valuable information [71]. Instead, our approach involves applying a more conservative dimensionality reduction to maintain a high-dimensional space for clustering while achieving manageable runtimes. To accomplish this, standardized feature data were subjected to principal component analysis, retaining the first 30 principal components (PCs) as determined optimally by cross validation (see Methods). After an outlier removal step, a Gaussian mixture model (GMM) was then fitted to the data in the PC space. Unlike standard clustering algorithms such as K-means [72], GMMs can capture a broader variety of cluster shapes—not just hyperspherical ones—by modeling the underlying data distribution as a mixture of Gaussian components [73]. Moreover, no hypothesis is made regarding the sizes of the clusters such that clusters with heterogeneous weights can be retrieved [74]. Finally, as a statistical model, GMM allows existing statistical tools to be employed. In particular, the Bayesian information criterion (BIC), which comes with theoretical backing, can be used to estimate the number of clusters present in the data [75]. Thus, considering a grid over the number of clusters and computing BIC for each GMM fit [76], we obtained 48 mixture components (Fig. S4B). The corresponding 48 clusters were identified by assigning each segment to the mixture component to which it most likely belonged to. For visualization purposes, we employed a semi-supervised UMAP (S-UMAP) to project segments onto a two-dimensional embedding of the standardized feature space, using cluster identities obtained from the GMM. This embedding was not used for clustering, but rather to improve the interpretability of the feature space by aligning visual boundaries with cluster labels. Compared to a purely unsupervised embedding, S-UMAP more clearly reveals the internal structure of each cluster while preserving meaningful gradients in feature values (Fig. 1B; see Fig. S6 for comparison with unsupervised and shuffled-label embeddings).

To validate the usefulness of CPD, we conducted a posthoc comparison against a segmentation strategy based on fixed-length windows, a common default starting point in several behavioral analysis pipelines [63, 77]. To this end, we divided IMU recordings into non-overlapping windows of 210 samples (700 ms), matching the average segment duration obtained with CPD and thus preserving the total number of segments. We then applied the same downstream processing pipeline combining feature extraction, dimensionality reduction (PCA), and GMM clustering, retaining for these steps the hyperparameters obtained for varying-length segments. Specifically, we assessed how well GMMs, fitted to a PC representation of segments produced by each method, explained the underlying data. Fixed-length segmentation initially yielded higher log-likelihood values (Fig. S2E), likely due to the subdivision of immobility periods into numerous low-variability windows that are easily modeled. This effect artificially inflates GMM performance compared to the CPD approach, which isolates immobility bouts as individual segments. After ablating periods of immobility prior to modeling (see Methods), the CPD-based strategy significantly outperformed the fixed-length approach (Fig. S2D,E), highlighting CPD’s effectiveness in capturing meaningful structure for subsequent clustering.

In summary, DISSeCT provides a unified framework for pre-processing and parsing IMU data. It essentially relies on a fully unsupervised two-step process, wherein time series segmentation is used to isolate elementary kinematic motifs that are subsequently clustered in a high-dimensional space.

### Primary behavioral categories are parsed into distinct families of clusters

To assess the behavioral significance of IMU data decomposition using DISSeCT, we proceeded as follows: for each cluster, an observer was tasked with labeling the top one hundred segments with the highest posterior probabilities of belonging to it, and a *purity score* was computed to highlight the prevalence of the most frequent label (see Methods and Table S6). With the exception of three sparse clusters, which represented highly scattered and underpopulated mixture components (see Methods and Fig. S7), all clusters exhibited a purity score of at least 0.5 (with two-thirds of them displaying scores above 0.8, Fig. 1F) and could be clearly associated with a predominant behavioral category (orienting, rearing, grooming, locomoting or idle; see Fig. 1C,D) occupying a distinct sub-region of the supervised UMAP embedding (Fig. 1E). Purity scores tended to decrease when labeling was performed on randomly-selected segments (Fig. S12C), suggesting increased behavioral ambiguity in regions farther from mixture centroids and closer to cluster boundaries. A significant fraction of clusters (19 out of the 45 non sparse clusters), however, displayed similar purity scores irrespective of the segment selection method (Fig. S12E). This analysis shows that the decomposition of IMU data obtained with our pipeline largely aligns with the major classes of rodent activities. The fact that each behavioral category was subdivided into several clusters, with three of them (orienting, rearing and grooming) comprising no fewer than 12 clusters each (Fig. 1C), indicated a higher level of granularity where each category is further broken down into multiple components. This prompted us to conduct a comprehensive examination of the GMM output in order to assess the level of detail achieved with DISSeCT.

### Identification of clusters mapping the repertoire of rapid changes in head orientation

Considering that our primary data included direct measurements of three-dimensional head angular velocity, we anticipated that DISSeCT would be able to discern motor patterns associated with rapid changes in head orientation. A preliminary examination of the output of the CPD step indeed suggested that such movements were isolated as individual segments (see Movie S1). Among the features extracted from all segments, we focused on net angular speed, calculated by dividing the net head angular displacement (from segment start to finish) by the segment’s duration. The distribution of per-cluster average net angular speed values exhibited a bimodal pattern, with a cutoff of about 150 ^°^/s (Fig. 2A). The upper half of this distribution contained 18 clusters representing brief (179 ± 45 ms) head rotations associated with the “orienting”, “rearing” and “grooming” categories (Fig. 2A,B) and accounting for 37% of all segments (11% of total recording duration). With the exception of the rearing termination phase, all these clusters corresponded to rotations of 30–50^°^ (mean: 42.8 ± 8.8^°^, Fig. 2B), regardless of the net angular speed (*r* = 0.033, *p* = 0.90). This can be explained by the fact that the average segment duration decreased as a function of the average net angular speed (*r* = −0.86, *p* = 1.6e−5), suggesting that rapid head movements might be tuned to produce relatively invariant head angular displacement.

**Figure 2:**
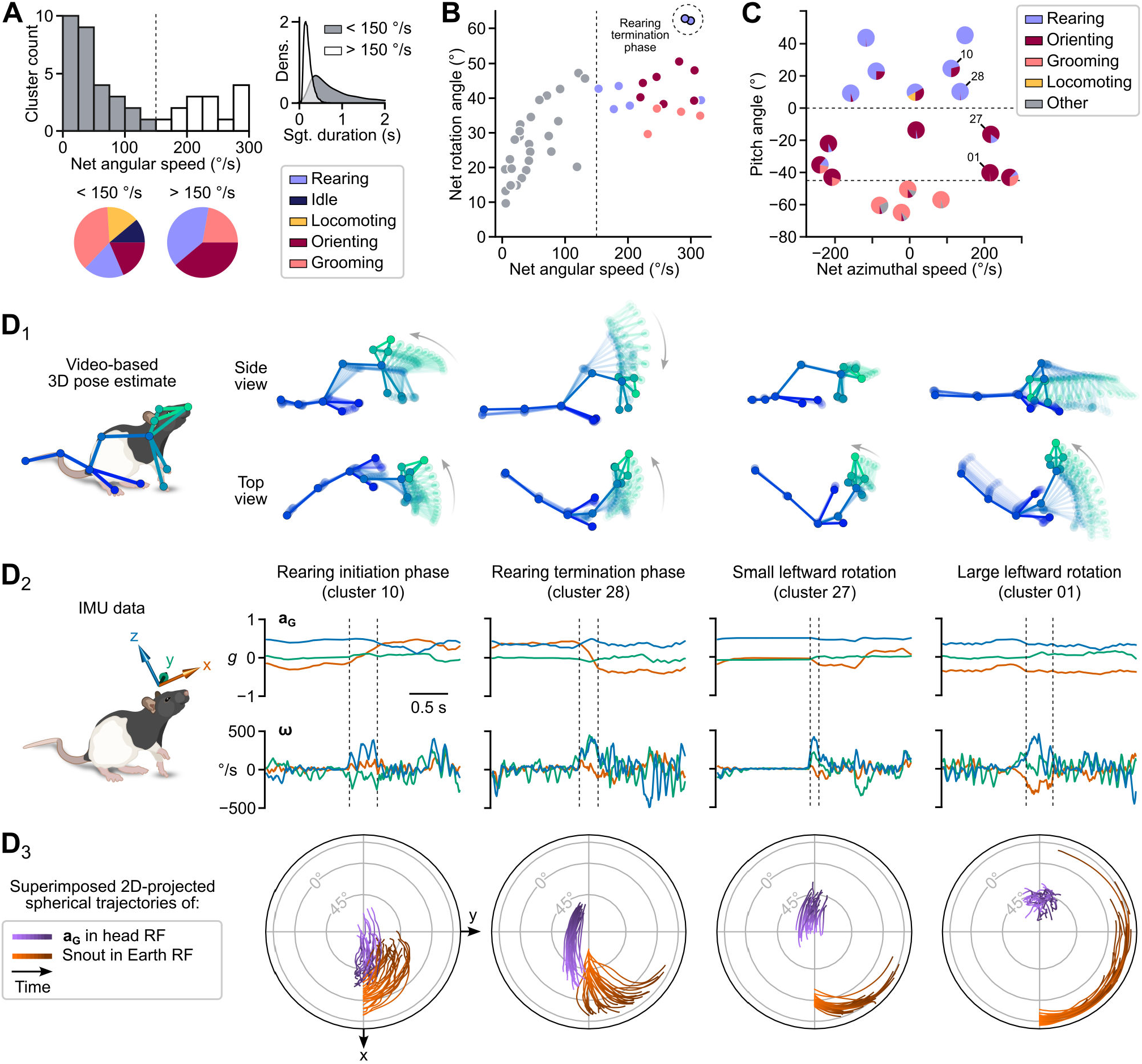
Diversity and fine dynamics of rapid changes in head orientation. **A**, Distribution of net angular speed values (median of per-segment average values) for all clusters. The dashed vertical line at 150 ^°^ /s corresponds to the threshold for selecting clusters representing rapid head rotations (highlighted in white in the histogram). Pie charts illustrate the proportion of cluster-associated behavioral categories (one unique category per cluster) for each side of the distribution. The upper right inset shows a density estimate of segment duration for each side of the distribution. **B**, Net rotation angle plotted against net angular speed (median of per-segment average values) for each cluster. Clusters are colored according to their associated behavioral category (using the same color code as in A and Fig. 1C). **C**, Head pitch angle (see Methods) plotted as a function of net azimuthal speed (median of per-segment average values) for each cluster. Each cluster is displayed as a pie chart showing the behavioral categories attributed to the top 100 segments of the cluster (cf. Fig. 1C). **D**_**1–3**_, Detailed examination of four example clusters (identified in C). Each column corresponds to one cluster. D_1_: Skeletons representing the animal’s 3D pose dynamics during one example segment. Zero transparency corresponds to the last video frame of the segment. D_2_: Gravitational acceleration and angular velocity signals (**a**_**G**_ and ω, respectively) for the same segment. Dashed lines represent segment boundaries. D_3_: Joint visualization of the trajectories of **a**_**G**_ in the head reference frame (purple) and of the animal’s heading in the Earth reference frame (orange) for the top 30 segments of each cluster (see Supplementary Methods; RF: reference frame).

Behavioral categories could be distinguished based on the head pitch angle and net azimuthal speed (Fig. 2C): grooming-associated rotations, corresponding to single strokes or brief head repositioning between grooming bouts (Movie S1), occurred with the head pitched downward (< −45^°^) and showed limited change in azimuth; rearing-associated rotations, mainly corresponding to rearing initiation or termination, took place with the head pitched upward (> 0^°^); “orienting” movements predominantly fell within an intermediate range of pitch angles, accompanied by either minimal or significant changes in azimuth.

These observations show that our pipeline offers an effective method for isolating and sorting rapid three-dimensional changes in head orientation. A visually informative overview depicting the corresponding repertoire of such movements was obtained using custom two-dimensional projected maps showing the segment-wise spherical trajectories of **a**_**G**_ in the head reference frame and of the animal’s heading in an absolute (external) reference frame (see Supplementary Methods and Movie S2). These maps are shown together with the associated 3D poses and IMU signals in Fig. 2D.

We noticed that some clusters contained segments belonging to different categories (Fig. 2C), interpreting this as a consequence of distinct behavioral motifs sometimes generating similar head movements. For example, rearing initiation and upward head rotation may share common head kinematic features, differing primarily in whether the forepaws are leaving the ground or not. This point is further investigated in a later section where we show that such ambiguities can be resolved by taking into account the sequences of behavioral motifs.

### Locomotion splits into different regimes, including a mobile olfactory exploration module

As shown in Fig. 1C,D, the “locomotion” and “idle” categories, which accounted for 18 % of all segments and 36 % of the total recording duration, contained fewer clusters compared to other behavioral categories, suggesting a higher degree of kinematic homogeneity. When considering actual animal displacement, three of the “locomotion” clusters emerged as those associated with the highest speed (Fig. 3A), representing 68 % of the time when the animal’s speed exceeded 15 cm/s. Two of the “idle” clusters emerged as those with the maximal fraction of time spent immobile (defined using a head angular speed threshold; see Methods; Fig. 3A). One of these (cluster 6) corresponded to true periods of immobility, while the other (cluster 38) contained longer duration segments during which the animal was nearly immobile but exhibited small head movements (Fig. 3A,B).

**Figure 3:**
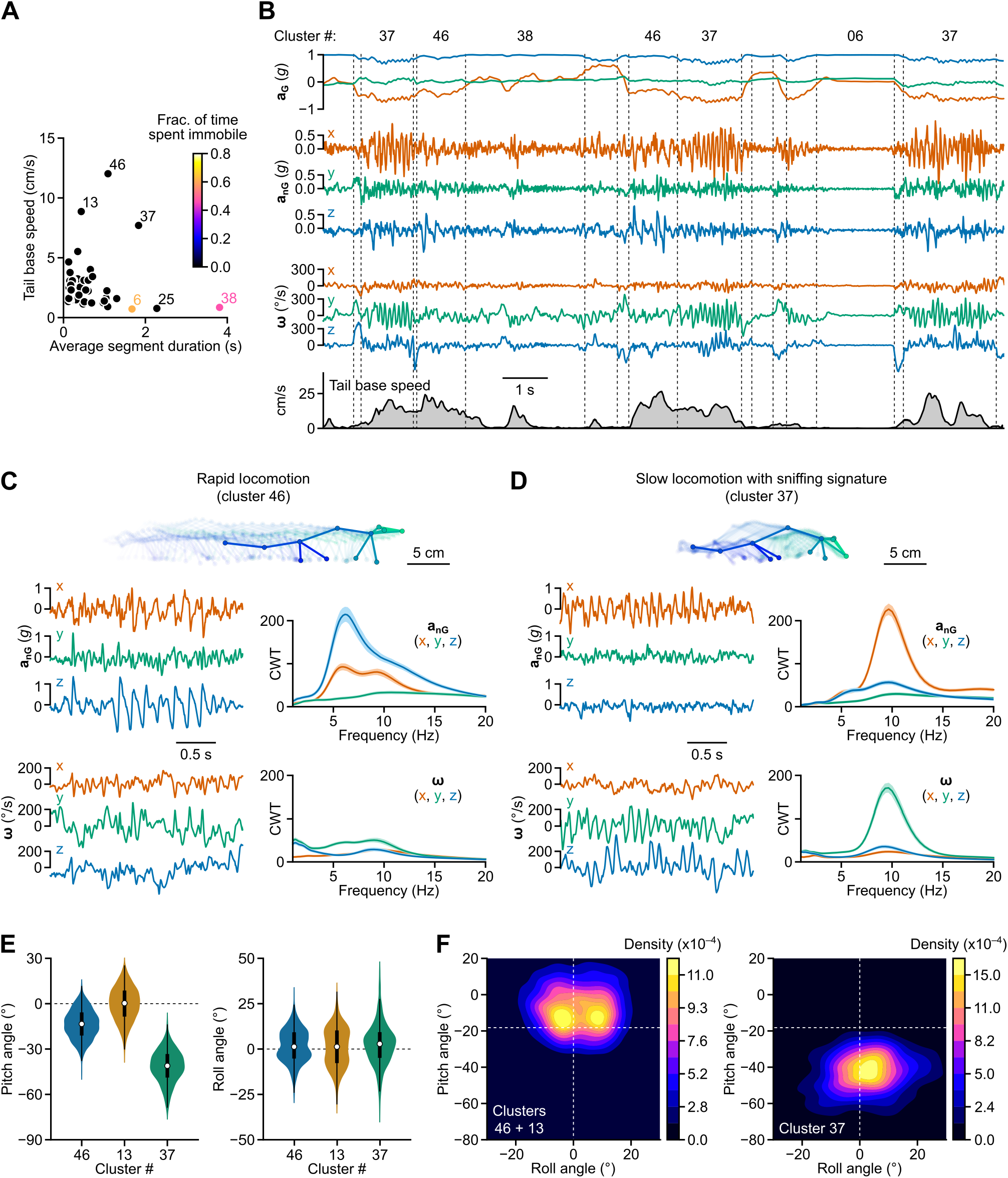
Identification of two locomotor regimes. **A**, Tail base speed (median of per-segment average values) plotted against average segment duration for each cluster. The color scale represents the median fraction of time spent immobile, computed for each segment based on head angular speed (see Methods). Cluster numbers are indicated for three “locomotion” (13, 37 and 46) and two “idle” clusters (6 and 38). **B**, Upper panels: Example traces of head gravitational and non-gravitational acceleration and angular velocity (**a**_**G**_, **a**_**nG**_ and ω, respectively). Lower panel: Concurrent horizontal tail base speed computed from 3D pose estimates. Dashed vertical lines indicate segment boundaries. Corresponding cluster numbers are shown above the traces. **C**, Top: Skeletons representing the animal’s 3D pose dynamics during 30 video frames (1 s) extracted from the segment with the highest posterior probability of belonging to cluster 46 (associated with rapid locomotion). Zero transparency corresponds to the last frame. Left: Example traces of head non-gravitational acceleration and angular velocity (**a**_**nG**_ and ω, respectively) for the same segment. Right: Continuous wavelet transform (CWT) coefficients for all three axes of **a**_**nG**_ and ω. Solid lines represent the average of per-segment median log-magnitude values (shared area: 95% confidence interval). **D**, Similar to panel **C**, but for cluster 37 (corresponding to mobile olfactory exploration). **E**, Distribution of per-segment average head pitch and roll angles for the three “locomotion” clusters identified in **A. F**, 2D histograms showing the distribution of per-segment average head pitch and roll angles for clusters associated with rapid locomotion (left) and mobile olfactory exploration (right). Dashed lines indicate the resting pitch and roll angles, computed as the median of per-segment average pitch and roll angles for all segments.

To understand what caused the clustering algorithm to isolate locomotor activity, we examined the head kinematic features associated with the three “locomotion” clusters highlighted in Fig. 3A. Two clusters (46 and 13), corresponding to second-scale fast locomotion bouts, were notably characterized by sawtooth-like patterns in the *z* component of non-gravitational acceleration (Fig. 3B,C) synced with each step cycle, as previously described from head acceleration measurements in running rats [28]. This signature likely reflects a bobbing head motion associated with the successive cycles of storage and release of kinetic and potential energy in the animal’s skeletal muscles [78]. Continuous wavelet transform revealed that this oscillatory regime took place in the 5–7 Hz band, consistent with the known step cycle duration during trotting in rats [79].

The third cluster (37) corresponded to a strikingly different locomotor regime in which most of the power was concentrated around 10 Hz in the *x* and *y* axes of non-gravitational acceleration and angular velocity, respectively (Fig. 3D). This signature, reflecting coordinated fore-aft motion and pitch rotation, has already been described as linked with sniffing in rodents [31, 80]. Compared to the other two “locomotion” clusters, this locomotor regime was also characterized by a larger downward head pitch angle of about 40^°^ (Fig. 3E). Video inspection clearly showed that this third locomotion cluster corresponded to episodes of olfactory exploration, during which animals slowly moved forward maintaining their snout close to the floor (Movie S3).

Head pitch was not the only feature distinguishing mobile olfactory exploration from rapid locomotion. While the distribution of head roll was unimodal in the first case, it appeared to be bimodal in the second (Fig. 3F), peaking at 5–10^°^ toward the left and right sides. This phenomenon did not result from variability in IMU placement on the head, as segments from the same animal, and even from the same session, showed both orientations. During rapid locomotion, per-segment average roll angles showed a weak correlation with net azimuthal speed (*r* = 0.15, *p* < 1e−4). This suggests that head roll tends to occur when the animal follows a curved path, with the head tilting towards the concave side of the trajectory.

### The vigor of olfactory exploration can be tracked by analyzing its spectral signature

Having observed a head kinematic signature linked with sniffing during slow locomotion, we wondered if a similar pattern could be detected in other behavioral contexts. Visual inspection of segments and their associated video snippets indeed suggested that ≃ 10 Hz rhythmicity in the fore-aft non-gravitational acceleration and pitch angular velocity was a prevalent feature of head IMU recordings also manifesting in other clusters. To identify these clusters, we devised a “sniffing score” reflecting the magnitude of this spectral signature based on a per-segment ratio of continuous wavelet transform coefficients in specific frequency bands (see Fig. 4A and Methods).

**Figure 4:**
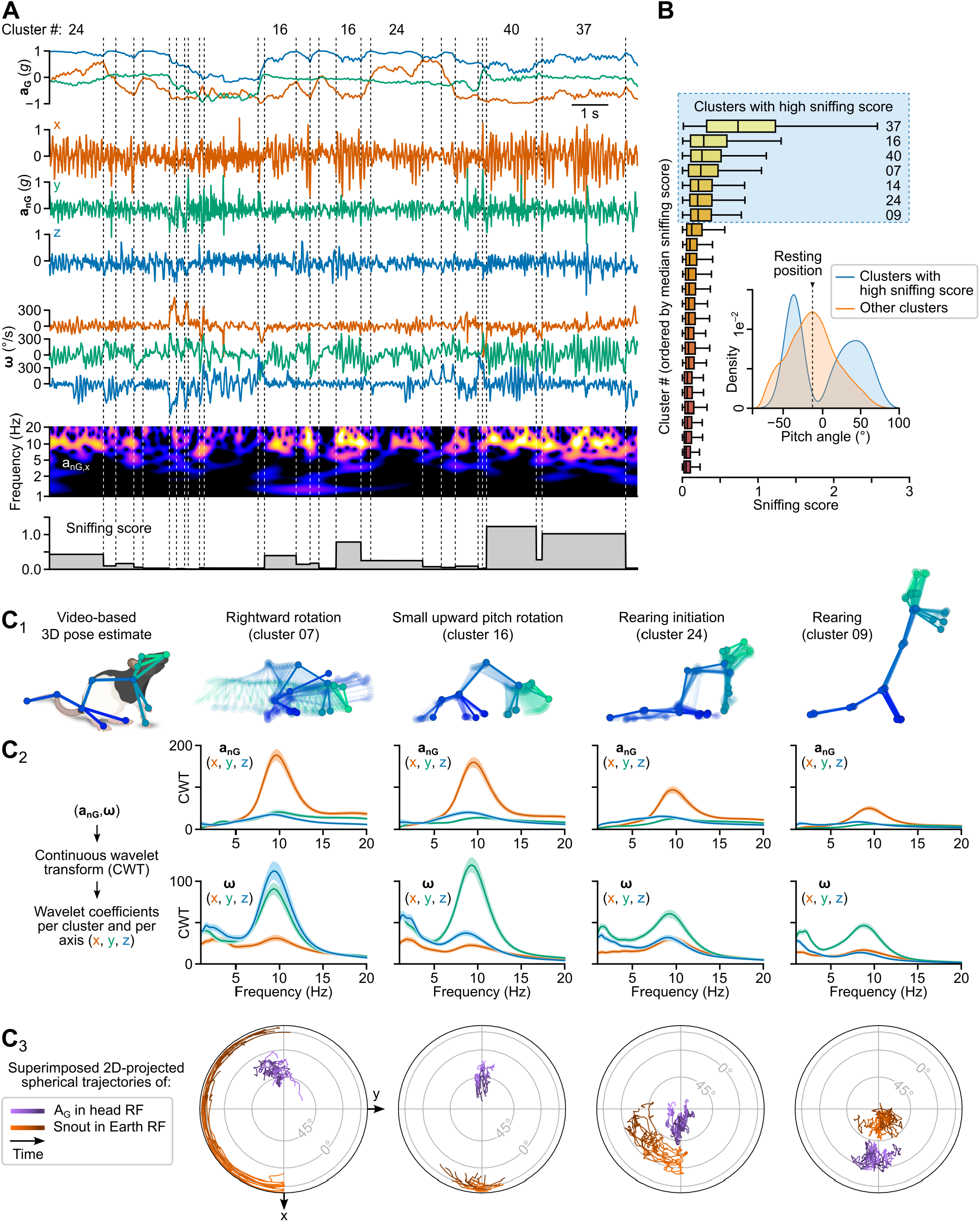
Identification of a sniffing signature. **A**, Upper panels: example traces of head gravitational acceleration (**a**_**G**_), non-gravitational acceleration (**a**_**nG**_) and angular velocity (ω). Lower panels: log-magnitude of continuous wavelet transform coefficients computed for fore-aft non-gravitational acceleration (*x*-axis component of **a**_**G**_), and per-segment sniffing score values. Dashed vertical lines indicate segment boundaries. Corresponding cluster numbers are indicated above the traces. **B**, Box plot of sniffing score values shown for 24 clusters. Clusters are sorted by decreasing median sniffing score. Inset: Distribution of per-segment median head pitch angles computed for the seven clusters with highest sniffing scores (blue curve) and for all other clusters (orange curve). **C**_**1–3**_, Detailed examination of four example clusters with high sniffing scores. Each column corresponds to one cluster. C_1_: Skeletons representing the animal’s 3D pose dynamics during one example segment. Zero transparency corresponds to the last video frame of the segment. C_2_: Log-magnitude of continuous wavelet transform (CWT) coefficients (average of per-segment median values) for all three axes of **a**_**nG**_ and ω, shown for each example cluster (shaded area: 95% confidence interval). C_3_: Joint visualization of the trajectories of **a**_**G**_ in the head reference frame (purple) and of the animal’s heading (orange) for the top 10 segments of each cluster (see Supplementary Methods; RF: reference frame).

In addition to cluster 37, associated with the slow locomotor regime identified in Fig. 3D, this approach revealed six clusters exhibiting a high sniffing score (Fig. 4B) and associated with the “orienting” and “rearing” categories. The strongest scores were observed for three “orienting” clusters not part of the rapid head movements identified in Fig. 2B, corresponding to small upward pitch movements or rightward or leftward head rotations with the head pitched down (Fig. 4C), and displaying net angular speed values in the 35–60 ^°^/s range. The two clusters with lower scores corresponded to rearing initiation and high rear with a vertically stretched body (Fig. 4C). These observations, along with the bimodal distribution of average head pitch angles associated with these clusters (Fig. 4C), suggest that animals preferentially engaged in robust sniffing either when investigating the floor or during rearing.

The average peak frequencies observed for these eight clusters with high sniffing score (9.1 ± 2.1 Hz and 8.2 ± 1.7 Hz for fore-aft non-gravitational acceleration and pitch angular velocity, respectively) were in the upper range of sniffing rates reported for rats during odor discrimination in stationary or head fixed conditions (6–9 Hz) [31, 81], and were consistent with those observed in freely-moving animals tracking an odor trail (8–12 Hz) [82]. Overall, these results indicate that head IMU signals can be leveraged to obtain a detailed and quantitative readout of olfactory exploration, using a simple metric that can be applied independently of our pipeline.

### The main types of grooming activities distribute into separate clusters with distinctive spectral signatures

Self-grooming is a major component of the rodent behavioral repertoire, encompassing a variety of actions often performed sequentially [83]. Monitoring grooming activities through video recordings is however challenging due to the animal’s contorted posture (e.g. during body licking) or the rapid movements involved (e.g. during scratching). To assess DISSeCT’s ability to parse this family of actions, we conducted a detailed examination of the 14 clusters assigned to the “grooming” category during segment inspection (Fig. 1C), which accounted for 19 % of all segments and 23 % of the total recording duration.

These clusters were characterized by a specific range of head tilts, with the less ambiguous ones (corresponding to body licking) exhibiting a unique combination of head pitch and roll angles (see cluster 17 in Fig. 5A). Unexpectedly, their most distinctive feature was a clear deviation from a linear relationship observed for all other clusters between the correlation of *x* and *z*-axes gyroscope measurements and the head pitch angle (*r* = 0.95, *p* = 2.2e−16; Fig. 5A). In other words, the coupling between roll and yaw head rotations during grooming appears to depart from a general biomechanical rule linking it with head pitch in all other conditions.

**Figure 5:**
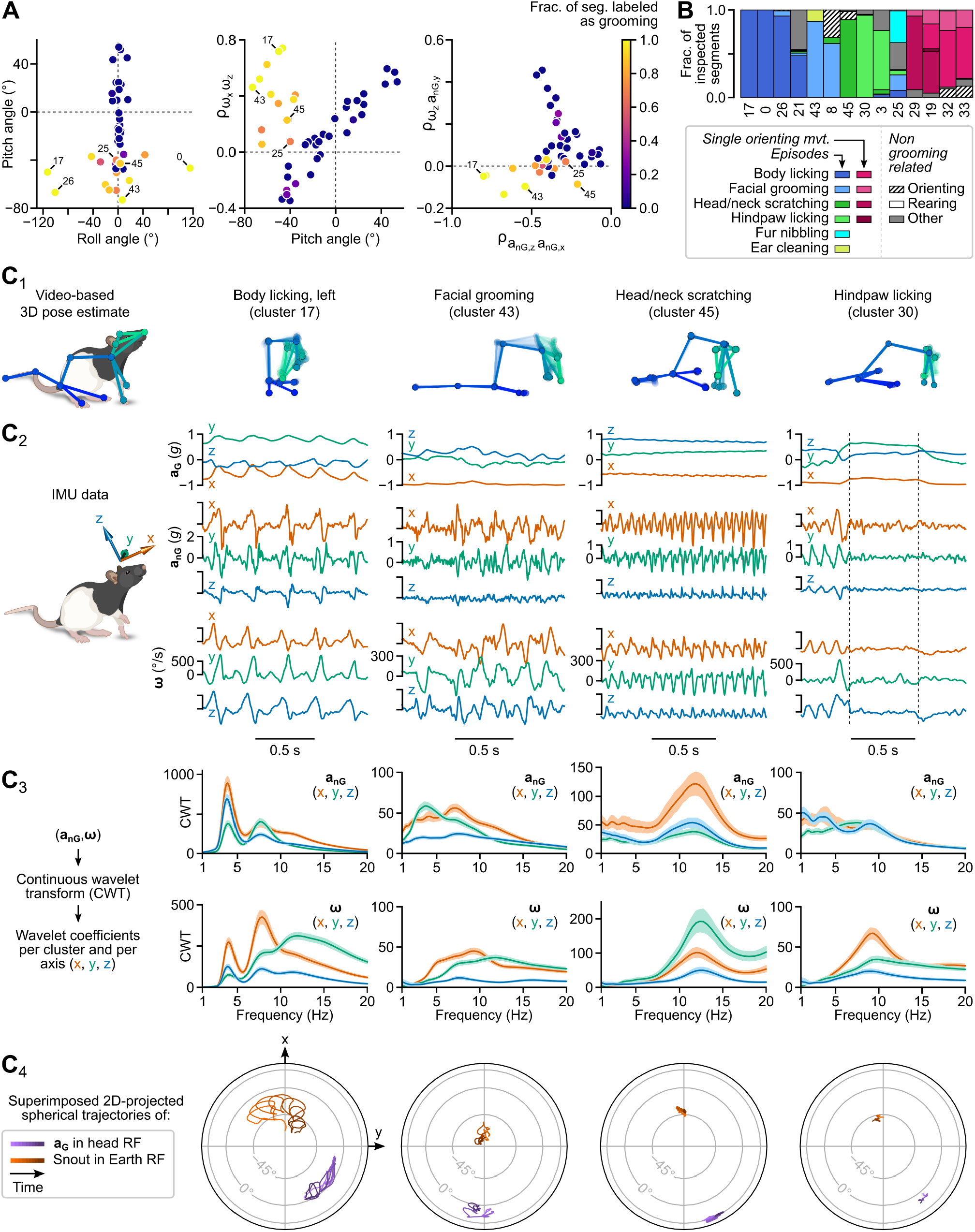
Parsing of grooming components. **A**, Distinctive features of grooming-related clusters. Left: head pitch vs. roll angles. Middle: correlation between angular velocities about the *x* and *z* axes (ω_*x*_ and ω_*z*_, respectively) plotted against head pitch angle. Right: correlation between angular velocity about the *z* axis (ω_*z*_) and medio-lateral non-gravitational acceleration (*a*_*nG,y*_) plotted against the correlation between the head-vertical and fore-aft components of non-gravitational acceleration (*a*_*nG,z*_ and *a*_*nG,x*_, respectively). These panels show per-cluster median values of per-segment average values. The color scale indicates the fraction of segments identified as belonging to a grooming sequence during inspection of the top 100 segments of each cluster (see Method). **B**, Stacked bar plot depicting the assignment of labels to inspected segments within each grooming-related cluster. **C**_**1–3**_, Detailed examination of four example clusters. Each column corresponds to one cluster. C_1_: Skeletons representing the animal’s 3D pose dynamics during one example segment. Zero transparency corresponds to the last video frame of the segment. C_2_: Gravitational and non-gravitational acceleration (**a**_**G**_ and **a**_**nG**_, respectively), and angular velocity (ω) for the same segment. Average segment-wise median log-magnitude of continuous wavelet transform (CWT) coefficients for all three axes of **a**_**nG**_ and ω (shared area: 95 % confidence interval).

To assess whether “grooming” clusters corresponded to ethologically distinct actions, we relied on the labels assigned during the inspection phase (Fig. 5B; see Methods). The less ambiguous clusters (where more than 85 % of inspected segments received the same label) corresponded to body licking (clusters 0, 17 and 26), facial grooming (cluster 43), head and neck scratching (cluster 45) and hindpaw licking (cluster 30), each exhibiting a characteristic spectral signature (Fig. 5C). Another group of clusters had less homogeneous labels, with short bouts of facial grooming mixed with non-grooming related head orienting movements (cluster 8), or individual body licking strokes mixed with brief head movements occurring when the animal investigated the corners of the arena with a tilted head posture (cluster 21). The non grooming-related segments included in these clusters likely correspond to head motion patterns whose kinematics resemble those encountered during grooming. Kinematic ambiguities may also explain why ethologically distinct grooming actions were mixed in the same clusters (see clusters 43 and 25, Fig. 5B). Finally, a last group of clusters (clusters 19, 29, 32 and 33) contained various head orienting movements executed in the context of grooming episodes (at episode onset or offset or between adjacent grooming bouts).

These results show that our pipeline successfully isolated the main components of grooming as a family of clusters with distinctive postural, kinematic and spectral signatures. They also point out kinematic ambiguities which could be taken into account to design strategies aiming at achieving an even greater separation of grooming actions.

### Behavioral patterns can be unveiled from cluster sequences

As detailed in the previous sections, DISSeCT effectively parsed head IMU signals into various behaviorally relevant components. To assess how these components assemble over time and potentially form sequential patterns, we fitted a simple hidden Markov model (HMM) with categorical emissions to the cluster sequence (i.e. where categories correspond to clusters). Hyperparameter optimization resulted in 16 states, each associated with a vector of emission probabilities for the different clusters (see Methods; Fig. 6A). We observed that states often appropriately gathered clusters corresponding to the same movement executed in opposite leftward vs. rightward directions (Fig. S8). Additionally, the highest emission probabilities for a given state most often corresponded to clusters belonging to the same behavioral category, allowing each state to be linked to a specific category (Fig. 6A).

**Figure 6:**
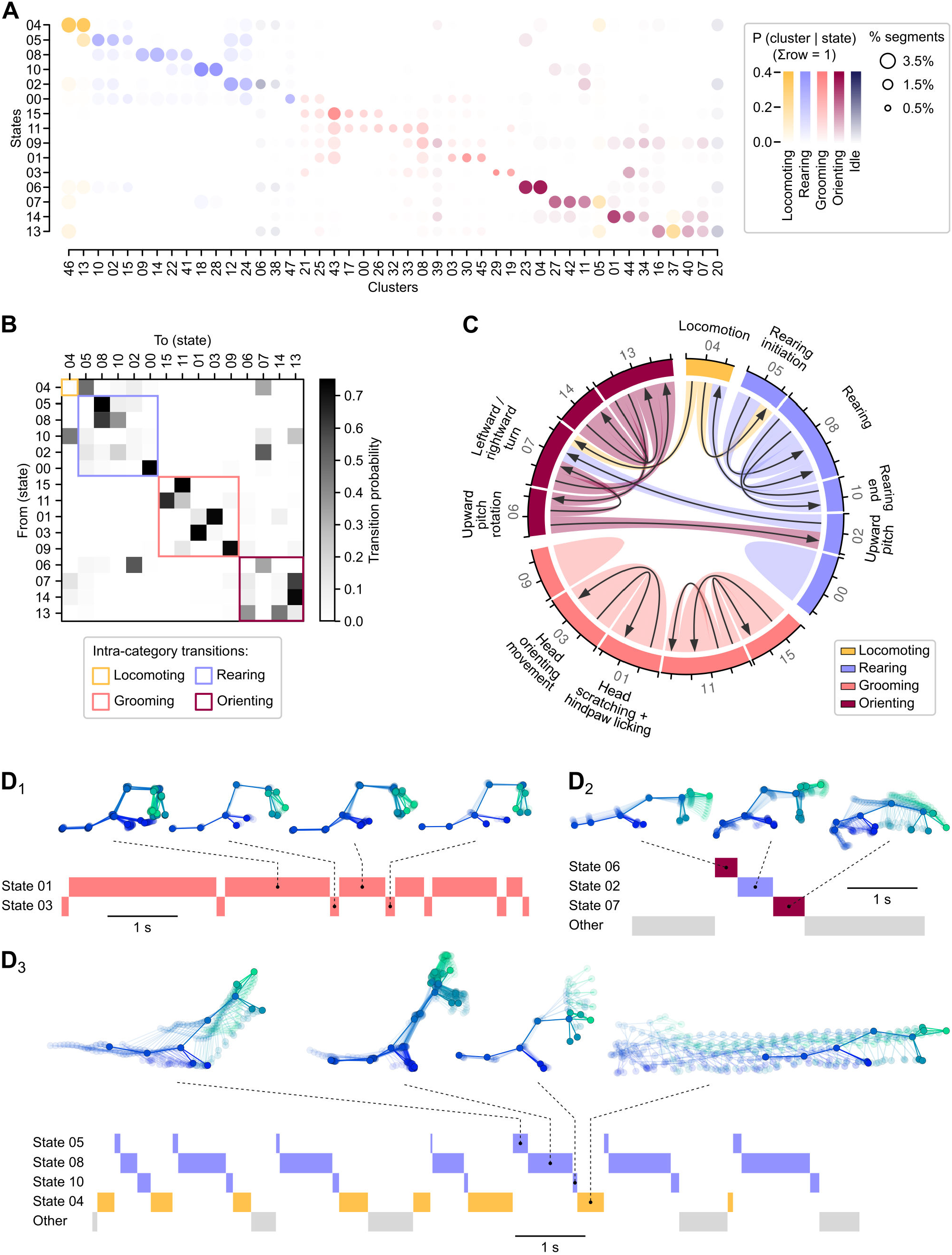
Identification of behavioral sequences using a post-hoc categorical hidden Markov model. **A**, Cluster emission probabilities (columns) for each hidden state (rows). The dots in each column are colored according to the cluster’s associated main behavioral category as defined in Fig. 1C, with the opacity representing probability values and dot area representing how much the cluster contributed to the total number of segments. Note that state 12, which contains outlier segments, and sparse clusters (see Methods and Fig. S7) are not represented in this plot. **B**, Hidden state transition matrix. States are ordered according to their dominant main behavioral category. Colored squares highlight intra-category transitions. **C**, Chord plot representing transition probabilities between states. Arrow width represents transition probability (tick spacing: 0.5). States are colored according to their dominant main behavioral category. Only transitions probabilities greater than 0.25 are represented. **D**_**1–3**_, Representative ethograms for three types of sequences. D_1_: Alternation between head scratching or hindpaw licking episodes (state 1) and brief head orienting movements (state 3). D_2_: Single episode during which the animal is lifting its head up before executing a turn. D_3_: Alternation between rearing episodes (states 5, 8 and 10) and locomotion bouts (state 4). For each ethogram, skeletons representing the animal’s 3D pose dynamics are shown for example segments. Zero transparency corresponds to the last video frame of the segment.

The state transition matrix revealed that the most likely transitions occurred within the “rearing”, “grooming” and “orienting” categories (Fig. 6B), indicating that these behaviors tended to manifest as multi-state sequences rather than isolated events. States associated with the “rearing” and “grooming” categories, in particular, almost never transitioned to one another, while the “locomoting”, “rearing” and “orienting” categories did exhibit specific temporal relationships. These observations were further confirmed by examining the cluster transition matrix (Fig. S9).

Characteristic intra-category sequences included back-and-forth switching between states 1 and 3, representing an alternation between head scratching or hindpaw licking episodes and brief head orienting movements (Fig. 6C,D_1_), and a 3-state chain (states 5, 8 and 10) representing rearing episodes (Fig. 6C). Prominent inter-category sequences included an alternation between rearing episodes and locomotion bouts (Fig. 6C,D_2_, Fig. S10A), and a 3-state sequence (states 6, 2 and 7) representing episodes during which the animal lifted its head up while keeping its front paws on or close to the ground (Fig. 6C,D_3_, Fig. S10B).

One key property of HMMs is that the probability of being in a given state for a particular observation changes over time depending of the local context. As a result, segments assigned to a given cluster by the GMM but occurring in different behavioral contexts might be associated with different states, potentially reflecting different behavioral motifs. To examine this, we plotted posterior probabilities for each cluster (Fig. S11A) and found clusters linked to multiple states. A detailed analysis of two dual-state clusters revealed that they could indeed be split into kinematically similar but ethologically distinct head movements based on the posterior probability of each segment. Cluster 12 was parsed into rearing segments (linked with state 8) and segments during which the head was pitched-up but the front paws remained on or near the ground (linked with state 2), as shown in Fig. S11B. Cluster 19 was parsed into brief orienting movements executed between head scratching and hindpaw licking bouts (linked with state 3) and single strokes during facial grooming or body licking (linked with state 11), as shown in Fig. S11C.

These observations demonstrate that a simple categorical HMM can be effectively used as a curation step after DISSeCT to merge ethologically-similar clusters and resolve certain kinematic ambiguities by splitting clusters containing similar head movements performed in different contexts.

### Comparison with video-based behavioral analysis

In our rat dataset, IMU recordings were acquired concurrently with multi-view video recordings (see Methods), enabling direct comparison of our pipeline with 3D pose-based behavioral mapping tools. To analyze 3D pose data, we used Keypoint-MoSeq (KpMS), an unsupervised pipeline that applies a hierarchical graphical model to jointly perform pose estimate refinement, segmentation and clustering, and which has been explicitly validated for 3D pose data [18]. KpMS parses a low-dimensional representation of keypoint trajectories into behavioral syllables, each composed of multiple instances. Functionally, these syllables and instances are equivalent to the clusters and segments identified by DISSeCT.

KpMS was parameterized to match the median duration of syllable instances to that of DISSeCT segments and previous studies [18, 62] (see Methods). Under these conditions, it produced a greater proportion of short (< 0.2 s) and intermediate (0.5–2 s) instances (Fig. S12A) and identified fewer syllables (Fig. S12B). In-depth analysis was conducted on syllables accounting for more than 5 % of the total recording duration. Their behavioral content was evaluated via visual inspection, following the same protocol used for DISSeCT clusters (see Methods). A clear behavioral category could be assigned to nearly all KpMS syllables. As expected, purity scores decreased following randomization, though not uniformly across clusters and syllables (Fig. S12C–E), as well as across behavioral categories (Fig. S12G). Syllable-wise purity scores were less affected than cluster-wise scores (Fig. S12F). When purity scores were computed at the level of the main behavioral categories, this difference was more nuanced, with DISSeCT achieving comparable of higher scores than KpMS for the “idle”, “grooming” and “orienting” categories (Fig. S12G). DISSeCT subdivided the “idle”, “grooming”, “rearing”, and “orienting” categories into more clusters, whereas KpMS identified a greater number of “locomotion” syllables. This difference likely reflects the complementarity of both methods: DISSeCT focuses on fine-grained head kinematics, while KpMS—relying on whole-body pose—distinguishes among multiple locomotion bouts based on trajectory information (see Movies S4 and S5).

To quantify the alignment between the two approaches, we computed contingency matrices (S13; see Methods). DISSeCT and KpMS showed broad agreement on the “idle”, “grooming”, and “rearing” categories, but less consensus on “locomoting” and “orienting” (S13B,D). This may be partly explained by the fact that some of DISSeCT’s “orienting” segments are embedded within KpMS’s curved locomotion bouts, as well as by the inherent ambiguity—apparent to human observers—of short video snippets that combine orienting movements with locomotor activity.

Importantly, several behavioral motifs captured by DISSeCT were not identified by KpMS. For example, while KpMS successfully differentiated body licking from facial grooming, it failed to isolate scratching and hindpaw licking as distinct syllables (compare Figs. S13G and 5B). KpMS also did not isolate wet dog shake events, which composed the vast majority of DISSeCT’s outlier segments (1C and Fig. S5). Part of these discrepancies likely stem from the higher temporal resolution required to resolve such rapid behaviors, compounded here by the tenfold difference in sampling rates between IMU and video data (300 Hz vs. 30 Hz). Differences between the outputs of DISSeCT and KpMS likely stem from both the nature of their input data and their underlying mathematical frameworks: DISSeCT yields a finer decomposition of behaviors with a marked head kinematic signature, whereas KpMS offers a coarser clustering—except for locomotion—based on whole-body dynamics. Quantitatively, the Adjusted Rand Index comparing the two clusterings was 0.19 (excluding DISSeCT’s outlier segments), indicating limited structural overlap.

The pipelines also differ substantially in computational demands. Despite the higher IMU sampling rate, DISSeCT—which is CPU-based—ran five times faster than the GPU-based version of KpMS (see Tables S4 and S5), not including the additional cost of video-based 3D pose estimation.

### Unsupervised analysis of behavioral alterations in a mouse model of Parkinson’s disease and levodopa-induced dyskinesia

One advantageous feature of GMMs is their probabilistic nature, which makes them suitable for estimating how well a new dataset conforms to a previously-established model and thus for detecting anomalies. To test the applicability of DISSeCT for quantifying behavioral changes, we used a classical toxin-induced mouse model of Parkinson’s disease (PD). 6-OHDA lesioned mice (PD mice, *n* = 12) were obtained through unilateral nigrostriatal dopaminergic denervation using 6-OHDA injection (see Methods) while a control group received saline injection (sham mice, *n* = 4). PD mice exhibited decreased locomotion, impaired movement initiation and increased ipsiversive rotations relative to the lesion side, as previously described [84]. PD mice were then treated with daily injections of levodopa (L-DOPA), the first-line drug for the management of PD motor symptoms, along with sham animals. L-DOPA induced abnormal involuntary movements (AIMs) in PD mice, mimicking the common complication of this treatment in PD patients, known as levodopa-induced dyskinesia (LID). These AIMs, which affect the musculature on the side contralateral to the lesion [84], were accompanied by contraversive rotations.

Weekly IMU recordings were performed on PD and sham mice before and during L-DOPA treatment (Fig. 7A). DISSeCT was then run on sham data to generate a GMM representing the behavior of healthy animals (sham-GMM, see Methods). A joint S-UMAP embedding, computed across all animals in all conditions using cluster labels from the sham-GMM, revealed a similar global structure for sham and PD mice, while entirely new regions emerged for PD mice during the LID phase (Fig. 7B). These novel regions were also clearly visible in fully unsupervised embeddings based solely on kinematic features (Fig. S14), confirming the emergence of L-DOPA-specific behavioral patterns that are not present in healthy or untreated PD mice.

**Figure 7:**
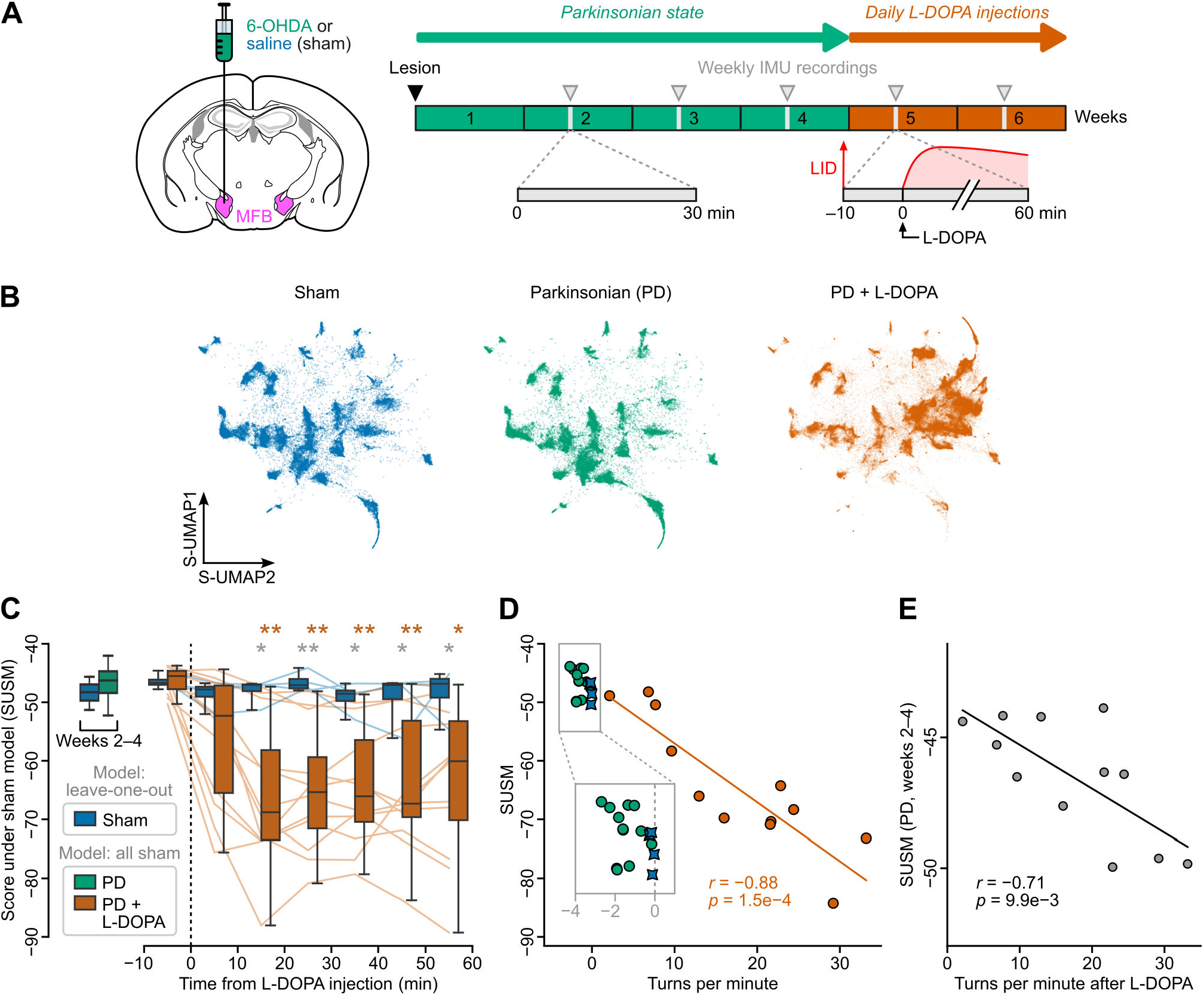
Macroscopic analysis of motor impairments in a mouse model of Parkinson’s disease and levodopa-induced dyskinesia. **A**, Experimental protocol. Left: drawing of a coronal brain slice (1.2 mm posterior to Bregma, adapted from [110]) showing the unilateral injection of 6-OHDA (PD mice, *n* = 12) or saline (sham mice, *n* = 4) into the medial forebrain bundle (MFB, purple). Right: protocol timeline (see Methods for details). The six-week protocol began with 6-OHDA (PD mice) or saline (sham mice) injection and IMU implantation. Weekly IMU recordings (gray arrowheads) were performed during weeks 2 to 4. Daily levodopa (L-DOPA) injections started in week 5, with IMU recording sessions conducted during weeks 5 and 6. Recordings began 10 min before injection. Each L-DOPA injection triggered an episode of abnormal involuntary movements (LID: L-DOPA induced dyskinesia, symbolized by a red curve). **B**, S-UMAP embeddings for sham mice (left, blue) and for PD mice before (middle, green) and during (right, orange) L-DOPA treatment, using a clustering model fitted on sham mice data. **C**, Left: box plots of the score under sham model (SUSM) for sham (*n* = 4) vs. PD (*n* = 12) mice before L-DOPA treatment (weeks 2 to 4). Right: box plots of the SUSM computed for the same mice over 10-min bins right before and after L-DOPA injection (weeks 5 and 6). Lines show the time course of the SUSM for individual PD (orange) and sham (blue) mice. Gray asterisks denote significant differences between PD vs. sham mice SUSM values for the same time bins (unpaired pairwise t-test with Bonferroni correction; *: p-value < 0.05; **: p-value < 0.01). Orange asterisks denote significant differences between PD mice SUSM values before vs. after L-DOPA injection (paired pairwise t-test with Bonferroni correction; *: p-value < 0.05; **: p-value < 0.01). **D**, Scatter plot of the SUSM vs. turns per minutes (estimated from IMU recordings, see Supplementary Methods) for each animal. Along the *x* axis (turns per minute), negative (resp. positive) values indicate ipsiversive (resp. contraversive) rotations relative to the lesion side. Each PD mouse (*n* = 12) is shown twice: during the Parkinsonian phase (green dots) and during the LID phase (10–60 min period following L-DOPA injection, orange dots). The linear regression was computed for the LID phase. **E**, Comparison of the SUSM before L-DOPA treatment (weeks 2 to 4) and the turns per minute during the LID phase (10–60 minutes following L-DOPA injection) for PD mice (*n* = 12), showing that the SUSM during the Parkinsonian phase is predictive of the amount of contraversive rotations induced by L-DOPA.

To quantify how much the motor repertoire of PD mice deviated from the one of healthy animals, we calculated the average log-likelihood of segments under the sham-GMM. To mitigate potential overfitting, the score for each sham animal was computed using a leave-one-out approach (see Methods). This “score under sham model” (SUSM) was only slightly lower in PD mice compared to sham mice before L-DOPA administration (Fig. 7C). After L-DOPA injection, the SUSM of PD mice decreased substantially (Fig. 7C), indicating that the sham-GMM poorly explained their behavior during the LID phase. Given the characteristic rotational behavior in this model and its inversion during the LID phase, we quantified rotations using IMU data as previously described [42]. PD mice displayed mild but significant ipsiversive rotations relative to the lesion side (−1.73 ± 0.80 turns per minute, vs. −0.12 ± 0.10 for sham mice; *p* = 1.1e−3, Wilcoxon test) before L-DOPA injection and a pronounced circling behavior in the opposite direction after injection (17 ± 9 turns per minute). Rotations during the LID phase negatively correlated with the SUSM (*r* = −0.89, *p* = 9.0e−5; Fig. 7D), suggesting that most of the L-DOPA-induced behavioral remapping captured by our pipeline is linked with an overrepresentation of contraversive movements. Notably, these rotations also negatively correlated with the SUSM measured before injection (*r* = −0.71, *p* = 9.5e−3; Fig. 7E), suggesting that changes occurring in the PD state are predictive of the overall response to L-DOPA.

To further characterize behavioral changes in PD mice, we examined how frequently sham-GMM clusters were used in these mice in comparison with sham mice (Fig. 8A). While the S-UMAP embedding appears similar between sham and PD mice before L-DOPA, this cluster usage analysis revealed clear behavioral differences. Before L-DOPA injections, PD mice predominantly employed two clusters: one corresponding to episodes of immobility and the other to slow, mobile olfactory exploration with ipsiversive turns. During the LID phase, cluster usage shifted to a series of five clusters, all associated with contraversive movements (Fig. 8B,C). Among clusters whose usage did not change drastically, we focused on one representing upward pitch movements, which exhibited a slight positional shift in the embedding space across groups and conditions (Fig. 8D). This shift occurred along a gradient of head roll angles, with PD mice segments concentrated on opposite sides of the gradient before and after L-DOPA injection. This rearrangement corresponded to a mild ipsiversive bias of upward pitch movements in PD mice, which reversed to the opposite side under L-DOPA (Fig. 8E). This example illustrates how DISSeCT can be used to single out subtle changes in motor patterns.

**Figure 8:**
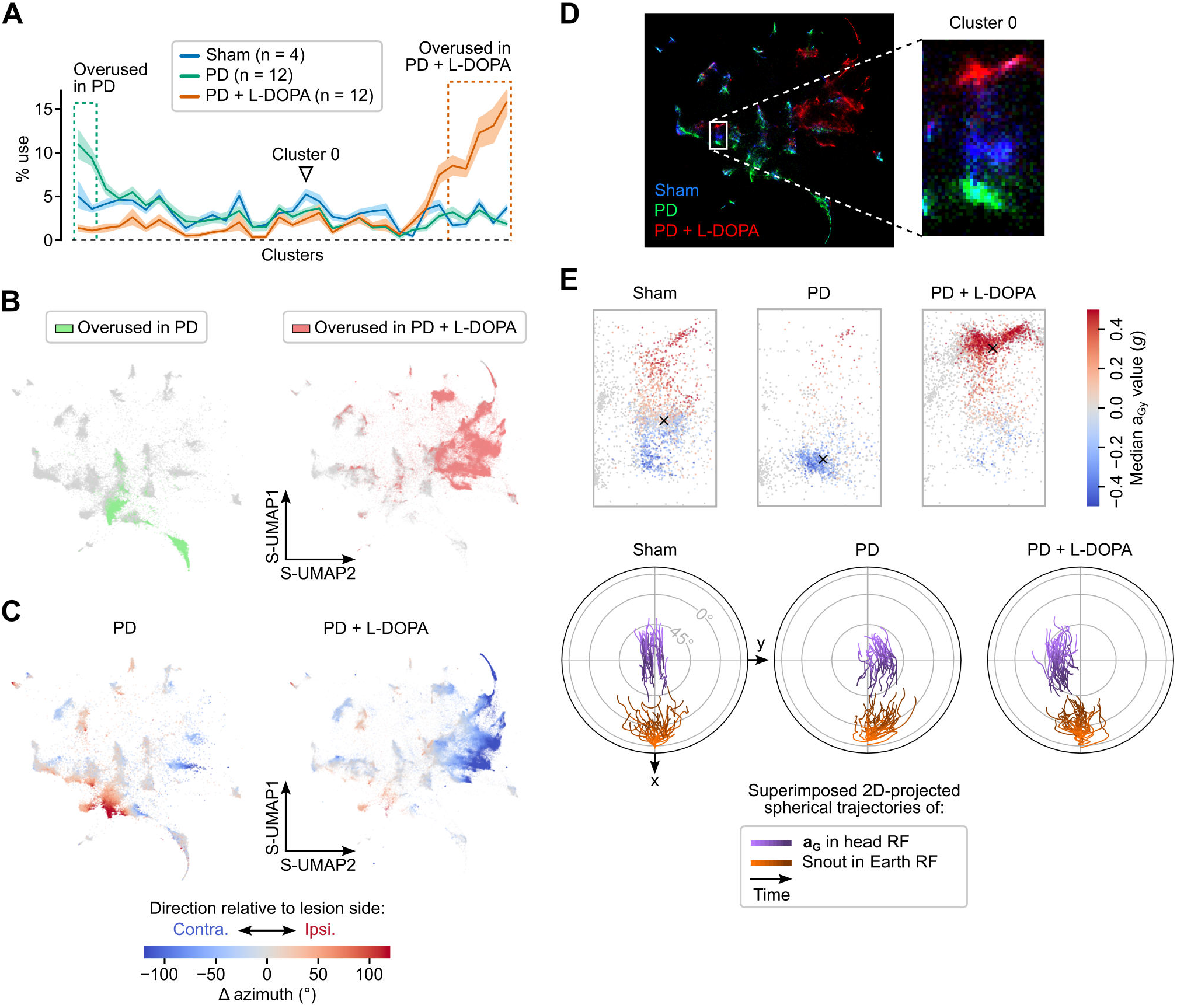
DISSeCT captures both coarse and fine-grained changes in a mouse model of Parkinson’s disease and levodopa- induced dyskinesia. **A**, Per-mouse average percentage use of individual clusters in sham (blue curve) vs. PD animals before (green curve) and after (orange curve) L-DOPA administration. Shaded areas represent 95% confidence intervals. Dashed rectangles indicate clusters exhibiting an increased use of more than 5 % compared to sham mice (overused clusters) for PD mice before (green rectangle) and after (orange rectangle) L-DOPA injection. **B**, S-UMAP embeddings highlighting segments belonging to overused clusters (defined in panel A) in PD mice before (left, green) and after (right, orange) L-DOPA injection. Gray points represent segments belonging to other clusters. **C**, S-UMAP embeddings showing segments color coded according to the change of head azimuth. Positive (resp. negative) values represent ipsiversive (resp. contraversive) rotations relative to the lesion side. **D**, Right: Superimposed S-UMAP embeddings for sham (blue), PD (green) and PD under L-DOPA (red) mice. Left: close-up view of cluster 0 (indicated by an arrowhead in panel A), which corresponds to upward head movements. **E**, Left: close-up view of cluster 0 in the S-UMAP embedding space, with segments color coded according to the median value of **a**_**G**_ along the interaural axis. Negative (resp. positive) values indicate a roll towards (resp. away from) the lesion side. Right: corresponding joint visualization of the trajectories of **a**_**G**_ in the head reference frame (purple) and of the animal’s heading in the Earth reference frame (orange) for the top 30 segments of cluster 0.

## Discussion

IMUs are becoming an increasingly popular tool for motion tracking in neurophysiological research, yet their potential for parsing rodent behavior remains largely untapped. Unsupervised approaches, which do not rely on human labels to uncover data structure, are essential to fully assess this potential. Drawing inspiration from previous work on accelerometer or GPS data collected from free-roaming animals [85–87] and egocentric videos recorded in a clinical context [88], we developed DISSeCT, a two-step unsupervised segmentation and clustering pipeline specifically designed for rodent IMU data.

Similar to current video-based unsupervised behavioral mapping procedures [13, 15, 16], DISSeCT was able to distinguish the main categories of rodent activities (Fig. 1), while also delivering fine-grained information on their various components. Our pipeline mapped the full repertoire of rapid three-dimensional head orienting movements (Fig. 2), offering a convenient alternative to methods based on threshold crossing detection in gyroscope measurements [39]. Interestingly, the duration and angular speed of the corresponding motifs co-varied to produce comparable head angular displacements (Fig. 2B), suggesting a default angular set point for orienting movements. DISSeCT also successfully isolated episodes of locomotion, characterized by a rhythmic ≃ 6 Hz pattern in head-vertical acceleration (Fig. 3). A similar signature at 8–10 Hz, scaling with animal speed (50–100 cm/s) was previously described in running rats [28]. The lower frequency observed here is probably related to the lower speed (10–15 cm/s) imposed by the restricted size of our arena. Inspection of a slow locomotion cluster revealed a ≃ 10 Hz signature of sniffing distributed along a specific combination of axes (Fig. 3D), and whose angular speed component had already been reported [31]. This signature, found in several other clusters (Fig. 4), could be used in the future as a standalone metric to assess the level of olfactory exploration. Finally, grooming activities emerged as a family of clusters with diverse time-frequency properties, uniquely characterized by an uncoupling of the correlation between roll and yaw rotations from the head pitch angle (Fig. 5A). These results demonstrate that DISSeCT is indeed a relevant approach for in-depth IMU data mining applied to rodent behavioral mapping.

A central question in behavioral mapping pipelines is whether a learned model can generalize to new observations. In our case, the learned GMM parameters—namely, centers, covariance matrices, and mixture weights—are inherently influenced by IMU placement, as variations in sensor orientation affect the comparability of extracted features. This constraint is not unique to our approach; it similarly applies to pose-based models, where meaningful comparisons require animals of similar size and anatomical proportions. Moreover, differences in environment and task structure can shape the behavioral repertoire expressed, further limiting the generalizability of a single model across diverse conditions. As discussed below, however, GMMs are ideally suited for model comparison, mitigating these intrinsic methodological limitations. In light of these factors, we recommend running new instances of our pipeline on new datasets rather than directly reusing the GMM parameters we derived, and using the BIC to tune the number of mixture components and select the appropriate covariance matrix (Fig. S4). Ideally, every new implementation should include a dedicated model exploration step. That said, future studies may still benefit from our detailed characterization, which robustly links distinct behavioral activities (e.g., locomotion, sniffing, grooming, and extended rearing) to specific regions of the feature space. To aid interpretability in novel experimental settings or animal models, we advise maintaining a minimal parallel video stream, as it supports posthoc visualization of cluster content. Our companion tool, INSpECT, streamlines this process by enabling fast browsing and labeling of relevant video snippets. Finally, we emphasize that DISSeCT’s speed and computational frugality, discussed below, lower the barrier to group comparisons across independently fitted models, while also enabling multiscale analyses through the iterative exploration of CPD penalty values to tune temporal resolution over selected recording periods.

Cluster inspection revealed that ethologically-distinct motifs may sometimes share similar head kinematic properties, such as upward head movements and rearing initiation. Using a post-hoc categorical HMM to model the symbolic sequence resulting from segment clustering (Fig. 6), we show that some of these ambiguities can be resolved if the corresponding motifs occur within distinct identifiable sequences (Fig. S11). Other solutions to improve the separation of similar motifs include fitting GMMs to the corresponding subsets of segments using tailored features or employing hierarchical clustering methods. Missing body kinematic information could be obtained using additional IMUs, either subcutaneously implanted [47, 51] or embedded into jackets worn by animals, keeping in mind that sensor position and orientation may vary over time [89]. Since maintaining a minimal concurrent video stream is a practical option for rapid cluster inspection, as discussed above, basic video-derived features could be integrated into DISSeCT to help resolve kinematic ambiguities. The potential utility of this approach is supported by our observation that cluster-wise purity scores were more robust to random sampling for KpMS syllables—derived from video—than for DISSeCT clusters (Fig. S12C–G). This hybrid strategy could enhance the pipeline’s discriminative power without substantially increasing experimental complexity. It would also allow social context–such as inter-animal proximity–to be incorporated into IMU-based analyses involving multiple interacting animals. In this setting, wireless IMU recordings would provide fully individualized and independent per-animal measurements [30], each of which could be processed separately using our pipeline. Provided that individual identities can be reliably tracked from video, this combined optical-IMU framework could enable the discovery of social interaction clusters, marked for example by high sniffing activity and close proximity to a conspecific.

In rodent research, IMUs have primarily been used for automatic behavioral scoring, typically relying on fixed-length data segmentation combined with supervised classifiers [26, 51, 52]. To our knowledge, only two main approaches have employed IMU recordings for unsupervised rodent behavioral mapping, with the goal of linking the resulting cluster sequence to concurrent recordings of neuronal activity. In one case, feature extraction over fixed-length windows was performed on combined IMU and video data, and behavioral motifs were uncovered by applying affinity propagation to a pairwise window similarity matrix [33, 90]. In the other, IMU data were used primarily to refine 3D head orientation estimates within a broader pose-based behavioral mapping framework [16]. In contrast, our approach represents, to our knowledge, the first unsupervised rodent behavioral mapping pipeline that relies exclusively on inertial recordings.

Our pipeline also differs in several ways from existing video-based, unsupervised rodent behavioral analysis frameworks, which can be broadly grouped into two categories. The first category relies on extracting features from fixed-length video segments and embedding them into two-dimensional maps, where recurring behavioral patterns manifest as high-density areas. This strategy is employed in pipelines such as MotionMapper [70], B-SOiD [77], and DANNCE [13]. While effective for broad behavioral mapping [15–17], this approach is not designed to precisely delineate behavioral motifs in time. Additionally, its drastic dimensionality reduction prior to clustering potentially diminishes its discriminative power [71]. In contrast, DISSeCT uses CPD to accurately locate behavioral transitions and performs clustering in a high-dimensional space, minimizing information loss. The second category aims at explicitly modeling temporal behavioral dynamics using graphical models. For example, Keypoint-MoSeq [18] combines autoregressive HMMs with switching linear dynamical systems to jointly infer behavioral motif limits and identities from PCA-reduced keypoint trajectories. Another pipeline, VAME [63], occupies a middle ground between these two families: it first uses representation learning to construct a lower-dimensional embedding of pose trajectories from fixed-length windows, then applies an HMM to simultaneously segment and cluster the resulting latent time series.

Here, we propose an alternative approach in which time series segmentation is decoupled from the clustering step, with two main objectives. The first is to improve computational efficiency. In our conditions, a full CPU-based DISSeCT analysis ran approximately five times faster than a GPU-based KpMS analysis (see Tables S4 and S5), even before accounting for the additional time and resources required for video-based pose estimation. Both methods operated on a similar number of dimensions (DISSeCT used a 9-dimensional inertial time series, while KpMS modeled an 8-dimensional latent time series), though a key difference was that IMU data were sampled 10 times faster (300 Hz vs. 30 Hz). Given that KpMS’s computing time and GPU memory usage scale linearly with dataset size, applying it directly to IMU data would increase runtime and memory requirements by a factor of at least 50. This makes such an approach impractical for routine use in large-scale or long-duration behavioral recordings, unless significant computational resources—such as high-performance GPU clusters—are readily available. The second objective, discussed below, is to maintain a modular and adaptable pipeline.

DISSeCT’s modular design allows for the integration of alternative clustering methods or iterative processes, such as additional rounds of feature extraction and segment clustering for finer-grained mapping of specific data subspaces. Automated scoring of user-defined activity patterns could also be achieved using a supervised learning algorithm downstream of the CPD step. Regardless of whether segments are analyzed using supervised or unsupervised methods, feature design and selection remains a critical component of the pipeline. A promising direction involves formalizing feature extraction and dimensionality reduction through representation learning, an approach that seeks to uncover the most informative nonlinear embeddings prior to the modeling stage. For instance, automatic feature engineering was achieved using variational autoencoders in VAME, a recent pose-based, unsupervised analysis pipeline [63]. This technique, however, is currently constrained by its reliance on fixed-length input windows. Extending such methods to accommodate variable-length segments, such as those derived from CPD, would require further methodological advances, potentially leveraging transformer-based architectures [91].

The combined resolution and versatility of IMU recordings open up a wide range of potential applications for our pipeline. By accurately detecting abrupt behavioral transitions and parsing brief behavioral motifs, DISSeCT provides the temporal resolution required for neuroethological studies aimed at uncovering the neuronal mechanisms underlying natural behavior [2]. One particular area where our pipeline could significantly advance the state-of-the-art is in analyzing behaviors that are challenging to accurately assess via video. Self-grooming, for example, is a complex behavior associated with contorted postures where IMUs offer considerable advantages over video recordings for in-depth analysis [16, 26, 51]. DISSeCT successfully parsed the main components of self-grooming, shedding light on their distinct head kinematic signatures. While further refinement is needed to enhance the separation of grooming subcomponents, our approach introduces a powerful new method for automating the analysis of this elaborate syntactic behavior [83], with potential applications in the study of rodent models of neurological and psychiatric disorders [92]. Similarly, wet dog shakes events were easily isolated due to their prominent and unique spectral signature (Fig. S5), accounting for the vast majority of outlier segments (Fig. 1C). Their study is relevant in the context of rodent models of diseases including acute seizures [93], morphine abstinence [94], and nicotine withdrawal [95].

DISSeCT’s use of a GMM layer facilitates direct comparison between experimental groups. The probabilistic formulation of GMMs enables cross-likelihood evaluation, where a model trained on one dataset can be used to assess how well it explains a separate set of observations, and vice versa. We illustrated this approach in a mouse model of Parkinson’s disease and levodopa-induced dyskinesia, where our pipeline enabled the identification of both global and subtle motor alterations (Fig. 7 and 8). Alternatively, GMMs fitted independently on different datasets can be compared directly using entropy-based measures such as the Jensen–Shannon divergence, which quantify differences in the structure of behavioral distributions. By transforming high-resolution kinematic data into interpretable and comparable probabilistic representations, our pipeline offers a promising new solution for deep behavioral phenotyping. Potential applications could be found in the detection, quantification, and characterization of behavioral changes—both within and between individuals—across natural processes (e.g. development or aging) or in response to experimental interventions (e.g. in disease models, learning protocols or environment manipulations).

As outlined in the Introduction, inertial recordings bypass many of the common limitations of video recordings, but come with their own constraints—chiefly, the requirement for surgical implantation. Deploying IMUs in large cohorts can therefore entail substantial time investment compared to video-based strategies. This limitation is less relevant in neurophysiological studies, where animals are routinely implanted with recording devices that often already include an IMU. A key advantage of IMUs is their ability to track animals in large, naturalistic environments, including occluded or hidden spaces. Transitioning to such settings requires wireless IMUs, which introduces a necessary trade-off between sampling rate (a key determinant of power consumption) and battery life. Although low-power, data-logging IMUs offer a promising solution, they have yet to be adapted for use in small animals such as mice. Another limitation is that, unlike video, inertial signals are not readily interpretable by an untrained observer. To address this, both our previous study [42] and the present work introduce a set of metrics and graphical tools to aid in the exploration and interpretation of IMU data. Based on the output of our pipeline, we also highlight distinct head kinematic signatures that reliably map onto specific behaviors. Together, these resources aim to support and standardize future IMU-based behavioral investigations.

In conclusion, we propose and validate a novel unsupervised framework for the analysis of rodent inertial data that combines kernel change-point detection and probabilistic modeling. By leveraging the unique properties of IMUs, our pipeline enables access to an underexplored region of the “behavioral science space” [96], where high-resolution, fine-grained behavioral analysis can be conducted with minimal environmental constraints and computational cost.

## Material and Methods

### Animals

Four adult male Long-Evans rats (Janvier Labs) and sixteen RJOrl:SWISS mice (CD-1, eight males and eight females, Janvier Labs) were used in this study. They were group-housed under standard conditions (12-hour non-inverted light/dark cycle, 21–24 ^°^C, 50–60 % humidity) and provided with unrestricted access to food and water along with appropriate cage enrichment such as cardboard tunnels, shredded paper, and dental cotton rolls.

### Surgical procedures

#### Wired IMU fixation in rats

A wired IMU measuring 5.0 × 7.5 × 3.0 mm and weighting 0.3 g (mptek.fr), housing a 9-axis digital inertial motion sensor (MPU-9250, Invensense) and operated via a dedicated USB acquisition board (mptek.fr) was used for rat recording sessions. The IMU featured a 2 × 4 pin header (TSM-104-01-S-DV, Samtec Inc.) used to connect the device to its counterpart female socket header (NPPC042KFMS-RC, Sullins Connector Solutions), which was permanently affixed to the animal’s skull through a surgical procedure outlined below. Animals underwent this procedure at six weeks of age and were allowed one week to recover before starting the recording sessions.

The connector implantation surgery was carried out as follows. Preemptive analgesia was ensured by subcutaneously injecting buprenorphine (0.01 mg/kg) thirty minutes before the procedure. Anesthesia was induced with 3% isoflurane in oxygen delivered at 3 L/min. The animal was then positioned in a stereotaxic apparatus, with its eyes protected by ophtalmic gel. Throughout the remainder of the procedure, body temperature was kept at 37 ^°^C using a homeothermic monitoring system, while anesthesia was maintained with 1% isoflurane delivered at 0.5 L/min. The animal’s scalp was shaved and cleansed with povidoneiodine followed by 70% ethanol. Prior to incising the scalp, a subcutaneous injection of lidocaine (2%) was administered. The periosteum of the skull was gently scraped using a scalpel blade, followed by application of a 3% hydrogen peroxide solution. The female socket header, held upright using one arm of the apparatus, was then affixed to the cranial surface using self-curing dental adhesive (Super-Bond C&B, Sun Medical). Finally, the skin around the connector was sutured both anteriorly and posteriorly, and the animal was placed on a heating blanket while its recovery was monitored. Once fully alert and mobile, the rat was returned to its home cage.

#### 6-OHDA lesion and wireless IMU fixation in mic

A wireless IMU (WEAR wireless sensor v2.1, Champalimaud Scientific Hardware Platform) measuring 14 × 6 × 26 mm and weighting 1.85 g, housing a 9-axis digital inertial motion sensor (MPU-9250, Invensense) and operated via a dedicated USB acquisition board (WEAR basestation, Champalimaud Scientific Hardware Platform), was used for mouse recording sessions. The IMU was mounted on the head using a 1 × 4 pin header (M52-040023V0445, Harwin Inc.) on the IMU side and its counterpart female socket header (M52-5000445, Harwin Inc.) on the animal side (see surgical procedure below).

Unilateral nigrostriatal dopaminergic denervation was achieved at 9–10 weeks of age by injecting 6- hydroxydopamine hydrochloride (6-OHDA, Sigma-Aldrich) into the medial forebrain bundle (MFB) as previously described [97, 98]. A subcutaneous dose of buprenorphine (0.1 mg/kg) and an intraperitoneal dose of desipramine (25 mg/kg, Sigma-Aldrich) were administered thirty minutes before initiating the procedure. The initial steps of the surgery, from anesthesia induction to skull preparation, were similar to the rat connector implantation protocol described above. For the Parkinsonian model (PD mice), the left MFB (−1.2 mm AP, 1.25 mm ML and −4.7 mm DV relative to Bregma) was injected with 1 µL of a 6-OHDA solution (3.2 µg/µL free-base concentration in 0.02% ascorbic acid, Sigma-Aldrich); whereas for control animals (sham mice), an equivalent volume of the vehicle solution (0.02% ascorbic acid) was administered. The injection was performed with a Hamilton syringe at a rate of 0.3 µL/min, with the syringe maintained in situ for two minutes before and four minutes after injection. The female socket header used for IMU fixation, held upright using one arm of the apparatus, was then affixed to the cranial surface using self-curing dental adhesive (Super-Bond C&B, Sun Medical). The skin around the connector was sutured both anteriorly and posteriorly, and the animal received a subcutaneous injection of Meloxicam (5 mg/kg, Metacam, Boehringer Ingelheim) and of a glucose-saline solution (Glucose 5%, 10 mL/kg, Osalia). Throughout the recovery phase, the mouse was placed on a heating blanket while being closely monitored. Once fully alert and mobile, it was returned to its home cage. To mitigate the risk of post-surgical complications and distress, mice received daily injections of glucose-saline solution and were nourished with a blend of Blédine (Blédina, Danone) and concentrated milk for a period spanning 6 to 10 days post-surgery, resulting in a 100% survival rate.

### Data acquisition

#### Synchronized inertial and video data acquisition for quantifying rat behavior

Rat recordings started one week after connector implantation. Once equipped with the wired IMU, rats were placed in a 70 × 70 cm arena with transparent glass floor and transparent PMMA walls, allowing for video monitoring from multiple angles, including from underneath, similarly to previous studies [47, 77, 99]. Inertial signals were acquired at 300 Hz with accelerometer and gyroscope ranges set at ±2 *g* and ±1000 ^°^/s, respectively. Video recordings were captured using five off-the-shelf industrial cameras (DMK 37BUX273, The Imaging Source) connected via USB to an acquisition computer. Frame acquisition was triggered at 30 Hz by an Arduino Uno running a custom pulse-width modulation script. These triggers were simultaneously recorded by the IMU acquisition board to precisely timestamp video frame acquisition within the IMU data stream. In our conditions, occasional frame drops were observed, at a frequency which gradually increased during continuous acquisition. We found that delivering triggers in one-minute chunks (consisting of 1800 frames), separated by three-second pauses, and saving the corresponding videos as separate files, drastically reduced the occurrence of frame drops to 0.023 % (0.39 ± 0.65 missing frame per chunk).

#### Synchronized inertial and video acquisition for quantifying behavior in a mouse model of Parkinsons’s disease and levodopa-induced dyskinesia

Considering that 6-OHDA injection and connector implantation were conducted at day 0, IMU recordings started on day 10. Once equipped with the wireless IMU, mice were placed in a 30 × 35 cm arena with transparent PMMA floor and walls. Video recordings were captured using two off-the-shelf industrial cameras (DMK 37BUX273, The Imaging Source) positioned below and on the side of the arena. Synchronized acquisition of video and IMU data was carried out using dedicated acquisition hardware (WEAR basestation, Champalimaud Scientific Hardware Platform) controlled via a custom pipeline programmed in Bonsai [100]. Videos were acquired at a rate of 50 Hz, while IMU data was recorded at 200 Hz with accelerometer and gyroscope ranges set at ±2 *g* and ±1000 ^°^/s, respectively.

Recordings were conducted over a five-week period, with one recording session per week (Fig. 7A). The first three sessions (days 10, 17 and 24) lasted 30 minutes each. Starting on the 24^th^ day, mice received daily intraperitoneal injections of L3,4-dihydroxyphenylalanine methyl (L-DOPA or levodopa, 3 days at 3 mg/kg and then 6 mg/kg, Sigma-Aldrich) and benserazide hydrochloride (12 mg/kg, Sigma-Aldrich), a peripheral decarboxylase inhibitor. The final two recording sessions were conducted on days 31 and 38, starting 10 minutes before and extending 60 minutes after levodopa injection.

### Markerless 3D pose estimation

In the case of rat video recordings, the use of five different viewpoints enabled 3D pose estimation. Using the DeepLabCut toolbox [6], a unique deep neural network was trained to recognize twelve anatomical keypoints from all viewpoints, providing an independent 2D pose estimate for each video. The training dataset consisted of 1 950 hand-labeled frames selected to encompass all different viewpoints and a wide range of animal positions and postures within the arena. Additional details regarding the parameters employed can be found in Table S1. Before each session, the intrinsic and extrinsic parameters of the cameras were estimated by recording a video of a ChArUco board moved throughout the arena, and applying the calibration pipeline available in Anipose [12]. This pipeline combines checkerboard detection using OpenCV with an optimization procedure aimed at minimizing reprojection errors.

Spatio-temporal filtering is a crucial step in obtaining high quality estimates of 3D pose dynamics [12, 47, 101]. Fot this purpose, we used the filtering pipelines integrated into Anipose. Median filtering was first applied to the five independent 2D pose estimates to alleviate the impact of sudden keypoint jumps. Triangulation was then performed using camera calibration and spatio-temporal regularization, taking advantage of the continuity and relative smoothness of animal movements, as well as the stability of bodypart lengths over time. Finally, a 3D median filter was applied to the resulting 3D pose estimate to reduce keypoint jitter. A summary of the parameters used for triangulation and filtering with Anipose is provided in Table S2.

### Pre-processing of IMU data

#### Subtraction of sensor offsets

The estimation of sensor offsets followed a procedure described previously [42]. For each IMU, data was collected while placing the device in a set of 50 different static positions using a custom robotic arm. Keeping only the periods of immobility, accelerometer offsets were estimated by minimizing the average squared difference between the Euclidean norm of offset-corrected acceleration measurements and 1 *g*. Gyroscope offsets were computed as the median gyroscope values observed during these periods of immobility. Accelerometer and gyroscope offsets were then subtracted from acceleration and angular velocity measurements, respectively.

#### Estimation of head tilt

Head orientation relative to gravity (head tilt) can be accurately estimated from rodent IMU data using the appropriate Attitude and Heading Reference System (AHRS) filtering algorithm. For each IMU data point, an extended Kalman filter [102] with a previously validated set of parameters (*var*_*acc*_ = 0.002, *var*_*gyr*_ = 0.75) [42] was used to compute a unit quaternion describing sensor orientation relative to an arbitrary, gravity-polarized Earth reference frame (absolute reference frame). This quaternion represents a 3D rotation which, when applied to this absolute reference frame, aligns it with the sensor’s orientation. To estimate head orientation relative to gravity (head tilt, i.e. the orientation of the gravitational acceleration vector in the head reference frame), the inverse rotation, corresponding to the conjugate quaternion, was applied to the unit vector (0 0 1).

### Change-point detection

#### Algorithm

Pre-processed IMU data were segmented by applying the PELT (Pruned Exact Linear Time) change-point detection method to z-scored gravitational acceleration and angular velocity time series (non-gravitational acceleration was not used here because of its lower signal-to-noise ratio). This algorithm leverages dynamical programming to find the optimal number and location of change points in time series by minimizing a cost function (see Supplementary Methods), while maintaining a linear computation time [65]. Specifically, we used a kernel-based change-point detection algorithm [64] relying on a Gaussian kernel, which offers the advantage of being characteristic; meaning that changes in the mean of the embedded data correspond to changes in the data distribution [69, 103, 104] (See Supplementary methods).

### Selection of the penalty parameter

Penalty for segmenting IMU data was set to *λ* = 14, yielding segments with a median duration of 367 ms, to align with the typical duration of individual elements identified by MoSeq, a depth video-based unsupervised method used to parse mouse behavior [62]. This duration also matched the timescale of the finest behavioral motifs identifiable by a human observer (see Fig. S2).

### Segment clustering

#### Feature extraction

A total of 240 features were extracted for each segment, effectively capturing postural and kinematic aspects of the animal’s activity within the segment. These features included statistics on the signals, parameters derived from head tilt and changes in head azimuth, as well as coefficients from continuous wavelet transforms of angular velocity and non-gravitational acceleration data (see Supplementary Methods).

#### Outlier removal

A small subset of segments exhibited outlier feature values, leading to convergence issues during clustering. To address this, a threshold of 35 was applied to absolute robustly rescaled feature values, obtained through median subtraction and division by the interquartile range. Segments exceeding this threshold were excluded from the analysis. These outliers consisted of extended periods of immobility (17 %) and high magnitude events corresponding either to vigorous movements (21 %, typically > 1 *g* of non-gravitational acceleration and > 500 ^°^/s of angular speed) or to wet dog shakes [105] (with the norm of non-gravitational acceleration and angular velocity exceeding 2 g and 1000 ^°^/s, respectively). The latter type of outlier is characterized by a high amplitude 16–20 Hz oscillatory regime whose wavelet transform signature artificially contaminates neighboring segments due to edge effects (Fig. S5). Consequently, segments immediately before and after wet-dog shake events were also excluded from the analysis. This curation procedure resulted in the exclusion of 1.7% of all segments, representing 3.7% of the total recording duration.

#### Gaussian Mixture Model clusterin

After having excluded outlier segments as described above, features were z-scored and projected in a lower-dimensional space via PCA. The number of principal components was determined through cross-validation [106] (Fig. S4A). A clustering step was then executed on the resulting projection using scikit-learn’s GMM implementation. Model selection and hyperparameter tuning were conducted through grid search with the aim of maximizing the Bayesian information criterion (BIC) [107]. The grid search encompassed covariance type (full, diagonal, tied, spherical) and number of clusters (ranging from 5 to 100), employing 10 random initialization for each hyperparameter set (Fig. S4B), yielding 48 mixture components with full covariance matrices. In our terminology, *clusters* are groups of segments assigned to the same most likely generating mixture component, as determined by posterior probabilities. Accordingly, this hard assignment step yielded 48 clusters, each corresponding to a mixture component in the GMM. These clusters are defined in the PC space and do not correspond *a priori* to behavioral categories. Their ethological relevance is assessed in subsequent analyses. More information on the procedure is available in the Supplementary Methods.

#### Ablation of immobility periods prior to modeling

To evaluate whether a CPD-based segmentation strategy provided a better fit than a simple fixed-length windowing approach, we performed an ablation of immobility periods prior to Gaussian mixture modeling. Fixed-length windows falling within immobility periods were identified using a threshold of 12 ^°^/s on the standard deviation of angular speed (i.e., the norm of gyroscope measurements), as illustrated in Fig. S2D. This procedure excluded 7.3 % of all windows (3 222 out of 44 426), totaling 2255.4 s. Because immobility segments isolated by CPD often corresponded to long periods of immobility briefly interrupted by small movements, the same threshold-based exclusion could not be directly applied. Instead, we ranked CPD segments by angular speed standard deviation and incrementally removed the lowest-activity segments until their total duration matched that of the excluded fixed-length windows. This resulted in the removal of 2.05 % of all CPD segments (908 out of 44 322), corresponding to a total duration of 2255.1 s. After feature extraction and PCA, we fit a series of 10 independent GMMs over a range of component numbers (*K*), using either the full or ablated sets of segments/windows. For each *K*, the average log-likelihood was computed on the corresponding set of segments/windows.

#### Two-dimensional embedding of clustered data

For visualization, z-scored features associated with individual segments were projected onto a two-dimensional space using supervised UMAP embedding [108] (with parameters: *n*_*neighbors* = 8, *target*_*weight* = 1*e* − 9, *min*_*dist* = 1*e* − 3). This method incorporates cluster label information to guide the embedding process, ensuring that points belonging to the same cluster are placed closer together, while still preserving the overall structure of the data. Feature values computed on outlier segments, excluded from the clustering step, were also included in this embedding step, albeit ignoring the associated label.

#### Sparse clusters

The dispersion of individual mixture components was evaluated by calculating the trace of their covariance matrix. Clusters associated with the three mixtures exhibiting the highest trace values were considered as sparse clusters (Fig. S7A). Collectively, these clusters accounted for 2.1% of the total number of segments and for 1.0% of the total recording duration. Examination of these clusters revealed that they comprised heterogeneous segments, corresponding to occasional brief and vigorous movements (approaching 2 *g* of non-gravitational acceleration and 1000 ^°^/s of angular velocity), and which appeared highly dispersed on the UMAP embedding (Fig. S7B). The presence of such clusters is expected, as GMMs tend to assign rare and/or outlier observations to a few highly dispersed, poorly informative clusters [109].

### *Ad hoc* metrics

#### Per-segment average head roll and pitch angles

Within the head’s reference frame, gravitational acceleration can be conceptualized as a unit position vector, more specifically as a point on a head-centered sphere of radius 1 *g* that fully represents head tilt. Due to the non-linearity of trigonometric functions, calculating the average head tilt over a certain period of time is not as straightforward as averaging the coordinates of this point. Instead, the average head tilt can determined as the centroid (estimated mean direction) of a von Mises-Fisher distribution fitted to the various positions assumed by this point on the unit sphere. Noting this centroid 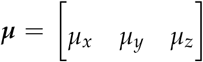, the corresponding average head roll and pitch angles are computed as the opposite values of the polar coordinates of ***µ***:

- *µ*_*roll*_ = −*arctan*2(*µ*_*y*_, *µ*_*z*_)
- 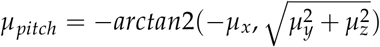

#### Detection of immobility

Angular speed was computed as the Euclidean norm of offset-corrected gyroscope measurements and then smoothed using a Gaussian kernel (*σ* = 100 ms). The animal was considered immobile when this value was below 20 ^°^/s.

#### Sniffing score

To identify segments displaying the ≃ 10 Hz sniffing signature primarily observed in the naso-occipital non-gravitational acceleration (noted *a*_*nG,x*_) and pitch angular velocity (noted *ω*_*y*_), we designed an *ad hoc* sniffing score (noted Φ). This metric was constructed empirically using the maximal wavelet power (noted *W*) across all available frequencies (1–20 Hz) or frequencies largely above (> 14 Hz), below (< 6 Hz) or within the band of interest (8.5–11 Hz):

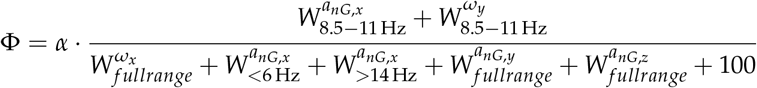

where *α* is a factor that penalizes Φ for segments associated with rapid head reorientation or large roll angles, which often exhibit non-sniffing related 8–10 Hz signatures. Brief reorientations can indeed last approximately 100 ms, thereby displaying a ≃ 10 Hz time-frequency signature, while substantial roll angles are linked to body grooming, an activity with a typical ≃ 4 Hz pattern that may manifest a notable harmonic at ≃ 8 Hz. This factor was computed as follows:

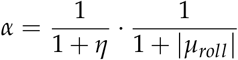

with *η* representing the heading change rate in ^°^/s (the angle between heading estimates at the beginning and end of a segment, divided by its duration) and *µ*_*roll*_ the roll of the average tilt in radians.

### 2-dimensional visualization of per-segment head attitude and heading trajectories

To visualize 3D head orientation during individual segments, we overlaid the trajectories of head-tilt (pitch and roll) and 3D-heading (pitch and azimuth). Head-tilt was represented using the coordinates of the Earth’s vertical in the head reference frame, estimated by the gravitational component of acceleration signals obtained through AHRS filtering of IMU data (see Methods). The estimate of 3D-heading was derived by applying the rotation from the head reference frame to a gravity-polarized Earth reference frame, achieved via AHRS filtering, to the x-axis of the sensor’s reference frame. The resulting vector was then rotated along the vertical axis to set the first sample of the segment to an azimuth of 0 in the Earth reference frame (see Supplementary Methods).

These 3-dimensional time series of unit-norm vectors were projected to two dimensions using a Lambert Azimuthal-Equal Area projection, centered either on the *North Pole* (i.e., (0, 0, 1) or *South Pole* (i.e., (0, 0, −1) of the respective reference frames. The trajectories were then superimposed on a single plot using the cartography module cartopy.

### Post hoc categorical HMM

A categorical hidden Markov model was fitted to the sequence of clusters obtained from GMM modeling, using the implementation provided by hmmlearn. Segments deemed outliers, as defined earlier, were treated as an independent cluster. Hyperparameter optimization was conducted via BIC maximization, employing ten independent fits per hyperparameter value, to determine the optimal number of states. More information on the procedure is available in the Supplementary Methods.

### Keypoint-MoSeq analysis

To analyze our 3D rat pose dataset using the Keypoint-MoSeq (KpMS) pipeline [18], we first optimized the stickiness hyperparameter (*λ*) to achieve a median syllable duration of 367 ms, aligning with previously reported values [18, 62] and with the median segment duration derived from change-point detection applied to IMU data (367 ms; see above). We then fit six independent KpMS models to the dataset and selected the best-performing one based on the expected marginal likelihood (EML) score [18]. All parameters used for running KpMS are listed in Table S3. Computation times for the GPU and CPU versions of KpMS are reported in Table S5.

Behavioral syllables obtained from KpMS were compared to DISSeCT clusters by aligning each video frame to the corresponding IMU sample. This alignment was based on detecting the rising edge of the trigger pulse that initiated video frame acquisition, as recorded in the IMU data stream. A contingency matrix (Fig. S13) was then constructed by counting the number of co-occurring IMU-video frame pairs assigned to each KpMS syllable and DISSeCT cluster.

### Inspection and labeling of video segments

#### Software

To expedite the inspection and labeling of video snippets corresponding to selected DISSeCT segments or KpMS syllable instances, we contracted AquiNeuro to develop custom software. This software, named INSpECT (Interface for Navigating Spates of video Excerpts and Categorizing Them), features a graphical user interface and keyboard shortcuts for navigating a series of multi-camera video snippets and assigning user-defined labels to each. The code of INSpECT is available at https://github.com/rfayat/INSpECT.

#### Inspection and labeling

Given a set of video snippets corresponding to selected DISSeCT segments or KpMS syllable instances, an observer was tasked with assigning a unique primary category to each one (choosing from the following options: “orienting”, “rearing”, “grooming”, “locomoting”, “idle” or “other”), along with an additional unconstrained secondary label providing a more detailed description of the ongoing behavioral activity. The observer was blind to the identity of the DISSeCT cluster or KpMS syllable from which each snippet originated. The category “orienting” was used for rapid changes in head orientation unrelated to rearing or grooming episodes. “Rearing” referred to situations during which the animal stood on its hind legs with both forelegs above the ground. “Grooming” encompassed all types of self-cleaning activities. The category “idle” applied to situations where the animal stood immobile, occasionally accompanied by small twitches or head sways.

To quantify how consistently each cluster or syllable was associated with a specific behavior, we computed a *purity score* [63], defined as the proportion of snippets assigned to the most frequent category:

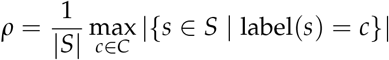

where *S* is a set of video snippets *s* associated with a given cluster or syllable, and *C* is the set of behavioral categories *c*.

Prior to snippet selection, the following eligibility criteria were applied: (1) a minimum duration of 7 video frames (233 ms) to ensure visibility of behavior by the observer, (2) DISSeCT segments had to be classified as non-outliers (see Methods) and fully contained within video chunks (see Methods), (3) KpMS instances had to originate from syllables accounting for more than 5 % of the total recording duration. Applying these criteria yielded 41 005 eligible DISSeCT segments (out of 44 322) and 30 230 eligible KpMS syllable instances (out of 48 854).

Snippet selection was then performed in one of two ways: (1) by selecting the top one hundred most representative segments or instances per cluster or syllable, or (2) by randomly sampling 10 % of all eligible segments or instances. For DISSeCT, the top one hundred segments per cluster were defined as those with the highest posterior probabilities assigned to that cluster. For KpMS, we used the best-performing model to estimate frame-wise marginal probabilities over syllable labels. Consecutive frames with the same highest-probability label were grouped to form new syllable instances. The instance-level probability was computed as the average of the frame-wise probabilities and used to rank instances. For each syllable, the top one hundred instances with the highest scores were selected.

### Mapping of healthy and pathological mouse behavior

#### Pipeline implementation

Change-point detection was conducted on all available mouse IMU data. Similar to the approach taken with rat data, the penalty level was set to a value (*λ* = 9 here) resulting in a median segment duration of ≃ 350 ms. The same set of features used for rats was extracted for each segment, and the identical procedure for detecting outlier segments was applied, leading to the exclusion of 0.7 % of all segments from the clustering step. These segments primarily consisted of extended periods of immobility and accounted for 10.6 % of the total recording duration.

Parameters for feature rescaling, PCA dimensionality reduction and Gaussian mixture modeling were solely fitted on sham (healthy) mouse recordings before levodopa treatment. Hyperparameter optimization based on BIC maximization yielded 33 clusters with full covariance matrices. Based on this model, cluster assignment was then carried out for all other non-outlier segments (from sham mice under levodopa treatment and PD mice in all conditions).

The final supervised UMAP embedding was fitted using only one out of three segments for computational efficiency and using the same hyperparameters as for rat data, and was then applied for all mice and conditions.

#### Score under a sham model

The GMM fitted on segments from sham mice prior to levodopa treatment served as the reference model for computing the log-likelihood of all available segments. For any given segment, this value indicates how likely the segment is under the assumption that the mouse is healthy. The average log-likelihood across a set of segments, referred to hereafter as the *score under a sham model* (SUSM), was used as a measure of how well the corresponding IMU recording bout could be explained by the healthy behavioral model.

To prevent potential overfitting when evaluating sham animals, we adopted a leave-one-out approach: for each sham mouse, the SUSM was computed using a GMM trained on all other sham animals, excluding the test animal. This allowed for a more accurate estimate of model generalization within the sham group. In contrast, segments from PD and L-DOPA mice were evaluated using a single GMM trained on the full sham dataset, reflecting the use of this model as a normative reference. All GMMs used identical hyperparameters (number of components and covariance structure) as determined by BIC on the full sham data.

#### Overlay of behavioral maps

To overlay UMAP scatter plots (Fig. 8D), we first transformed them into two-dimensional histograms using a 500 × 500 grid. The bin values were then rescaled to fall between 0 and 1 by dividing each value by the 99.8^th^ percentile of the histogram, with any values exceeding one clipped to one to mitigate the impact of outliers. The resulting histograms were visualized as an image using the red, green and blue channels for PD-LDOPA, PD and sham mice, respectively.

## Code availability

The code of DISSeCT is available at https://github.com/rfayat/DISSeCT. The code of INSpECT, a companion software used to examine and label video snippets, is available at https://github.com/rfayat/INSpECT.

## Acknowledgements

We extend our gratitude to Ahmed Boughdiri and Stefano Bettani for their assistance in setting up the multi-camera video acquisition system, and to Idriss Tsayem Ngueguim for his help in rat implantation surgeries. We also thank Ana Margarida Pinto for her help in establishing the acquisition system used for mouse recordings, Adja Sissokho for her assistance with mouse experiments and John Jacoby for his guidance in setting up Keypoint-MoSeq analysis. Additionally, we are grateful to Laurent Oudre, Éric Burguière, Ashesh Dhawale and Bence Ölveczky for their valuable feedback on the project. Our thanks also go to the IBENS computing platform, fablab, and animal facility for their support.

## Author contribution

R.F. assembled and programmed IMU and video recording setups, designed and performed rat experiments, co-designed the architectures of DISSeCT and INSpECT, wrote the DISSeCT code, performed the analysis, generated the figures and drafted the manuscript. M.S. implemented the mouse model of Parkinson’s disease and levodopa-induced dyskinesia and conducted mouse recordings. D.P. and C.L. contributed to project design and edited the manuscript. P.L. and G.D. co-supervised R.F.’s work, co-designed the DISSeCT pipeline and wrote the manuscript.

## Ethical approval

This project has undergone review and approval by the ethical review board and the department responsible for overseeing the use of animals for scientific purposes within the French Ministry of Higher Education, Research, and Innovation. Approval was granted under reference APAFIS #29793-202102121752192 v3.

## Supplementary Methods

### Change-point detection

#### Principle and cost function

Given a time series *x* = (*x*_1_, …, *x*_*N*_) of length *N*, unsupervised change-point detection aims to find both the number *K* and the locations 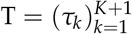 of significant changes in the signal or its dynamics. For a given segmentation T, the total cost is defined as the sum over all segments of a function *c*(·) that quantifies the internal homogeneity of each segment, penalized by a term that increases with the number of segments *K*:

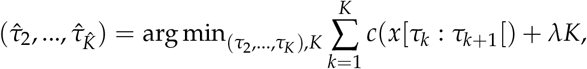

where *τ*_1_ = 1 and *τ*_*K*+1_ = *N* + 1 are both fixed by convention. We used a Gaussian kernel-based rbf cost function *c*_*Gaussian*_, which is sensitive to changes in the data distribution [111]. Using the kernel trick, the homogeneity cost of a segment *x*[*a* : *b*[is computed as:

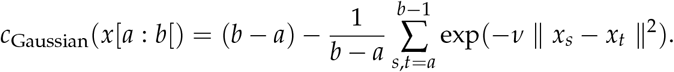

The additive term (*b* − *a*) arises from the general formulation of kernel-based cost rbf functions, where it represents the number of elements in the segment. While it does not affect the optimization outcome—since the total length of the time series is fixed across segmentations—it is conventionally included for mathematical completeness. In our implementation, this term has no impact on model selection or the detection of change-points.

The bandwidth parameter of the Gaussian kernel, ν, was set using the commonly applied median heuristic [111], based on a subset *x*_*selected*_ of 10k segments:

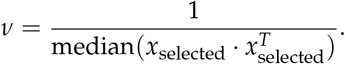

Additionally, the minimum duration of the segments was set to 10 IMU samples (i.e., 33.3 ms for rat recordings), corresponding to one video frame to prevent spurious, very short segments from being detected by the algorithm.

#### Implementation

The implementation of PELT available in the ruptures package [104] was modified in order to perform the change-point detection pipeline on 1 min blocks independently. This modification allowed parallel change points computation on the different sub-time series and limited the space complexity of the computation (𝒪(*n*^2^) for the implementation in ruptures due to the computation of a full Gram matrix). Although loosening the guaranty of obtaining an optimal segmentation, our approximation had minimal impact on the output (Fig. S2C).

### Feature extraction

#### Input data

After obtaining the estimated change-points in IMU data 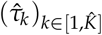, we obtain for each segment *k* among the 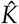:

- 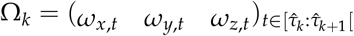: angular speed measurement (^°^/s).
- 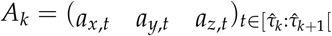: acceleration measurement (*g*).

Writing *sr* the sampling rate, we also obtain:

- 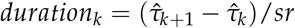: duration of the segment (s).

IMU data preprocessing (see Methods) yielded for each resulting segment:

- 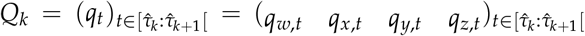: quaternion representing the orientation of the sensor relative to an arbitrary, gravity-polarized Earth reference-frame.
- 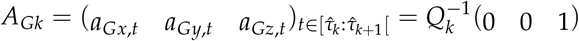: estimate of the gravitational component of acceleration in the head reference frame, or head tilt.
- 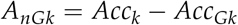: estimate of the non-gravitational component of acceleration in the head reference frame.

#### Per-segment spherical heading trajectories

Unit quaternions computed using an extended Kalman filter (see Methods) represent sensor orientation relative to an arbitrary, gravitationally polarized Earth reference frame and can therefore be considered as an estimate the head’s absolute orientation. While the tilt component of this estimate is highly reliable, as previously demonstrated [42], the azimuthal component (i.e., head direction in the Earth-horizontal plane) tends to drift due gyroscope noise integration. Although magnetometer readings can be used to compensate for this drift, we chose to ignore them due to an inadequate magnetic environment. Nevertheless, magnetometer-free estimates of absolute head orientation can be used for short-duration segments, where azimuthal drift is minimal, particularly during rapid head movements characterized by a high gyroscope signal-to-noise ratio.

For each segment *k*, an estimate of the animal’s absolute heading (corresponding to a spherical trajectory on the unit sphere) was computed by applying quaternions to the unit vector (1 0 0) followed by a rotation about the Earth-vertical axis to align its first sample with an azimuth of zero:

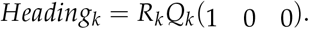

To obtain the rotation matrix *R*_*k*_ which performs the azimuthal alignment, the absolute orientation for the first sample of segment *k* is first computed:

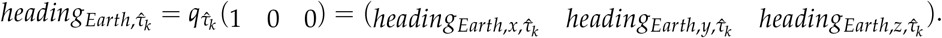

From this initial absolute orientation, we then derive the initial azimuth 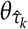 for segment *k*:

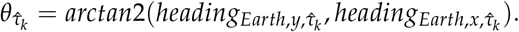

Lastly, we define *R*_*k*_ as a rotation by an angle 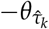 about the Earth-vertical axis:

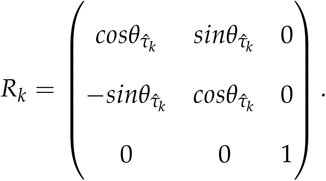

Aligning individual heading trajectories on the same initial azimuth allows superimposing them on the same graph to visualize head dynamics for multiple segments (see for example Fig. 2D_3_).

#### General statistics on inertial time series

For each segment *k*, the mean, standard deviation, minimum, maximum, median and quartiles were computed for each axis of Ω_*k*_, *A*_*Gk*_ and *A*_*nGk*_, as well as for their estimated gradients and the *L*_2_ norm of Ω_*k*_ and *A*_*nGk*_. This resulted in a total of 140 features.

We also calculated segment duration and its logarithm (base 10), adding 2 more features. All pairwise correlations between the axes of Ω_*k*_ and *A*_*nGk*_ were computed, resulting in an additional 15 features. Furthermore, the energy of gyroscope measurements and non-gravitational acceleration for each segment were included, adding 2 more features.

Additionally, we computed the number of zero-crossings in Ω_*k*_, *A*_*Gk*_ and *A*_*nGk*_, resulting in another 9 features.

#### Gyroscope measurements in a gravitationally polarized reference frame

Using our estimate of the head’s absolute orientation, the coordinates of head angular velocity (measured by gyroscopes in the head reference frame) can be computed in an external, gravitationally polarized reference frame:

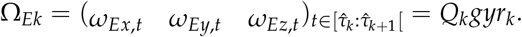

For each segment *k*, the component 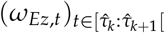 corresponds to the Earth-vertical (azimuthal) component of head angular velocity, i.e. its angular speed about the axis of gravity. This component, previously used to quantify circling behavior [36, 42], is here used to derive 9 features of particular interest for characterizing changes in head orientation:

- The mean, standard-deviation, minimum, maximum, median and quartiles of the cumulative change of azimuth over the segment, i.e. the cumulative sum of 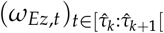 divided by the sampling rate.
- The total cumulative change of azimuth (net azimuthal change):

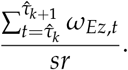
- The net azimuthal speed:

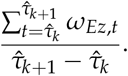

#### Head tilt and 3D heading estimate

For each segment *k*, gravitational acceleration in the head reference frame 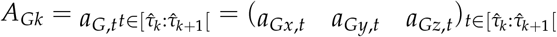 was used to compute a total of 11 features:

- The coordinates of the initial (first sample) gravitational acceleration vector 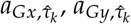 and 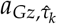
- The coordinates of the final (last sample) gravitational acceleration vector 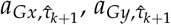 and 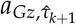
- The total change in the coordinates of the gravitational acceleration vector: 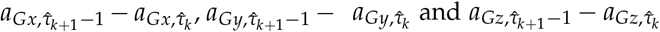
- The total change in head tilt (in degrees):

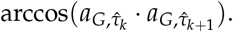
- The net head tilt change speed (in ^°^/s):

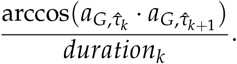

Similarly, the total change in 3D heading and net 3D heading change speed were computed from the first and last samples of the 3D heading estimate, totaling 2 features.

#### Mean orientation and dispersion of head tilt and 3D heading

Because head tilt and 3D heading follow trajectories on the unit sphere S ^2^, their mean orientation and dispersion were estimated by approximating the mean and concentration of a von Mises-Fisher distribution [112]. Dispersion was calculated as the invert square-root of the concentration. The resulting 8 features corresponded to the 3D coordinates of the mean orientation and the dispersion, for both head tilt and 3D heading.

#### Time-frequency features

The continuous wavelet transform of each axis of the angular velocity and non-gravitational acceleration time series was computed using the R package WaveletComp, with Morlet wavelets at dyadically spaced frequencies between 1 and 20 Hz (20 sub-octaves). The magnitude of wavelet coefficients were averaged into 7 linearly spaced bands between 2.5 and 20 Hz. For each segment, the base-10 logarithm of the median coefficients in these bands was taken, resulting in 42 additional features (7 frequency bands per axis).

#### Automatic dimensionality reduction

Principal component analysis (PCA) remains the most commonly used dimensionality reduction technique [113]. Contrary to variable selection approaches which focus on reducing the number of input variables of the original dataset, PCA builds a linear transformation of the data matrix to obtain a new data matrix characterized by less variables. By design, the new variables are all dependent on the original variables through this linear transformation. Introduced by Pearson[114] and rediscovered by Hotelling[115], PCA has been applied in numerous scientific fields across various applications. It can be described using geometrical properties or within an optimization framework. In the latter, considering *X* ∈ ℳ _*N*×*p*_(ℝ) as the original matrix of *N* observations (rows) and *p* quantitative variables (columns), and the linear transformation *Y* = *XU* with *U* ∈ ℳ _*p*×*d*_(ℝ) and *d* ≤ *p*, PCA can be framed through the following maximization problem:

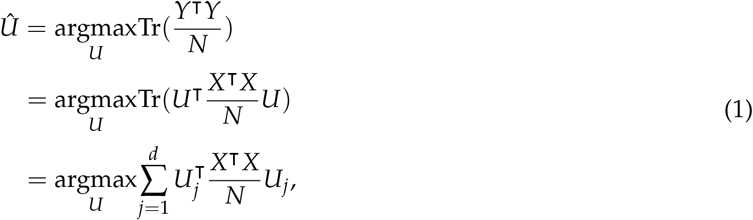

under the constraint that *U*^⊺^*U* = *I*_*d*_. In Equation (1), Tr denotes the trace operator which sums the diagonal elements of a square matrix and *U*_*j*_ ∈ ℝ ^*p*^ is column *j* of matrix *U*. By construction, if the dataset in *X* is centered (if not, a centering operation is employed), *X*^⊺^*X*/*N* = *C*_*X*_ is the empirical covariance matrix associated with the data in the original matrix *X*. To limit the influence of variables with large variances in *X* on the construction of *U* in PCA, the variables in *X* are almost always scaled so that *C*_*X*_ becomes the empirical correlation matrix. Using spectral theory arguments, it can be shown that the optimal solution to this problem is obtained through an eigen decomposition of *C*_*X*_ [116]. Thus, column *Û* _1_ of matrix *Û* is set to the eigen vector of *C*_*X*_ associated with the largest eigenvalue *λ*_1_, *Û* _2_ to the one with the second largest, and so on up to *Û* _*d*_. We emphasize that since *X* is centered and *Y* = *XÛ*, so is *Y*. Therefore, *Y*^⊺^*Y*/*N* = *C*_*Y*_ = *Û* ^⊺^*C*_*X*_*Û* = diag(*λ*_1_, *λ*_2_, …, *λ*_*d*_) is the empirical covariance matrix with entries *λ*_*j*_ on the diagonal and 0 elsewhere. So, the new *d* variables created in matrix *Y* have no correlation. Moreover, since the goal is to maximize Tr(*Y*^⊺^*Y*/*N*) = Tr(*C*_*Y*_) in Equation (1), it can emphasized that the key objective at the core of PCA is to build a new dataset *Y* ∈ ℳ _*N*×*d*_(ℝ) from *X* with less variables so that the new observations in *Y* have maximal spread (maximal variance), with uncorrelated variables. Besides, at the optimal matrix *Û*, the objective function becomes:

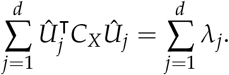

If *d* is set to *p*, corresponding to the case where all dimensions are preserved, then

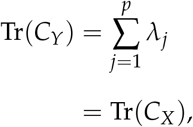

so that the sum of the empirical variances in the original dataset *X* are perfectly preserved in the new dataset *Y*. In practice, *d* is chosen to be small compared to *p* and the percentage of variance of *X* kept in *Y* is 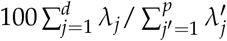.

In spite of the popularity of PCA, no authoritative solution has been widely accepted for choosing the number *d* of eigen vectors -called principal components (PCs)- in *U* to build *Y*. Most existing applications choose the dimension *d* by considering the eigenvalues scree of *C*_*X*_. This ad-hoc technique, popularized by Cattell[117], has been largely modified and perfected over the last 50 years [118]. Recently, focusing on the probabilistic version of PCA [119], an analytical expression of the marginal likelihood was obtained, allowing *d* to be estimated [118, 120]. In this paper, we rely on the cross validation approximation strategy of Josse and Husson[106] which scales well with larger data matrices.

#### Implementation

We used the package *scikit-learn* in Python to perform PCA with the empirical correlation matrix of the data. The number of dimension *d* was estimated using the cross validation approximation strategy of Josse and Husson[106], using the function *estim_ncp* from the package *FactoMineR* in R.

#### Clustering

In this paper, we relied on the expectation maximization algorithm (EM[76]) for GMMs which is widely seen as a reference for clustering [121]. Indeed, contrary to kmeans [122] for instance, which assumes the clusters of observations to be characterized by spheres in the space of variables, fewer assumptions are made in GMMs such that clusters with different shapes can be uncovered. In particular, GMMs allow both spheres as well as ellipsoids with different volumes, shapes, and directions, to be handled. Moreover, contrary to kmeans, no assumption is made on the average size that the clusters of observations should have [73]. Finally, it is a soft clustering approach based on a probabilistic model. As such, it provides statistics, which strongly help interpreting the clusters found, in practice, and it gives access to theoretically validated mathematical tools from computational statistics [75]. In particular, contrary to most clustering approaches which rely on heuristics to estimate the number of clusters present in the data, model selection criteria with theoretical guaranties can be used [123].

Denoting *X* ∈ ℳ _*N*×*p*_(ℝ) the matrix of *N* observations and *p* quantitative variables with row *i, x*_*i*_ ∈ ℝ ^*p*^, the GMM model considers the following mixture density function with *K* components:

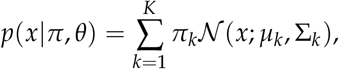

where 𝒩 (*x*; *µ*_*k*_, Σ_*k*_) is the density function of a multidimensional Gaussian law with mean *µ*_*k*_ and covariance matrix Σ_*k*_ evaluated at *x*. We note *θ* = {(*µ*_1_, Σ_1_, …, (*µ*_*K*_, Σ_*K*_} the set of all parameters controlling the Gaussian density functions. Moreover, *π* ∈ [0, 1]^*K*^ is a vector of mixing weights which are probabilities such that 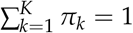. The log likelihood of the model associated with the observations in *X* is given by:

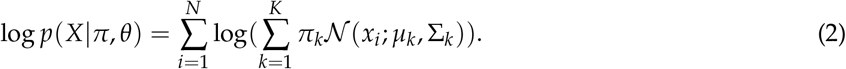

Unfortunately, maximizing Equation (2) does not lead to analytical expressions for *π* and *θ*. The EM algorithm [76] was introduced in 1977 to tackle this issue and has been widely used as a solution to this problem this then.

#### EM procedure

The EM algorithm for fitting a GMM involves two main steps as well as an initialization.

#### Initialization

The EM algorithm is initialized with a kmeans algorithm with *K* clusters applied on *X* such that *τ*_*ik*_ is set to 1 if observation *i* was found in cluster *k*, 0 otherwise.

- **E Step:** Calculate the expected posterior values *τ*_*ik*_ that observation *i* is actually from cluster *k*, using the current estimates of the model parameters (*π, θ*)
- **M Step:** Calculate the estimators for *π* and *θ*, using the current estimate of the *τ*_*ik*_

The E and M steps are repeated in a loop which maximizes the log likelihood of the data. The loop is stopped when (2) converges.

**E-Step** Calculate

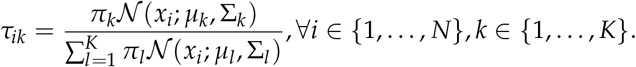

**M-Step** Calculate

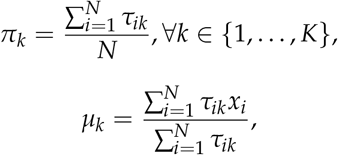

and

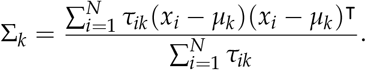

#### Model selection

In this paper, we relied on the BIC criterion for GMM[121] to estimate the number of clusters present in the data matrix *X*. It is given by:

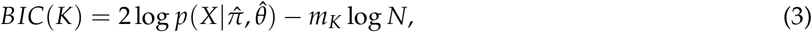

where 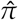 and 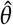 are the estimators found by the EM algorithm for *K* clusters, and *m*_*K*_ is the total number of free parameters. For a full GMM model with no constraint on the covariance matrices Σ_*k*_, *m*_*K*_ is given by:

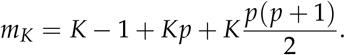

In practice, we considered a grid from 5 to 100 for *K*, and for each value, a full EM algorithm was employed to compute *BIC*(*K*. Moreover, several types of covariance structures (full, diagonal, tied, spherical) corresponding to assumptions on the shapes of the clusters were tested. Finally, the model retained was the one for which the number of clusters and the type of covariance structure were maximizing the BIC criterion over the grid search.

#### Implementation

We used the package *scikit-learn* in Python to perform BIC and EM for GMM on the data matrix. Since EM is only guarantied to reach a local maximum of the log likelihood (2), we repeated the all procedure 10 times for multiple initialization.

#### Sequence analysis using a Hidden Markov Model with categorical emissions

Following the application of DISSeCT to the inertial data, we obtained *N* segments, each assigned to one of *K* − 1 clusters identified by the GMM with the highest posterior probability, alongside an additional label for outliers. This process resulted in a symbolic sequence representing the clusters.

To capture the temporal dynamics of this sequence, we employed a categorical hidden Markov model (HMM) with S hidden states, utilizing the [hmmlearn](https://hmmlearn.readthedocs.io/en/latest/api.html#hmmlearn.hmm.CategoricalHMM) library. The model parameters, including transition and emission probabilities, were estimated using the EM algorithm, as detailed in the following sections.

#### Model of the symbolic sequence

The symbolic sequence of *K* possible clusters associated with each of the *N* segments, denoted by *Y*, was modeled as resulting from a sequence of hidden states *S*. The HMM used to represent this sequence is defined as follows:

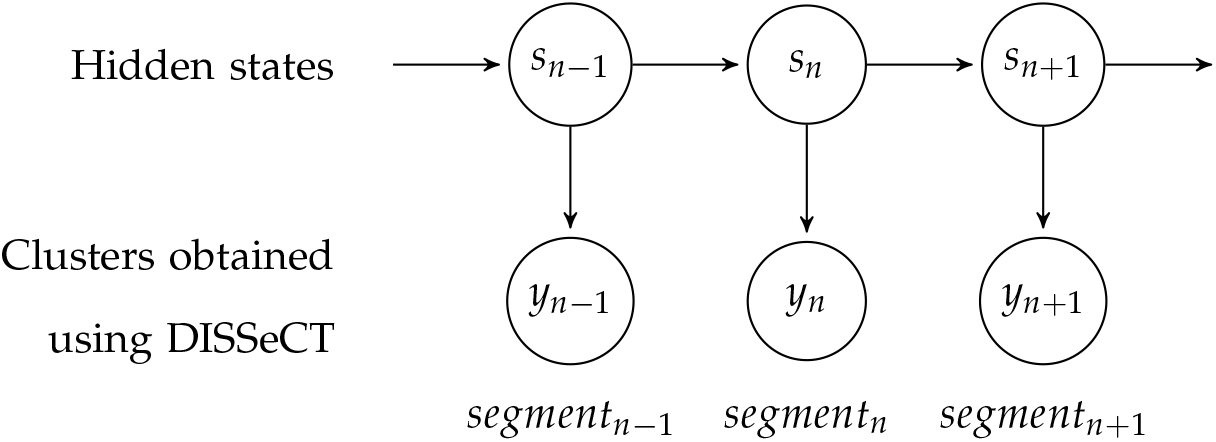

HMM graphical model of the symbolic sequence of clusters associated with individual segments.

In the following, we denote:

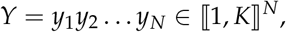

and

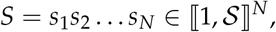

where *Y* is the observed sequence of clusters, and *S* is the corresponding sequence of hidden states.

##### Model parameters

The model parameters encompass:

- **Initial state distribution** *π* = (*π*_*i*_)_*i*∈ ⟦1, 𝒮⟧_:

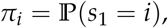
- **Transition matrix** 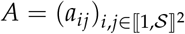:

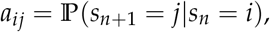
- **Emission matrix** *E* = (*e*_*ik*_)_*i*∈ ⟦1, 𝒮⟧,k∈ ⟦1,K⟧_:

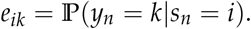

Under the categorical HMM, the likelihood of the observed data *Y*, given the model parameters *θ* = (*π, A, E*, is expressed as:

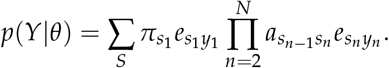

This likelihood represents the probability of observing the sequence *Y* by summing over all possible sequences of hidden states *S*.

#### Priors and initialization

We initialize the parameters of the model *θ* = (*π, A, E*) as follows:

- **Initial state distribution** *π*: The initial state distribution *π* is initialized using a Dirichlet prior distribution. Thus, each state *i* ∈ {1, …, 𝒮} is assigned an initial probability drawn from a Dirichlet distribution with a parameter vector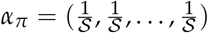:

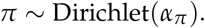
- **Transition matrix** *A*: The transition probabilities *A* are also initialized using a Dirichlet prior distribution. Thus, for each state *i* ∈ {1, …, 𝒮, the transition probabilities to all other states are drawn from a Dirichlet distribution with parameter vector 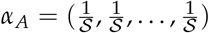:

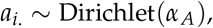

where *a*_*i*._ denotes row *i* of matrix *A* and is the set of all transition probabilities from state *i*.
- **Emission matrix** *E*: The emission probabilities *E* are initialized uniformly. Thus, each element *e*_*ik*_ is first drawn from a continuous uniform distribution between 0 and 1:

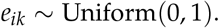

Then, the rows are normalized to ensure they sum to 1. Thus, for each state *i* ∈ {1, …, 𝒮}:

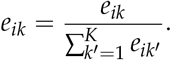

#### EM procedure

The EM algorithm for fitting a categorical HMM involves two main steps:

- **E Step:** Calculate the expected value of the complete log likelihood function, using the current estimates of the model parameters.
- **M Step:** Maximize this expected complete log likelihood function to update the model parameters.

The procedure was repeated for 100 iterations to ensure that the model parameters are optimized to best fit the observed data *Y*.

**E-Step** In the E step, we utilize the Viterbi filter to determine the most likely sequence of hidden states, given the observed data *Y* and the current estimates of the parameters.

1. **Initialization:** Initialize the Viterbi variables and the back-pointer array *ψ*:

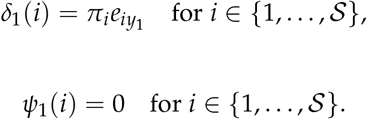
2. **Recursion:** For each time step *n* = 2, …, *N* and for each state *j* ∈ {1, …, 𝒮}:

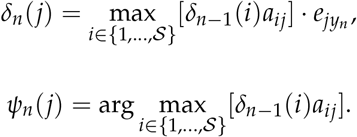
3. **Termination:** Find the highest probability of the final state:

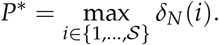
4. **Path Backtracking:** Backtrack through the computed variables to find the most probable sequence of hidden states. Start with the state that has the highest probability at the final time step:

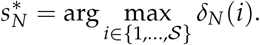

For *n* = *N* − 1, *N* − 2, …, 1, determine the preceding state using the back-pointers, which track the state transitions:

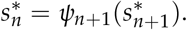

Here, *ψ* is an array of back-pointers that stores the indices of the states that led to the maximum probability at each time step, facilitating the reconstruction of the most likely state sequence.

**M-Step** In the M-step, we update the model parameters *θ* = (*π, A, E*) based on the state sequence 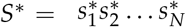 obtained from the E-step.

1. **Update initial probabilities:**

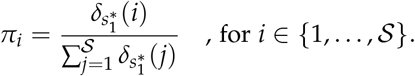
2. **Update transition probabilities:**

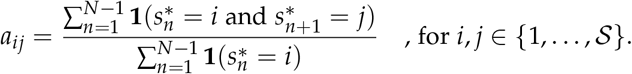
3. **Update emission probabilities:**

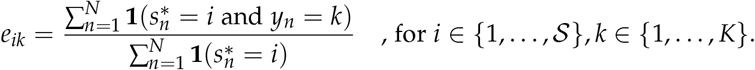

#### Hyperparameter optimization

Hyperparameter optimization was performed through maximization of the BIC. The BIC of the categorical HMM is defined as:

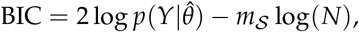

where 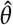 is obtained with the EM algorithm described above, *m*_𝒮_ is the number of parameters, and *N* is the number of observations. For each candidate number of hidden states 𝒮, ranging from 1 to 30, ten independent model fits were conducted to capture variability in the results. The model yielding the highest BIC score was then selected as the optimal configuration for the HMM.

#### 3D pose estimation

##### Parameters for 2D pose estimat

The parameters for 2D pose estimation using DeepLabCut are summarized in Table S1.

##### Frame los

To ensure that 2D pose estimates obtained from the five different cameras were properly synced despite occasional frame loss (approximately 0.02 % of the frames), frame time-stamping was used to estimate the exact timing of frame loss for a given camera. Pose estimates for these intermittently lost frames were computed using cubic interpolation.

##### Parameters for 3D pose estimation

The parameters for triangulating 2D pose estimates and post-processing of the resulting 3D pose are summarized in Table S2.

##### Obtaining a gravity-polarized Earth reference-frame

The Earth reference frame resulting from Anipose is oriented relative to one of the cameras. To rotate this reference frame so that its *z*-axis is aligned with the Earth-vertical axis, we computed the principal components of the estimated 3D snout trajectory and used the first two principal components as a proxy for the arena’s floor plane (Fig. S1A). After confirming that we obtained a right-handed reference frame oriented upward, we also translated this reference frame to obtain trajectories with positive values (minimum altitude and *x*/*y* values set to 0) (Fig. S1B). Visual inspection of the resulting pose estimates was performed to confirm the correct orientation of the reference frame (Fig. S1C,D).

## Supplementary Tables

**Table S1:** Summary of the parameters used for obtaining 2D pose estimates with DeepLabCut. The analysis was run on a Dell Precision 5820 Tower equipped with an NVIDIA Quadro RTX 6000 (24 GB memory).

| DeepLabCut |  |
| --- | --- |
| Version | 2.2.0.3 |
| Deep neural network | resnet 101 [124] |
| Training set | 1 950 frames with 12 anatomical landmarks |
| Data augmentation | Default DLC augmentation |
| Training iterations | 300 000 |
| Learning rate | 0.005 for iterations 0–10k then 0.02 |

**Table S2:**
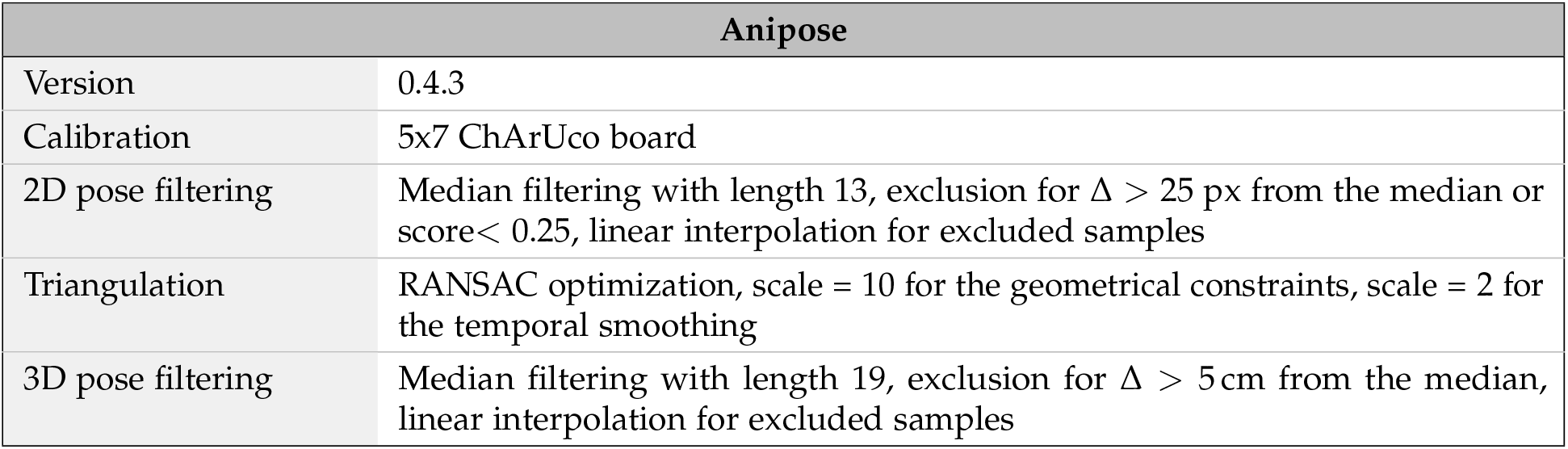
Summary of the parameters used for obtaining 3D pose estimates using Anipose. The analysis was run on a Dell Precision 5820 Tower equipped with an NVIDIA Quadro RTX 6000 (24 GB memory).

| Anipose |  |
| --- | --- |
| Version | 0.4.3 |
| Calibration | 5x7 ChArUco board |
| 2D pose filtering | Median filtering with length 13, exclusion for $\Delta > 25$ px from the median or score $< 0.25$, linear interpolation for excluded samples |
| Triangulation | RANSAC optimization, scale = 10 for the geometrical constraints, scale = 2 for the temporal smoothing |
| 3D pose filtering | Median filtering with length 19, exclusion for $\Delta > 5$ cm from the median, linear interpolation for excluded samples |

**Table S3:** Summary of the parameters used for running Keypoint-MoSeq on our dataset. The analysis was run on a Dell Precision 5820 Tower equipped with an NVIDIA Quadro RTX 6000 (24 GB memory).

| Keypoint-MoSeq |  |
| --- | --- |
| Version | 0.5.2 |
| Number of keypoints | 10 (excluding tail) |
| Confidence threshold | All coordinate values were set to nan for confidence $< 0.6$ |
| PCA | 8 first PCs ($\geq 90$ % of variance explained) kept for model fitting |
| Kappa (stickiness) | 3e5 for the AR step and 3e4 for fitting the full model |
| Number of iterations | 50 for the AR step and 500 for fitting the full model |
| Memory usage | To keep GPU memory usage under 24 GB, the argument `parallel_message_passing=False` was passed to `kpms.fit_model()` |

**Table S4:** Computation time for the different steps of DISSeCT, measured on the same workstation used for running DeepLabCut, Anipose and Keypoint-MoSeq. Runtimes shown here were obtained for one run for the rat dataset comprising 8 hours and 38 minutes of inertial measurements (9.3M samples with a sampling rate of 30 Hz). The feature extraction, dimensionality reduction, clustering and UMAP embedding steps are performed for the 44k segments resulting from the change-point detection step. Results are provided both as the total execution time on the dataset, excluding data loading time, and per hour of recording.

| DISSeCT |  |
| --- | --- |
| Step | Computation time |
| Preprocessing and EKF filtering | 17 min 39 s (122 s/h) |
| Kernel change-point detection | 4 min 58 s (34.5 s/h) |
| Feature extraction | 3 min 52 s (26.8 s/h) |
| Dimensionality reduction and clustering | 52 s (6.0 s/h) |
| UMAP embedding | 1 min 27 s (10.1 s/h) |
| Total | 28 min 48 s (200 s/h) |

**Table S5:** Computation times for the different steps of Keypoint-MoSeq, measured on the same workstation used for running DeepLabCut, Anipose and DISSeCT, and excluding data loading/formatting, and PCA. Runtimes shown here were obtained for one run for the rat dataset comprising 8 h and 15 min of video recordings (891 052 frames recorded at 30 Hz). The number of iterations correspond to ones recommended by KpMS’s online documention (keypointmoseq.readthedocs.io). Results are provided both as the total execution time on the dataset, excluding data loading time, and per hour of recording.

| Keypoint-MoSeq (GPU with CUDA 12) |  |
| --- | --- |
| Step | Computation time |
| Model fitting: AR-HMM step (50 iterations) | 4 min 7 s (0.8 s/h/it) |
| Model fitting: full model (500 iterations) | 2 h 17 min 11 s (3.0 s/h/it) |
| Total | 2 h 21 min 18 s (1027.5 s/h) |
| Keypoint-MoSeq (CPU only) |  |
| Step | Computation time |
| Model fitting: AR-HMM step (50 iterations) | 10 min 11 s (1.5 s/h/it) |
| Model fitting: full model (500 iterations) | 1 d 2 h 9 min 51 s (22.8 s/h/it) |
| Total | 1 d 2 h 20 min 2 s (11 490.5 s/h) |

**Table S6:** Posthoc examination of individual clusters (rat dataset). For each cluster, the video snippets corresponding to the top 100 segments with the highest posterior probabilities of belonging to this cluster were examined, given a unique free label and linked with a unique main behavioral category among the following options: orienting, rearing, grooming, locomoting, idle or other (see Fig. 1C and Methods). The table shows the first and second most frequent labels per cluster, sorting clusters according to their associated dominant category. For each cluster, the right column shows the percentage of segments belonging to this associated dominant category. The second and third columns from the left indicate the fraction of segments assigned to each cluster (% Segment) and the fraction of the total recording time that they represent (% Duration).

| Cluster | Segment | Duration | Human labels (examination of top 100 segments) |  |  |  |  |
| --- | --- | --- | --- | --- | --- | --- | --- |
|  |  |  | Most frequent label | % | Second most frequent label | % |  |
| Locomotion |  |  |  |  |  |  | % locomotion |
| 46 | 3.5 | 6.1 | Locomotion | 100 | - | - | 100 |
| 37 | 3.3 | 9.8 | Locomotion with head down | 100 | - | - | 100 |
| 13 | 3.1 | 2.0 | Locomotion | 99 | End of rearing | 1 | 99 |
| 5 | 2.9 | 1.6 | Locomotion with downward pitch | 42 | Rearing end (forepaws touching floor) | 25 | 61 |
| Immobility |  |  |  |  |  |  | % immobile |
| 6 | 1.6 | 4.5 | Immobility | 100 | - | - | 100 |
| 38 | 1.4 | 9.3 | Immobility | 48 | Idle with small movements | 31 | 90 |
| 20 | 2.4 | 3.6 | Short bouts of immobility | 73 | Idle with small movements | 13 | 86 |
| Grooming |  |  |  |  |  |  | % grooming |
| 17 | 1.2 | 2.7 | Body licking left | 100 | - | - | 100 |
| 0 | 1.0 | 1.7 | Body licking right | 99 | Licking right forepaw | 1 | 100 |
| 43 | 2.7 | 4.9 | Nose/face wiping with frontpaws | 87 | Ear scratching with left hindpaw | 13 | 100 |
| 26 | 1.3 | 1.0 | Short bouts of body licking | 93 | Nose/face wiping with frontpaws | 6 | 99 |
| 19 | 1.0 | 0.2 | Scratching-associated reorientation | 48 | Body licking-associated reorientation | 28 | 95 |
| 30 | 1.4 | 1.6 | Licking left hindpaw | 94 | Investigating corner | 5 | 94 |
| 45 | 1.2 | 2.2 | Head/neck scratching with hindpaw | 89 | Reorientation (head down) | 8 | 89 |
| 32 | 1.2 | 0.3 | Body licking-associated reorientation | 65 | Nose/face wiping-associated reorientation | 22 | 87 |
| 33 | 1.5 | 0.3 | Body licking-associated reorientation | 58 | Nose/face wiping-associated reorientation | 20 | 79 |
| 3 | 1.1 | 0.9 | Licking left hindpaw | 68 | Investigating corner | 22 | 77 |
| 8 | 2.4 | 1.1 | Nose/face wiping with frontpaws | 62 | Sniffing head down | 31 | 69 |
| 29 | 0.7 | 0.1 | Scratching-associated reorientation | 51 | Nose/face wiping-associated reorientation | 14 | 68 |
| 25 | 1.5 | 6.2 | Nibbling fur | 36 | Investigating corner | 30 | 66 |
| 21 | 1.2 | 0.3 | Short bouts of body licking | 48 | Investigating corner | 36 | 54 |
| Rearing |  |  |  |  |  |  | % rearing |
| 9 | 2.9 | 6.3 | Rearing | 100 | - | - | 100 |
| 14 | 3.4 | 2.8 | Rearing | 100 | - | - | 100 |
| 41 | 2.3 | 0.7 | Rearing | 100 | - | - | 100 |
| 22 | 2.3 | 0.8 | Rearing | 99 | Rightward head turn, forepaws on floor | 1 | 99 |
| 28 | 2.9 | 1.0 | Rearing end, Leftward | 99 | Downward, leftward head rotation, forepaws on floor | 1 | 99 |
| 18 | 3.3 | 1.1 | Rearing end, rightward | 95 | Downward, rightward head rotation, forepaws on floor | 5 | 95 |
| 2 | 2.3 | 1.0 | Rearing initiation (forepaws leaving floor) | 56 | Upward head rotation (at least one forepaw not leaving floor) | 24 | 76 |
| 10 | 2.5 | 1.0 | Rearing initiation (forepaws leaving floor) | 66 | Upward head rotation (at least one forepaw not leaving floor) | 29 | 71 |
| 12 | 3.1 | 2.9 | Rearing initiation (forepaws leaving floor) | 49 | Upward head rotation (at least one forepaw not leaving floor) | 29 | 71 |
| 24 | 2.7 | 4.7 | Rearing initiation (forepaws leaving floor) | 51 | Head pitched up | 29 | 65 |
| 47 | 1.5 | 0.9 | Rearing | 50 | Investigating corner | 30 | 63 |
| 15 | 1.6 | 0.3 | Rearing initiation (forepaws leaving floor) | 36 | Upward head rotation (at least one forepaw not leaving floor) | 33 | 50 |
| Reorientation (mainly non rearing or grooming-related) |  |  |  |  |  |  | % reorientation |
| 4 | 3.2 | 1.7 | Upward head rotation | 100 | - | - | 100 |
| 1 | 2.8 | 0.9 | Leftward head turn | 99 | Leftward head turn during scratching | 1 | 99 |
| 40 | 2.4 | 2.7 | Leftward head turn, head pitched down | 84 | Rightward head turn, head pitched down, locomoting | 15 | 99 |
| 7 | 2.2 | 2.3 | Rightward head turn, head pitched down | 90 | Rightward head turn, head pitched down, locomoting | 8 | 98 |
| 16 | 2.7 | 2.1 | Small upward pitch rotation | 84 | Small upward pitch rotation with locomotion | 14 | 98 |
| 23 | 3.0 | 0.9 | Upward head rotation | 97 | Rearing | 2 | 97 |
| 42 | 2.4 | 0.7 | Rightward head turn | 93 | Rearing end, rightward | 7 | 93 |
| 27 | 2.6 | 0.6 | Leftward head turn | 84 | Rearing end, leftward | 14 | 84 |
| 44 | 2.2 | 0.7 | Rightward head turn | 80 | Grooming-associated reorientation | 20 | 80 |
| 11 | 2.5 | 1.6 | Leftward, downward head rotation | 66 | Rearing end, leftward | 25 | 75 |
| 34 | 1.8 | 0.4 | Leftward head turn | 62 | Grooming-associated reorientation | 21 | 62 |
| 39 | 1.5 | 0.4 | Rightward head turn | 61 | Grooming-associated reorientation | 21 | 61 |

## Supplementary Figures

**Figure S1:**
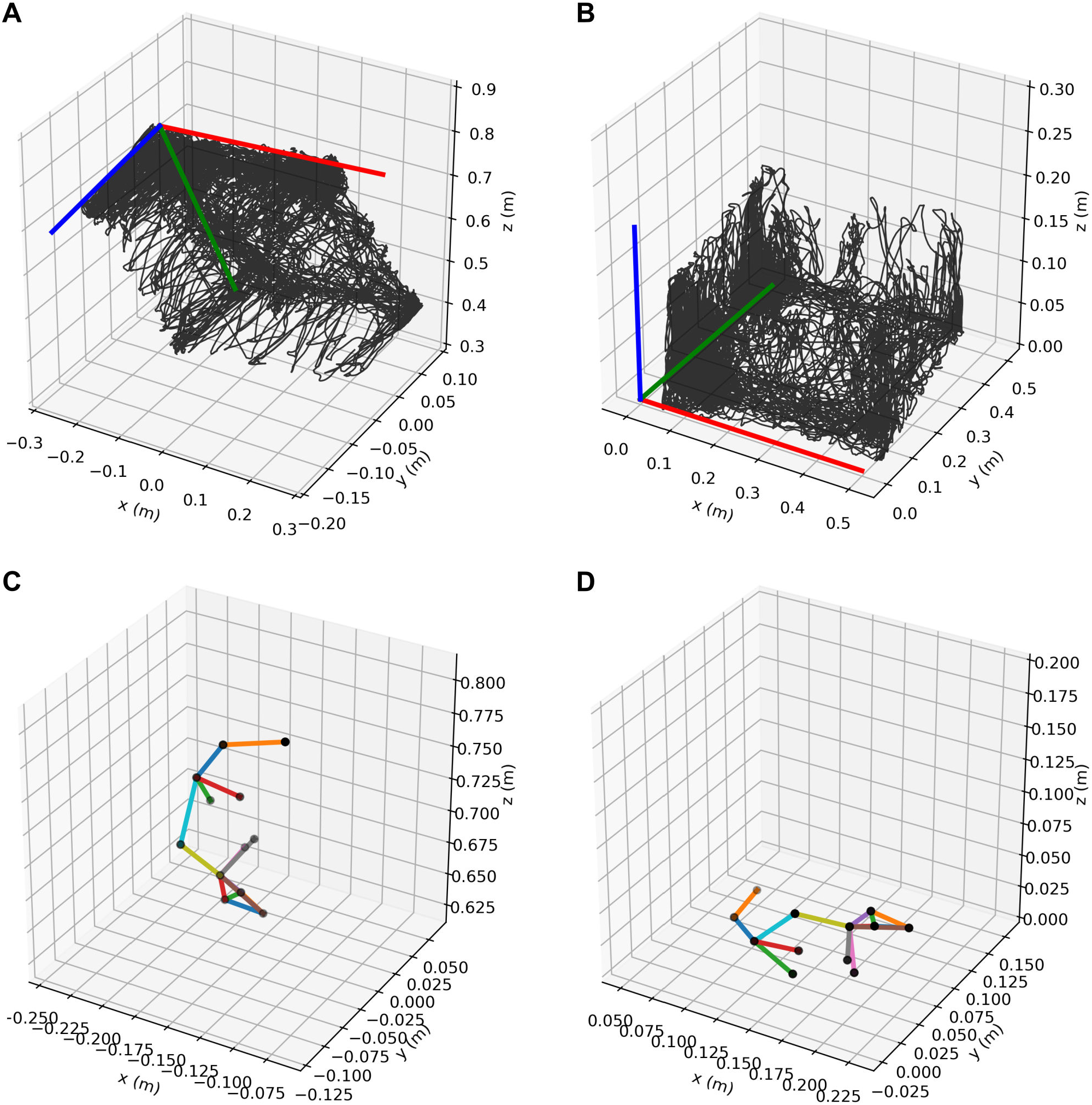
Method for obtaining a gravity-aligned reference frame for 3D keypoint tracking data. **A**, Snout trajectory (black) obtained using Anipose, shown with the orientations of its principal components: PC1 (red), PC2 (green), and PC3 (blue). **B**, The same trajectory after transformation into a gravity-aligned reference frame. **C**, Example 3D pose estimated using Anipose. **D**, The same pose after transformation into the gravity-aligned reference frame.

**Figure S2:**
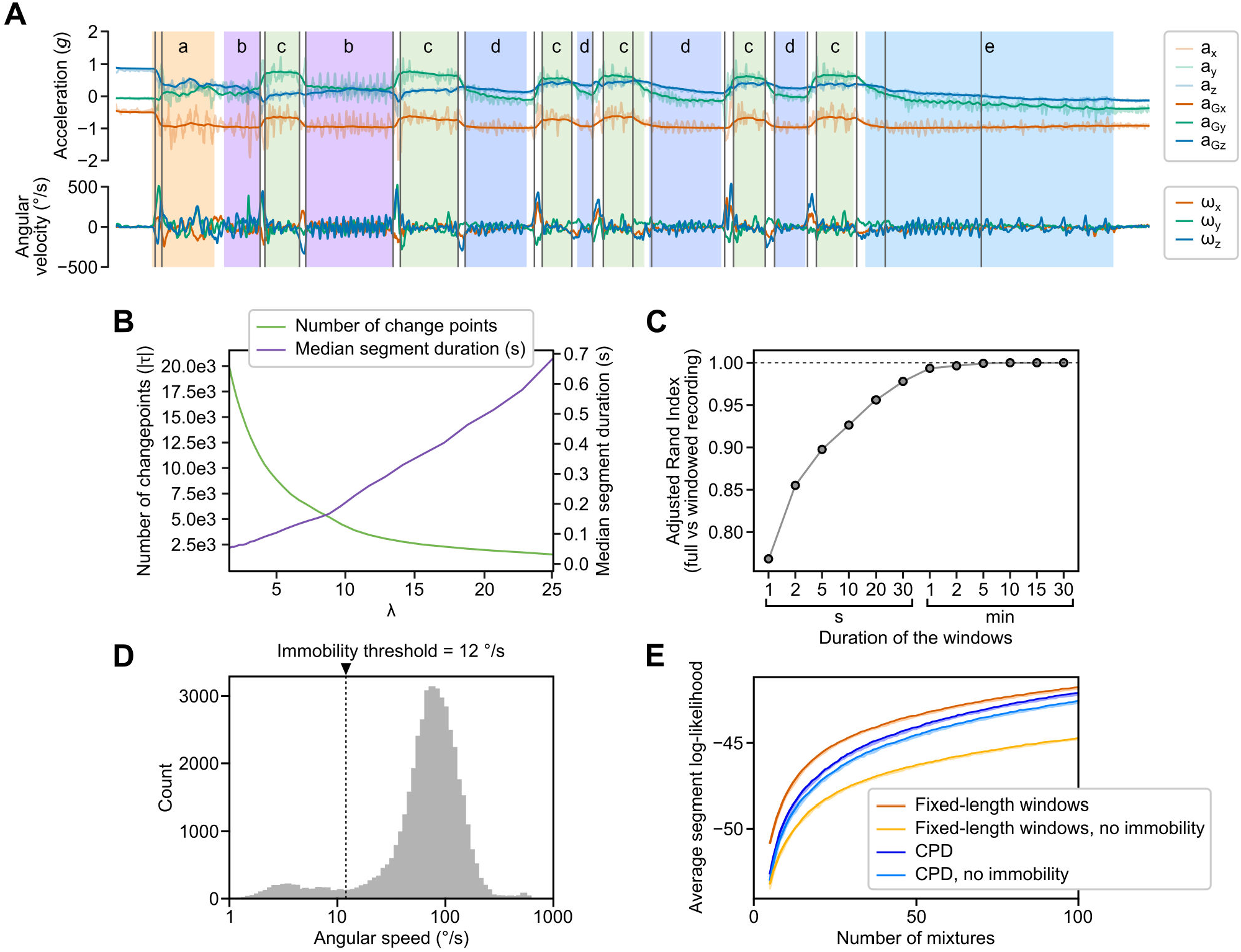
Justification and tuning of the change-point detection (CPD) method. **A**, Video-based, frame-level human annotations (colored areas) and detected change points (vertical gray lines) overlaid on IMU data from the 14-second scratching episode shown in Movie S1. Lowercase letters indicate the following behaviors: (a) sniffing with head pitched downward, (b) left body scratching, (c) left hind paw licking, (d) head scratching on the left side, and (e) decelerating head scratching on the left side. Change points were detected using Gaussian kernel-based Pruned Exact Linear Time (PELT) CPD applied to z-scored angular velocity (***ω***) and gravitational acceleration (**a**_**G**_) data. Total acceleration (**a**) is shown alongside **a**_**G**_ in lighter shades. **B**, Number of detected change points (green) and median segment duration (purple) as a function of the penalty parameter (*λ*), computed from z-scored **a**_**G**_ and ***ω*** data in a 30-minute example recording. **C**, Adjusted Rand Index (ARI) quantifying the similarity between segmentations obtained by running CPD on the full 30-minute recording versus running CPD in parallel on non-overlapping fixed-length windows. Notably, parallel CPD, which is computationally faster, produces nearly identical segmentations to the full-recording approach when window sizes are at least 5 minutes. **D**, Histogram of mean angular speed values (i.e., the norm of gyroscope measurements) computed over non-overlapping, fixed-length windows of 700 ms. The x-axis is logarithmic to highlight the bimodal distribution, which reflects periods of immobility and movement. A threshold of 12 ^°^ /s on the standard deviation of angular speed was applied to isolate immobility periods. **E**, Average log-likelihood (LL) computed across CPD segments or fixed-length windows from 10 independent GMM fits, plotted as a function of the number of mixture components *K*. LL values are shown with and without the ablation of immobility segments or windows prior to modeling (see main text for more details). For each *K*, the shaded region spans the full range of LL values across the 10 fits. Solid lines connect the maximum LL values at each *K*, indicating the best-fitting model.

**Figure S3:**
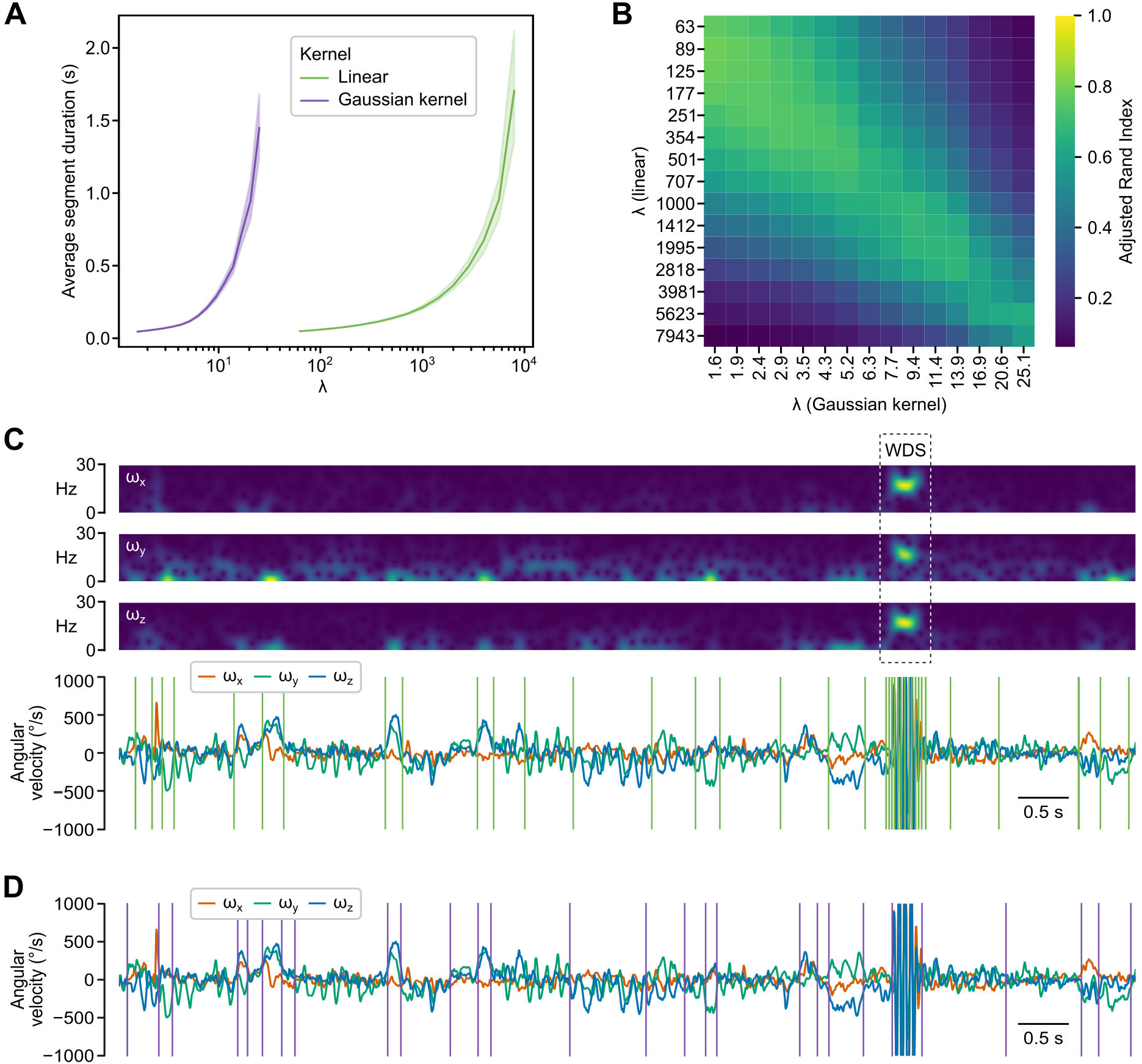
Comparison of linear and Gaussian kernel change-point detection (CPD) methods. **A**, Average segment duration as a function of the penalty parameter (*λ*) for a 5-minute recording. Results are shown for linear CPD applied to the z-scored time-frequency representation of gyroscope data (green) and for Gaussian kernel CPD directly applied to z-scored gyroscope data (purple). As CPD is deterministic, it was run once for each type of kernel. Shaded areas represent 95 % confidence intervals. **B**, Adjusted Rand Index comparing the segmentations obtained from the two methods in panel A. **C**, Example recording containing a wet dog shake (WDS) event. Top: Time-frequency representation of gyroscope signals, with the WDS’ spectral signature highlighted by a dashed rectangle. Bottom: Corresponding gyroscope signals, with segmentation results from linear CPD on their z-scored time-frequency representation shown as green vertical lines. The time-frequency representation was computed using a Short-Time Fourier Transform (STFT) with a DPSS window (Length = 1 s, Half-bandwidth = 125 ms). Frequencies above 30 Hz were excluded. **D**, Same recording as in C, with segmentation results from Gaussian kernel CPD on the z-scored gyroscope data shown as purple vertical lines.

**Figure S4:**
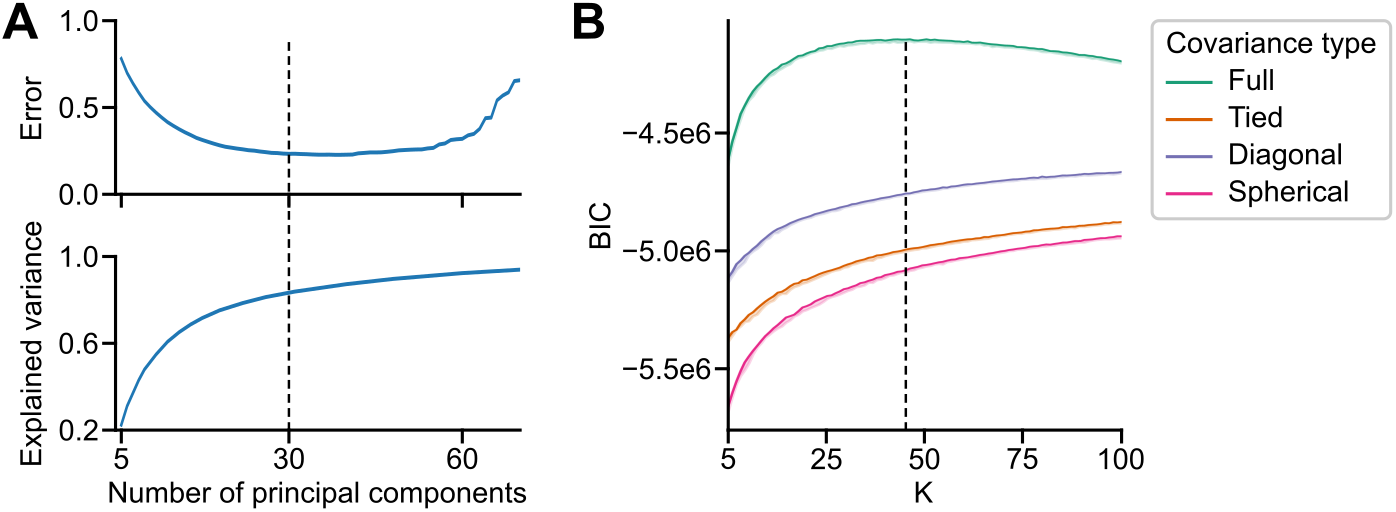
Hyperparameter selection for GMM clustering of segment-wise features. **A**, Selection of the number of principal components for dimensionality reduction, based on cross-validation error computed using the procedure described in [106]. The top panel shows cross-validation error as a function of the number of principal components; the bottom panel shows the corresponding fraction of explained variance. **B**, Bayesian Information Criterion (BIC) values for increasing numbers of mixture components *K* and different covariance matrix types (10 independent GMM fits per *K* value and covariance matrix type). For each *K*, the shaded region spans the full range of BIC values across the 10 fits. Solid lines connect the maximum BIC values at each *K*, indicating the best-fitting model.

**Figure S5:**
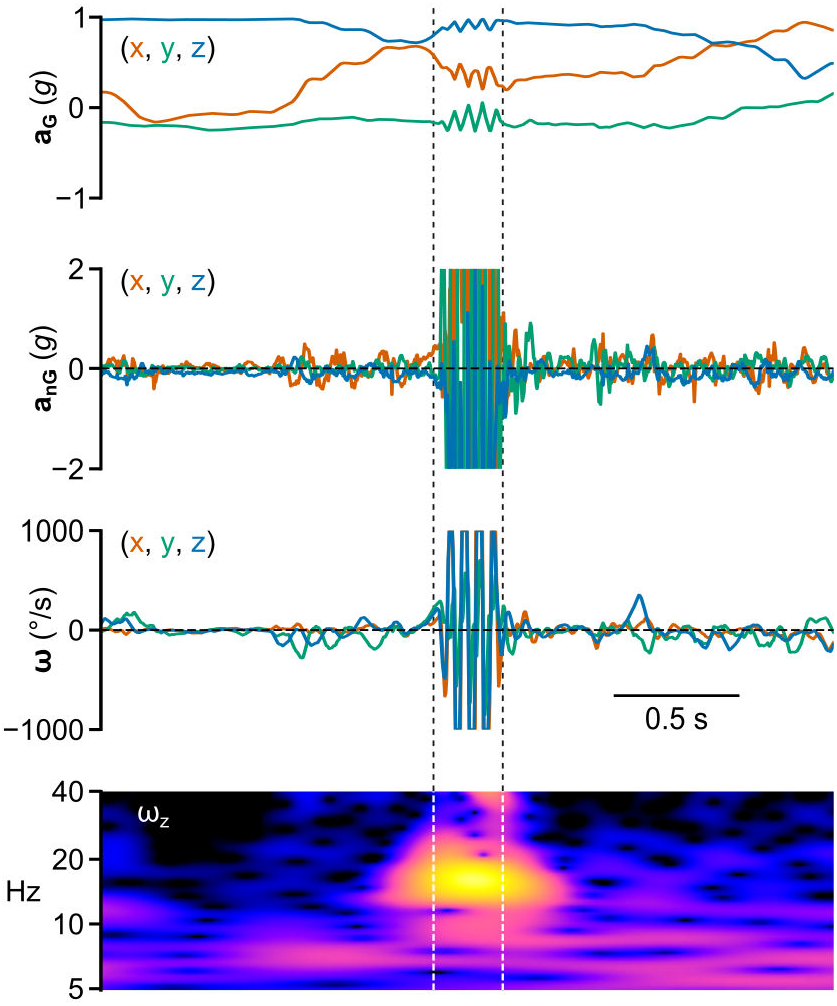
Inertial and spectral signatures of an example wet dog shake event. Traces of head gravitational acceleration (**a**_**G**_), non-gravitational acceleration (**a**_**nG**_) and angular velocity (***ω***) are shown. The bottom panel displays the log-magnitude of continuous wavelet transform coefficients computed on the yaw component of angular velocity. Vertical lines mark the start and end of the segment identified using Gaussian kernel-based PELT change-point detection (see Fig. S3).

**Figure S6:**
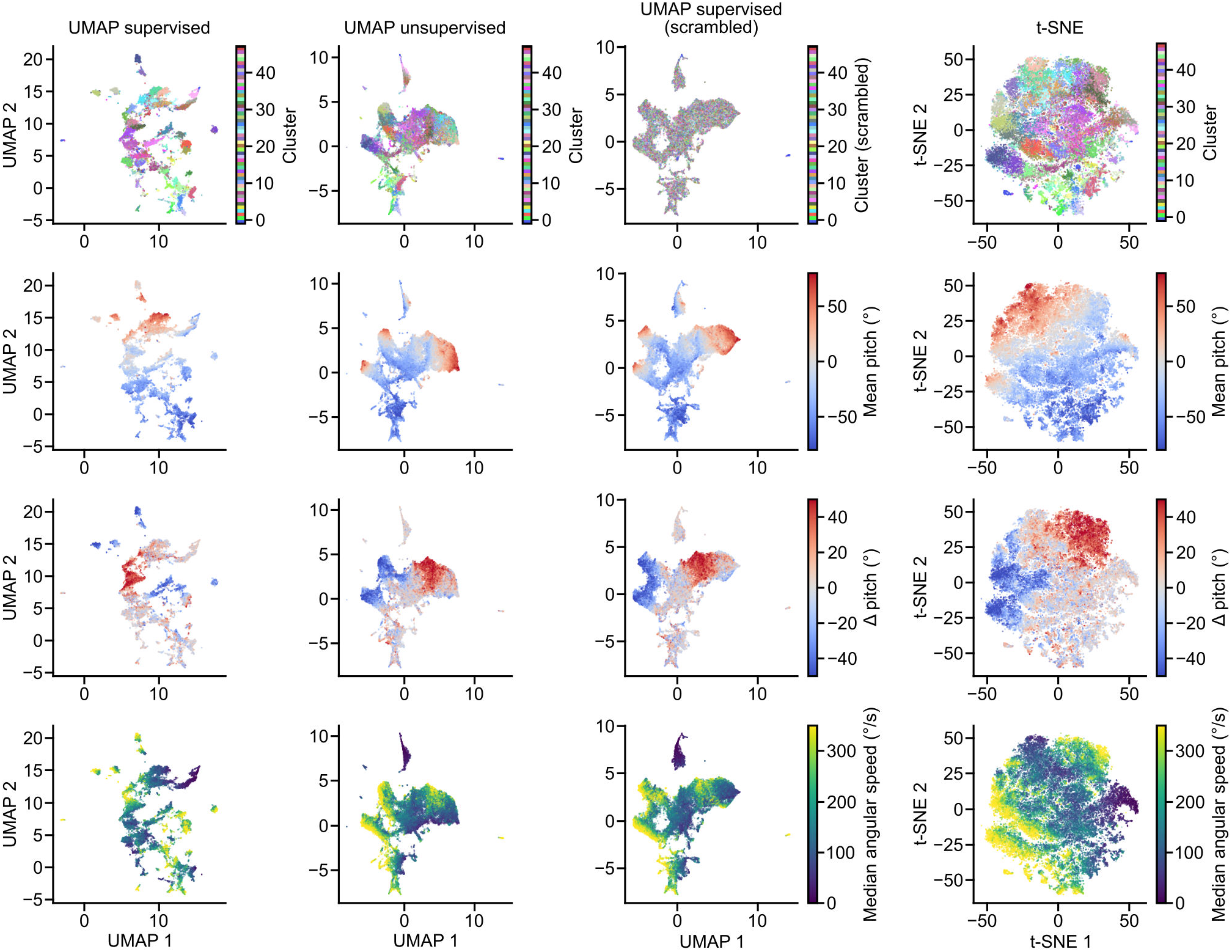
Comparison of 2D embedding methods for visualizing cluster distributions and segment-wise feature values in the feature space. Columns (left to right): (1) Supervised UMAP incorporating both feature values and cluster identities, (2) Unsupervised UMAP using only feature values, (3) Supervised UMAP with shuffled cluster identities, and (4) t-SNE projection based on feature values (early_exaggeration = 12, perplexity = 30). Top row: segments are colored according to their assigned cluster identity. Subsequent rows: segments are colored according to the values of selected example features.

**Figure S7:**
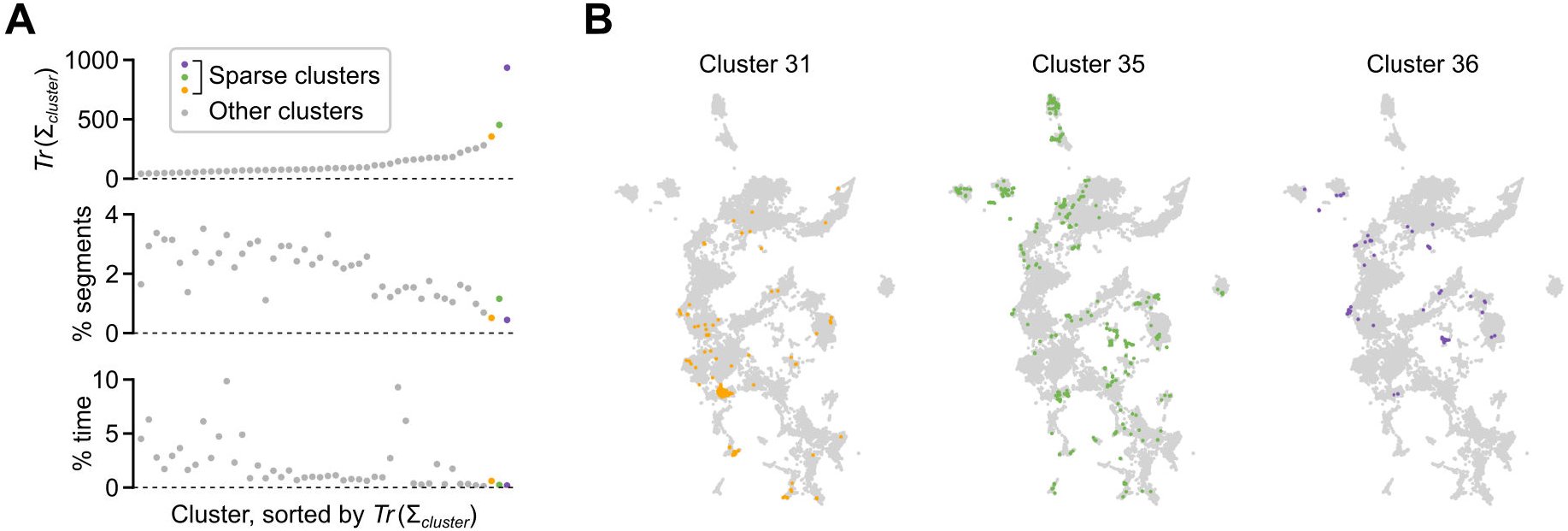
Identification of sparse clusters. **A**, Top: trace of the covariance matrix for each cluster, indicating cluster dispersion. The three highlighted clusters (in orange, green and purple) correspond to the most dispersed clusters. Middle: percentage of the total number of segments contained in each cluster. Bottom: percentage of the total recording duration associated to each cluster. Clusters are sorted by ascending trace values in all panels. **B**, Supervised UMAP embeddings displaying segments belonging to the three most dispersed clusters (in color) overlaid onto other segments (in gray).

**Figure S8:**
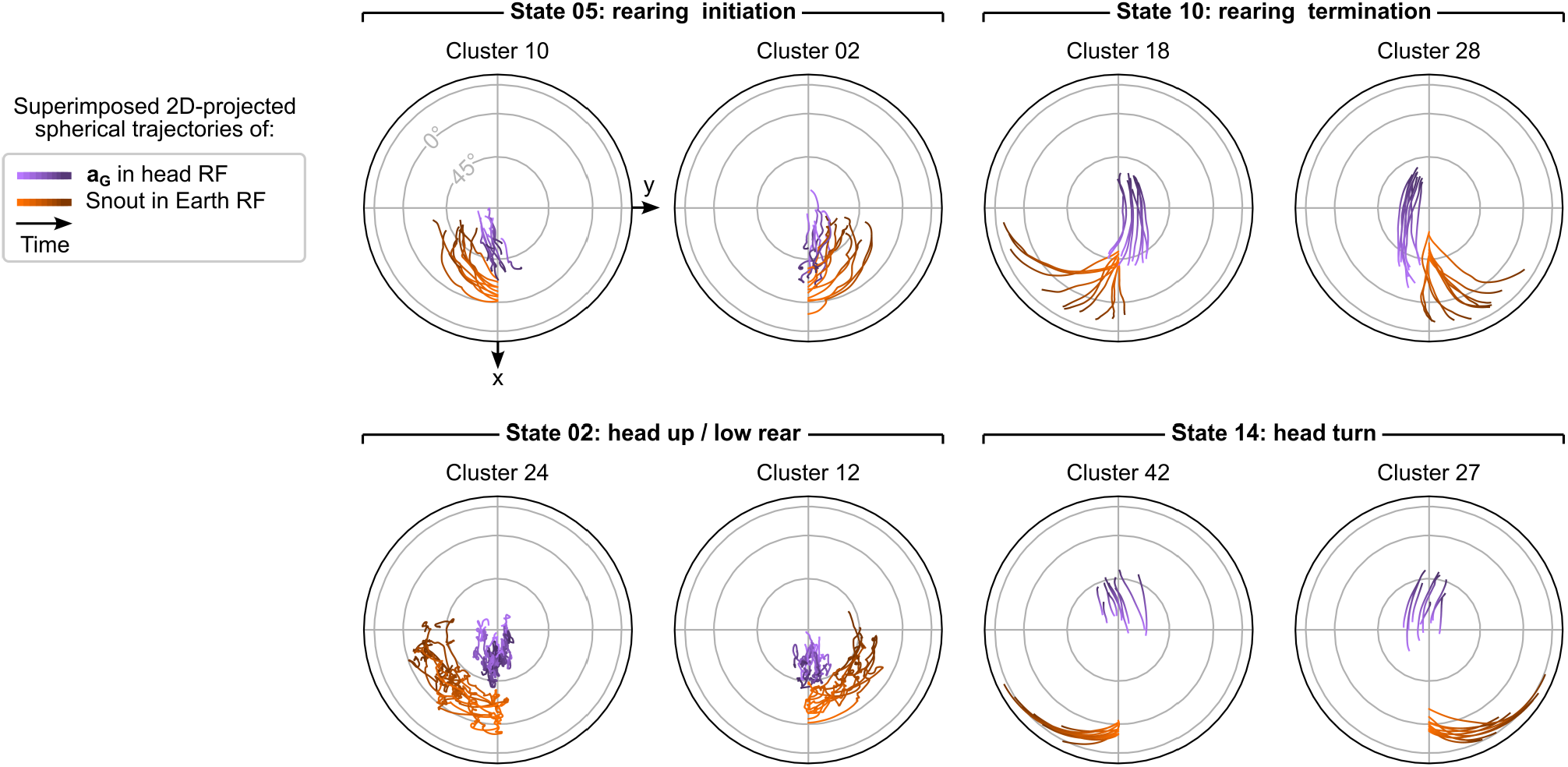
Grouping of similar leftward and rightward head movements by the categorical HMM. Plots represent the joint visualization of the trajectories of **a**_**G**_ in the head reference frame (purple) and of the animal’s heading in the Earth reference frame (orange) for the top 10 segments of pairs of clusters associated with four hidden states (see Supplementary Methods; RF: reference frame). Each pair represents the same head movement executed toward the left or right.

**Figure S9:**
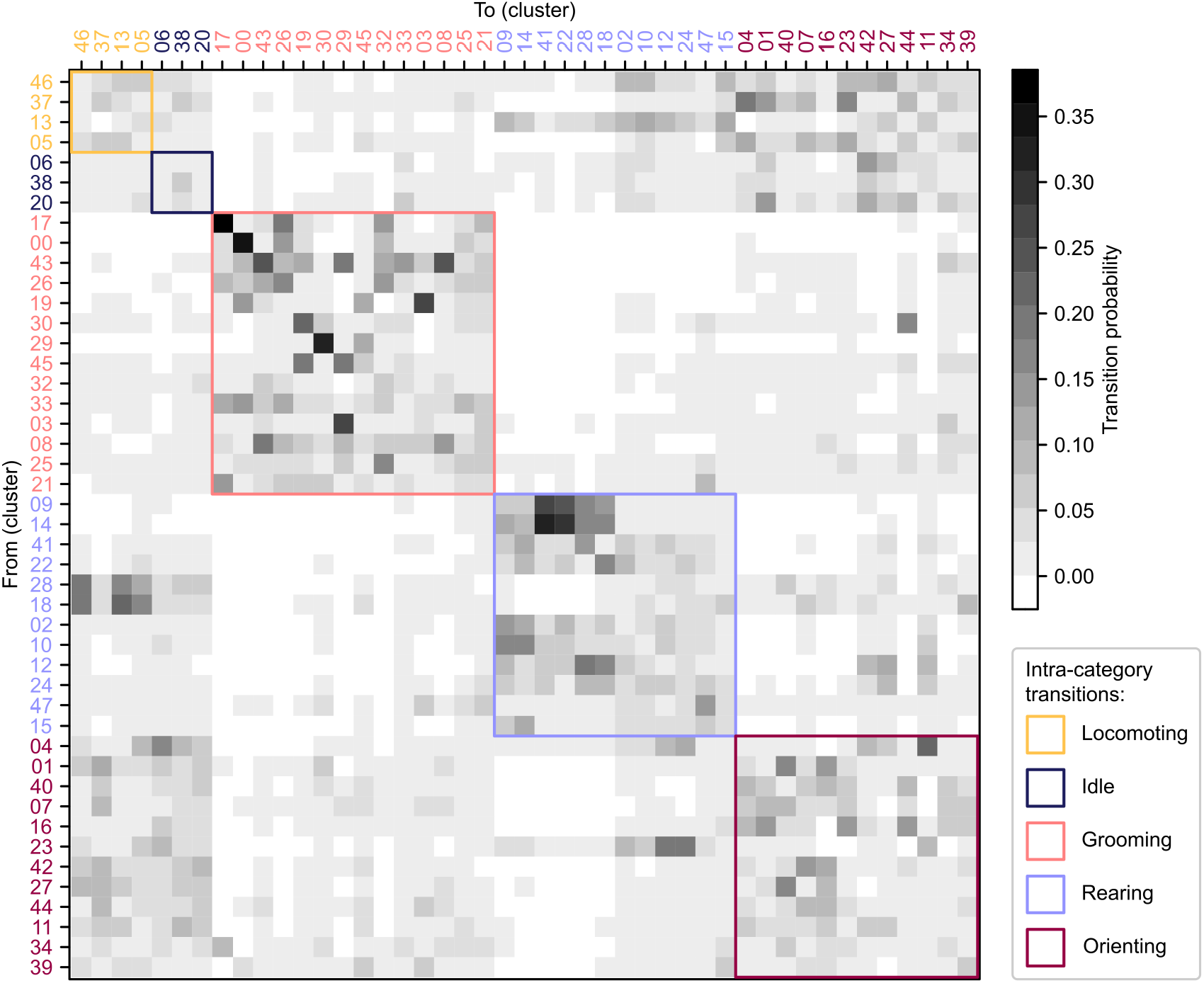
Cluster transition matrix. This graph represents the transition probability between clusters. Cluster numbers are ordered and color-coded according to their dominant main behavioral category (1C). Colored squares highlight intra-category transitions.

**Figure S10:**
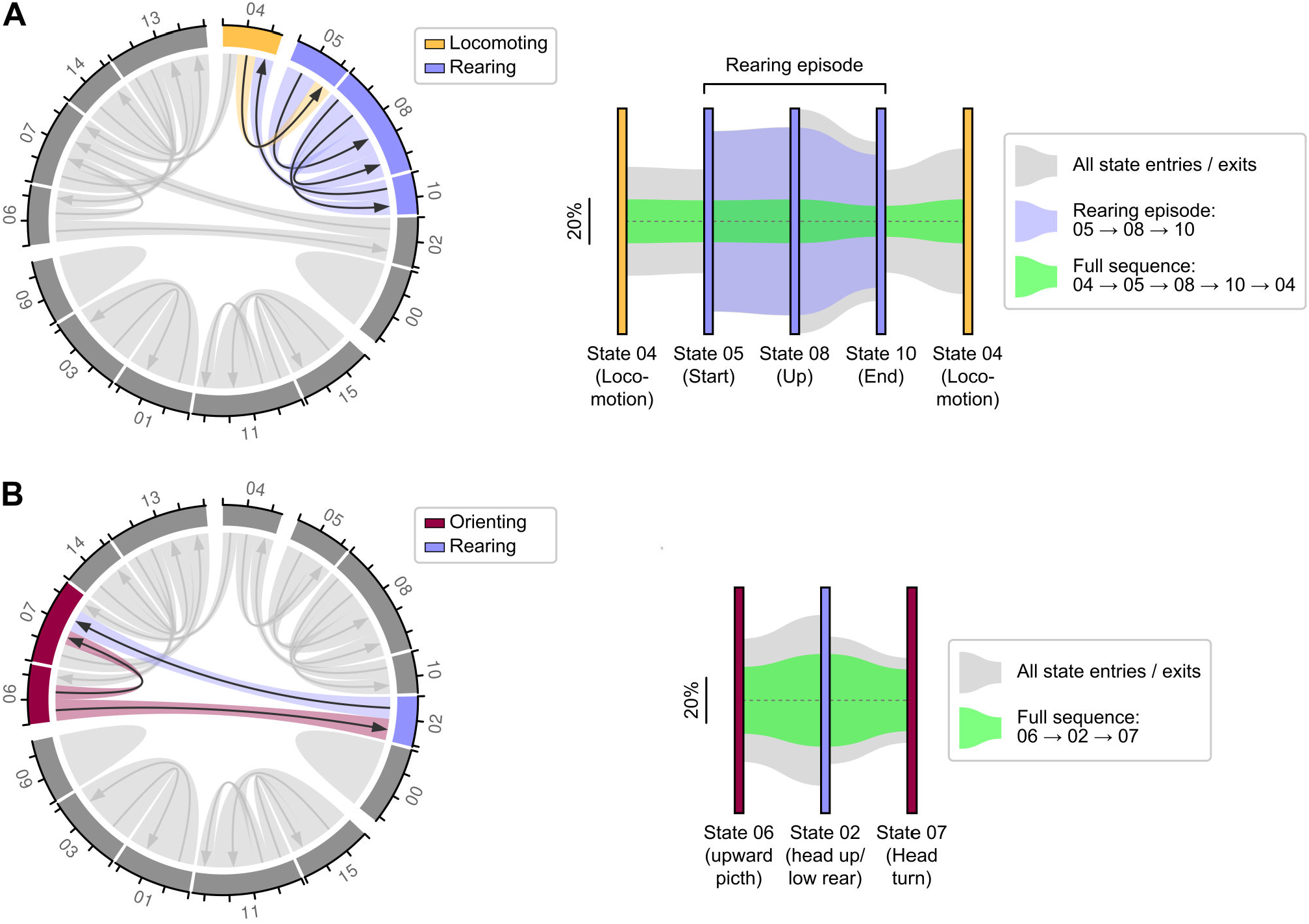
Main inter-category transitions. **A**, Left: chord plot highlighting state transitions corresponding to alternating rearing episodes and locomotor bouts. Right: Sankey diagram showing the proportion of state transitions occurring within isolated rearing episodes (blue) or within a full locomotion-rearing-locomotion sequence (green). The diagram is read from left to right. Colored vertical bars represent states, while shaded areas connecting states represent the proportion of state exits (right side of bars) and entries (left side of bars). **B**, Left: chord plot highlighting state transitions corresponding to sequences during which the animal adopts a pitched-up head posture while keeping its front paws on the ground or close to it (low rear). Right: Sankey diagram showing the proportion of state transitions occurring in the context of this type of sequence (green).

**Figure S11:**
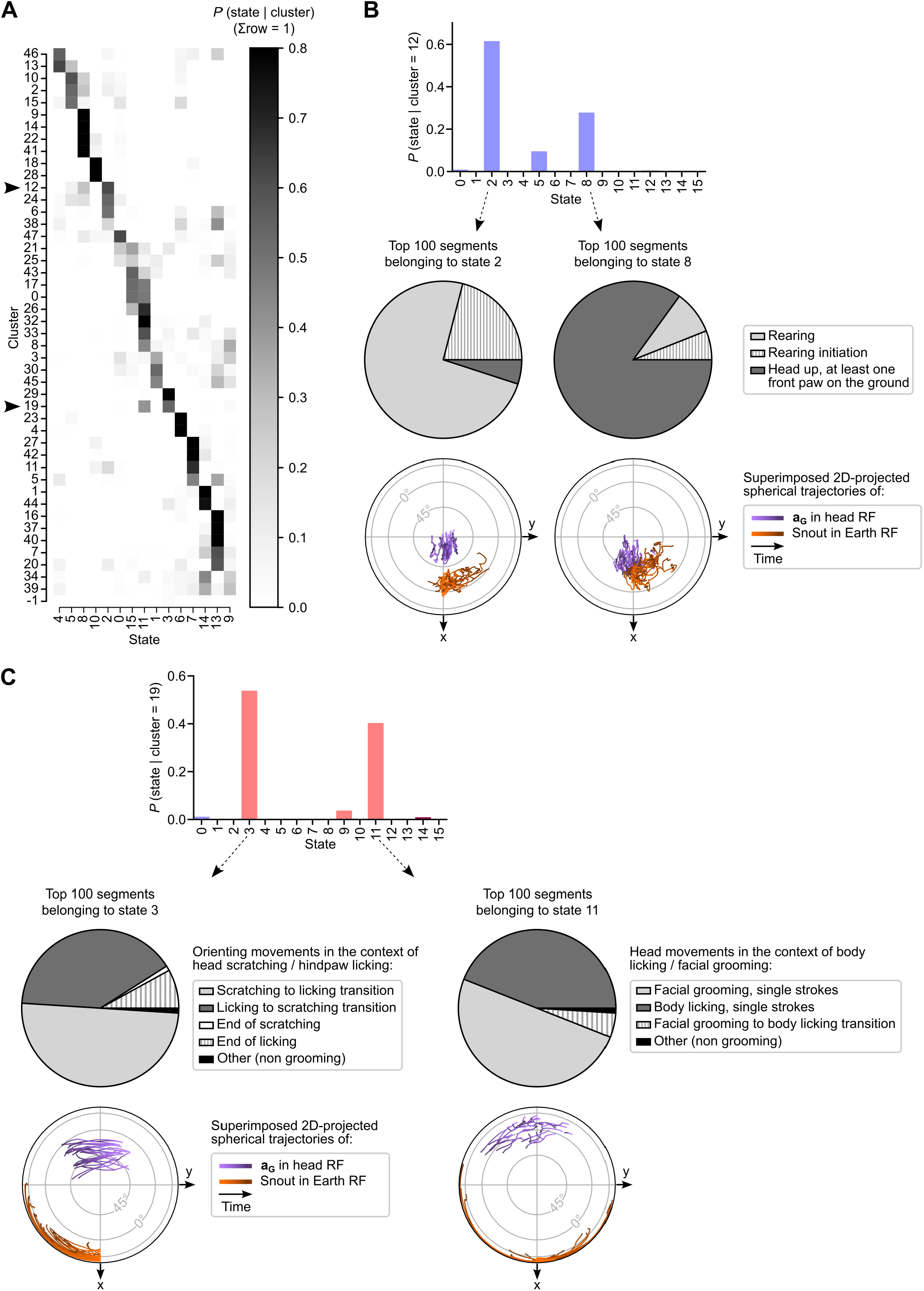
A categorical HMM enables the disambiguation of mixed clusters. **A**, Posterior probability matrix, displaying the likelihood of each state given each cluster, denoted as *P*(*state*|*cluster*. Arrowheads highlight two example clusters (12 and 19) that are associated with multiple states. **B**, Top: Bar plot illustrating the posterior probabilities for cluster 12, showing a strong association with states 2 and 8. Middle: Pie charts displaying the proportion of unique labels assigned to the top 100 segments with the highest *P*(*state* = 2|*cluster* = 12) (left) or *P*(*state* = 8|*cluster* = 12) (right). Note that while the same labels appear in both groups, their proportions differ drastically. Bottom: joint visualization of the trajectories of **a**_**G**_ in the head reference frame (purple) and of the animal’s heading in the Earth reference frame (orange) for the top 30 segments in each group. **C**, Top: Bar plot showing the posterior probabilities for cluster 19, indicating a predominant association with states 3 and 11. Middle: Pie charts displaying the proportion of unique labels given to the top 100 segments with the highest *P*(*state* = 3|*cluster* = 19) (left) or *P*(*state* = 11|*cluster* = 19) (right). Note that labels differ between the two groups, although the same pattern fills are used. Bottom: Joint visualization of the trajectories of **a**_**G**_ in the head reference frame (purple) and of the animal’s heading in the Earth reference frame (orange) for the top 30 segments in each group.

**Figure S12:**
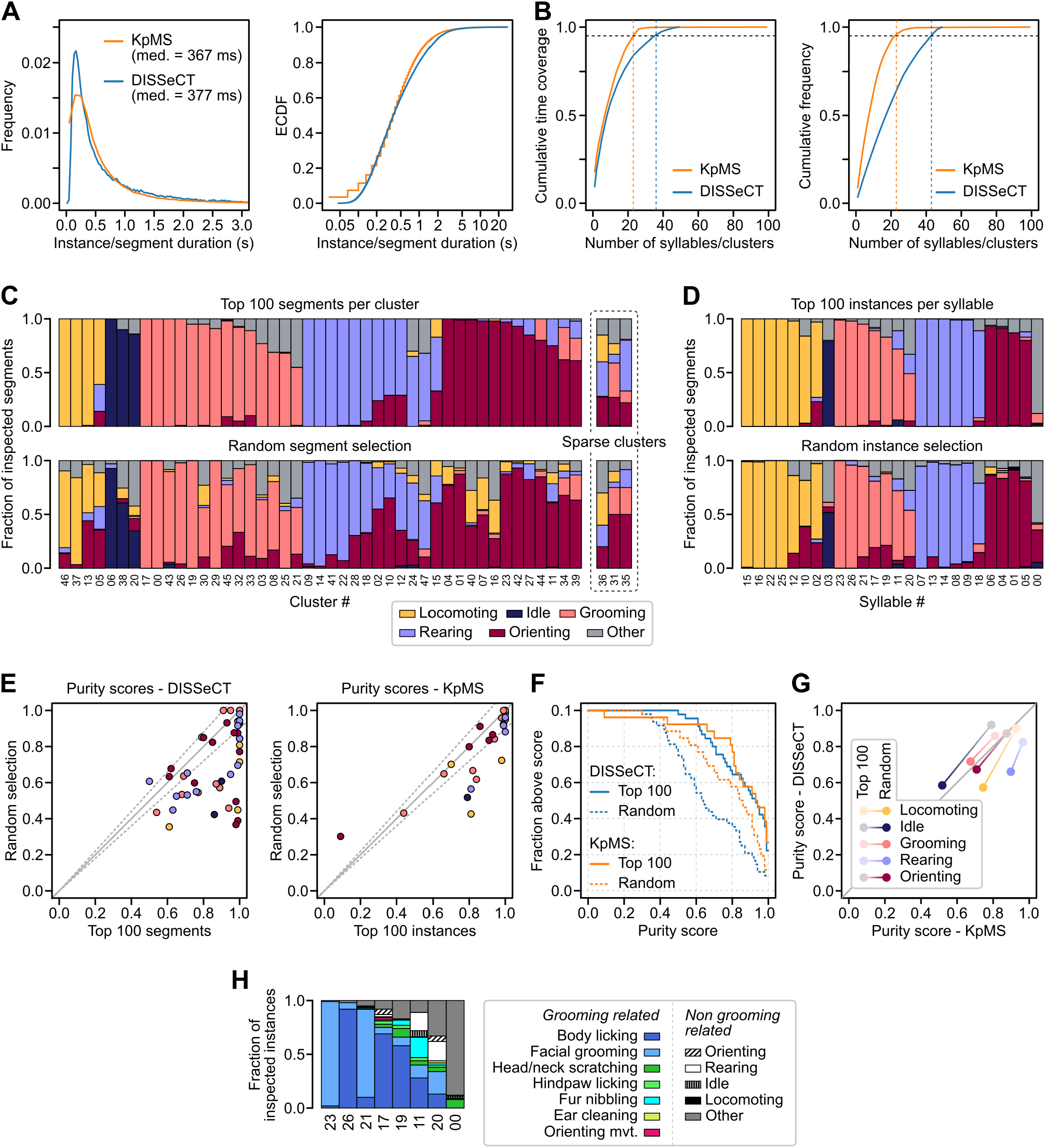
Comparison of the output of DISSeCT and Keypoint-MoSeq (KpMS). **A**, Distribution of segment durations for DISSeCT and syllable instance durations for KpMS, shown as frequency histograms (left) and empirical cumulative distribution functions (ECDFs; right). Median durations are indicated above the histograms. **B**, Left: cumulative time coverage—i.e., the cumulative proportion of total recording time—as a function of the number of clusters (DISSeCT) or syllables (KpMS), sorted by decreasing contribution. A minimum of 23 KpMS syllables (vertical orange dashed line) and 36 DISSeCT clusters (vertical blue dashed line) were required to account for at least 95 % of the total recording time (horizontal black dashed line). Right: cumulative frequency—i.e., cumulative proportion of total instances or segments—as a function of the number of syllables or clusters, sorted by decreasing contribution. At least 23 KpMS syllables and 43 DISSeCT clusters were needed to account for 95 % of all instances and segments, respectively. **C**, Stacked bar plots showing the fraction of segments in each DISSeCT cluster assigned to different behavioral categories, based on manual inspection of either the top 100 segments per cluster (top; same data as in Fig. 1C) or a random sample (bottom; see Methods). **D**, Same as C, but for KpMS syllables. Top: top 100 instances per syllable. Bottom: randomly selected instances (see Methods). **E**, Purity scores for each DISSeCT cluster (left; excluding sparse clusters) and KpMS syllable (right), based on either random samples (*x*-axis) or the top 100 segments/instances (*y*-axis). Each dot represents a cluster or syllable, colored by its dominant behavioral category (as in C-D). The solid diagonal line indicates identity; dashed lines denote ±10 % deviation. **F**, Fraction of clusters or syllables with a purity score above a given threshold, plotted as a function of that threshold. Solid lines: top 100 segments/instances. Dashed lines: random samples. **G**, DISSeCT vs. KpMS purity scores at the behavioral category level. Each behavioral category was associated with a specific set of clusters/syllables, based on the dominant label among their top 100 segments/instances (top panels in C and D). Purity scores were then computed by aggregating the labels of these segments/instances, using either the top 100 most representative ones per cluster/syllable (light colors) or a random global selection (dark colors; see Methods). **H**, Stacked bar plots showing label assignments for the top 100 instances of each grooming-related KpMS syllable.

**Figure S13:**
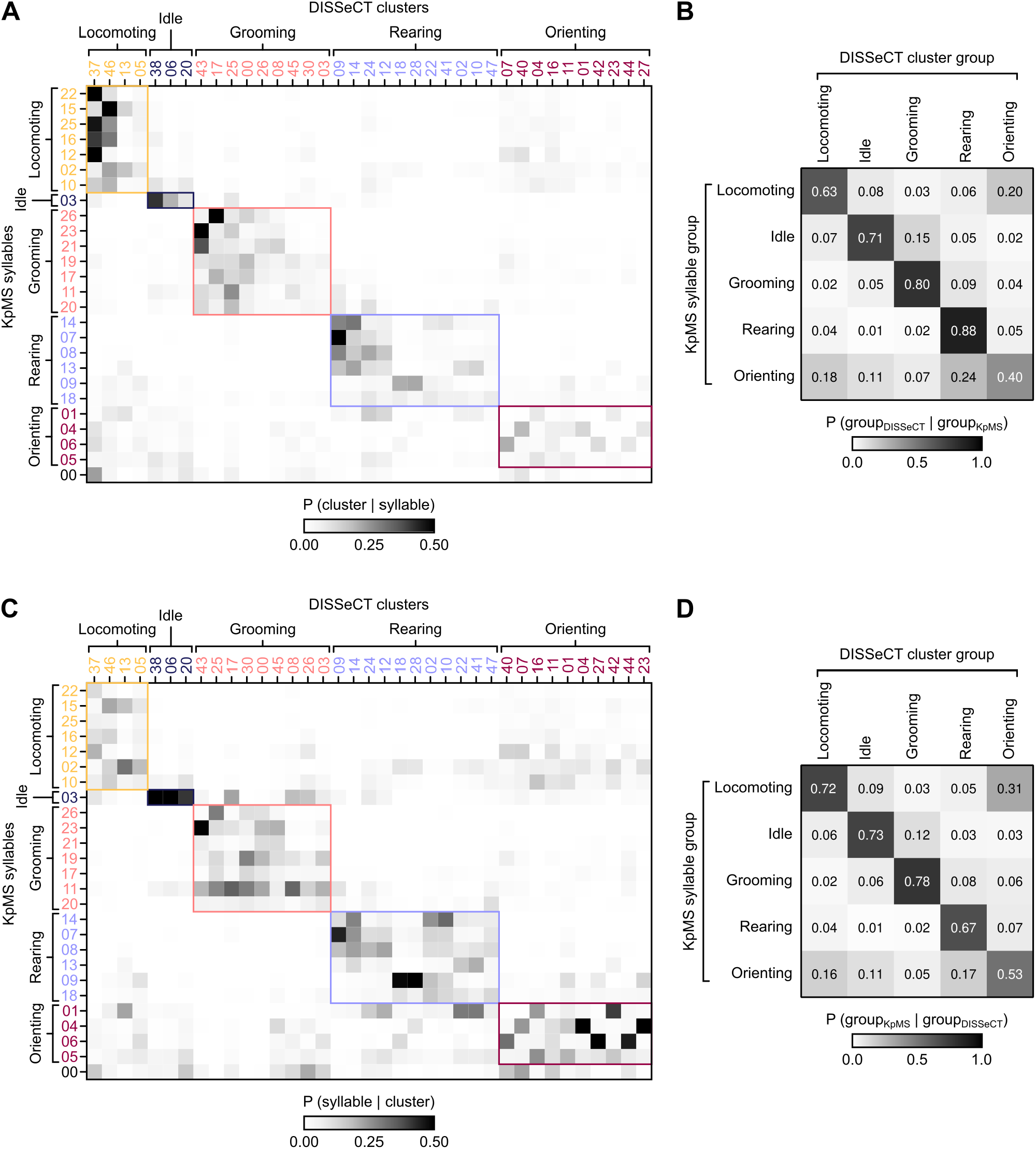
Comparison of cluster and syllable assignment in DISSeCT vs. KpMS. For this analysis, KpMS syllables and DISSeCT clusters accounting for less than 5 % of the total recording duration were excluded. DISSeCT outlier segments and sparse clusters were also excluded. **A**, Row-normalized contingency matrix showing the conditional probability of DISSeCT clusters given KpMS syllables. Both syllables and clusters are grouped by behavioral category. **B**, Same analysis at the behavioral category level. **C**, Column-normalized contingency matrix showing the conditional probability of KpMS syllables given DISSeCT clusters. **D**, Same analysis at the behavioral category level.

**Figure S14:**
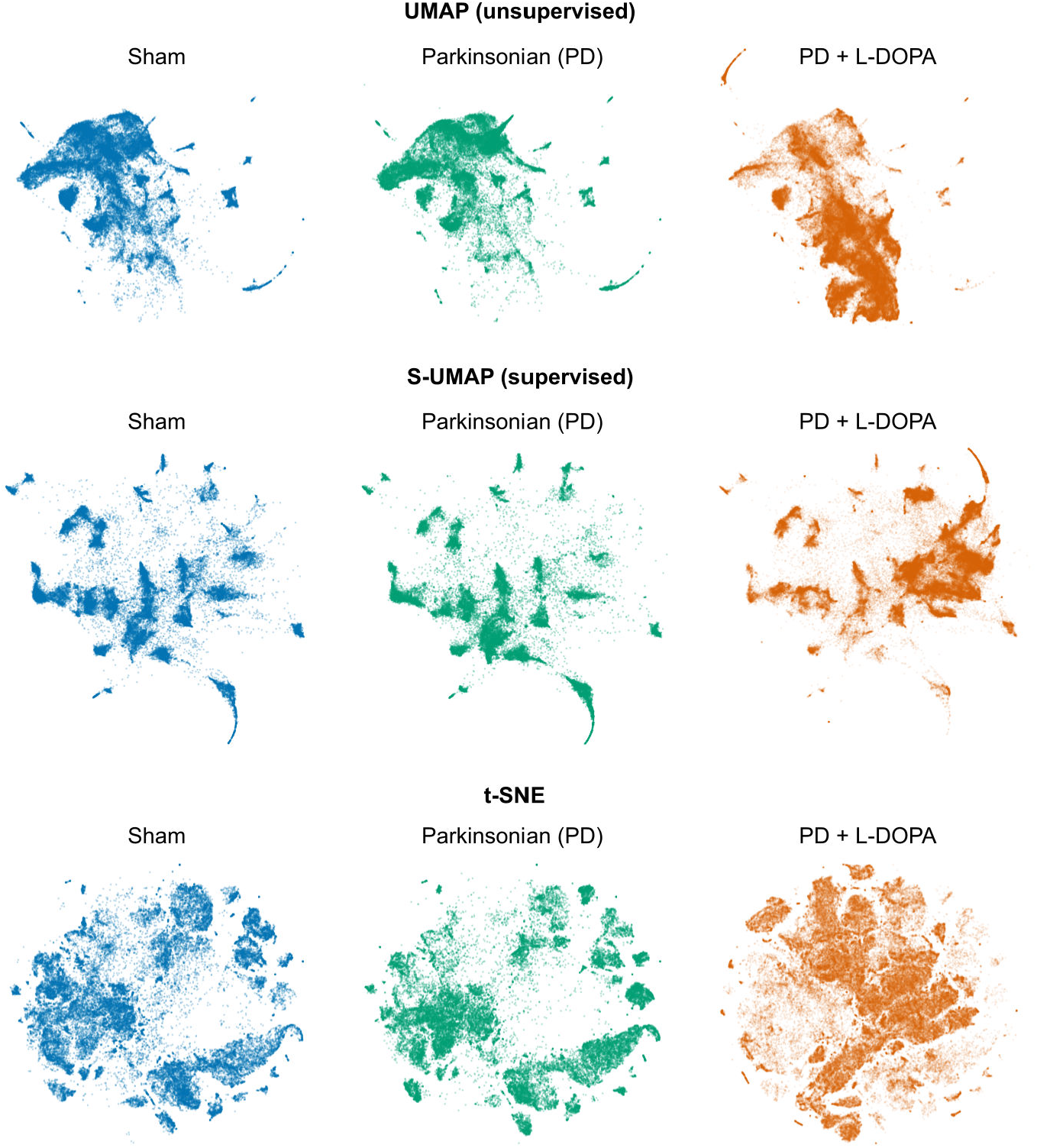
Distribution of segments in two-dimensional embeddings of the standardized feature space for sham and lesioned mice. This figure shows the results of two fully unsupervised embedding methods (UMAP: top; t-SNE: bottom) and one semi-supervised method (S-UMAP: middle), applied to Sham (left, blue) and PD mice before (middle, green) and during (right, orange) L-DOPA treatment. For S-UMAP, cluster identities were derived from a Gaussian Mixture Model (GMM) fitted on data from sham mice. Hyperparameters were set as follows: n_neighbors = 8, n_components = 2 and min_dist = 1e−3 for both UMAP and S-UMAP, as well as target_weight = 1e−9 for S-UMAP; early_exaggeration = 12 and perplexity = 30 for t-SNE.

## Legends of supplementary movies

**Movie S1:**
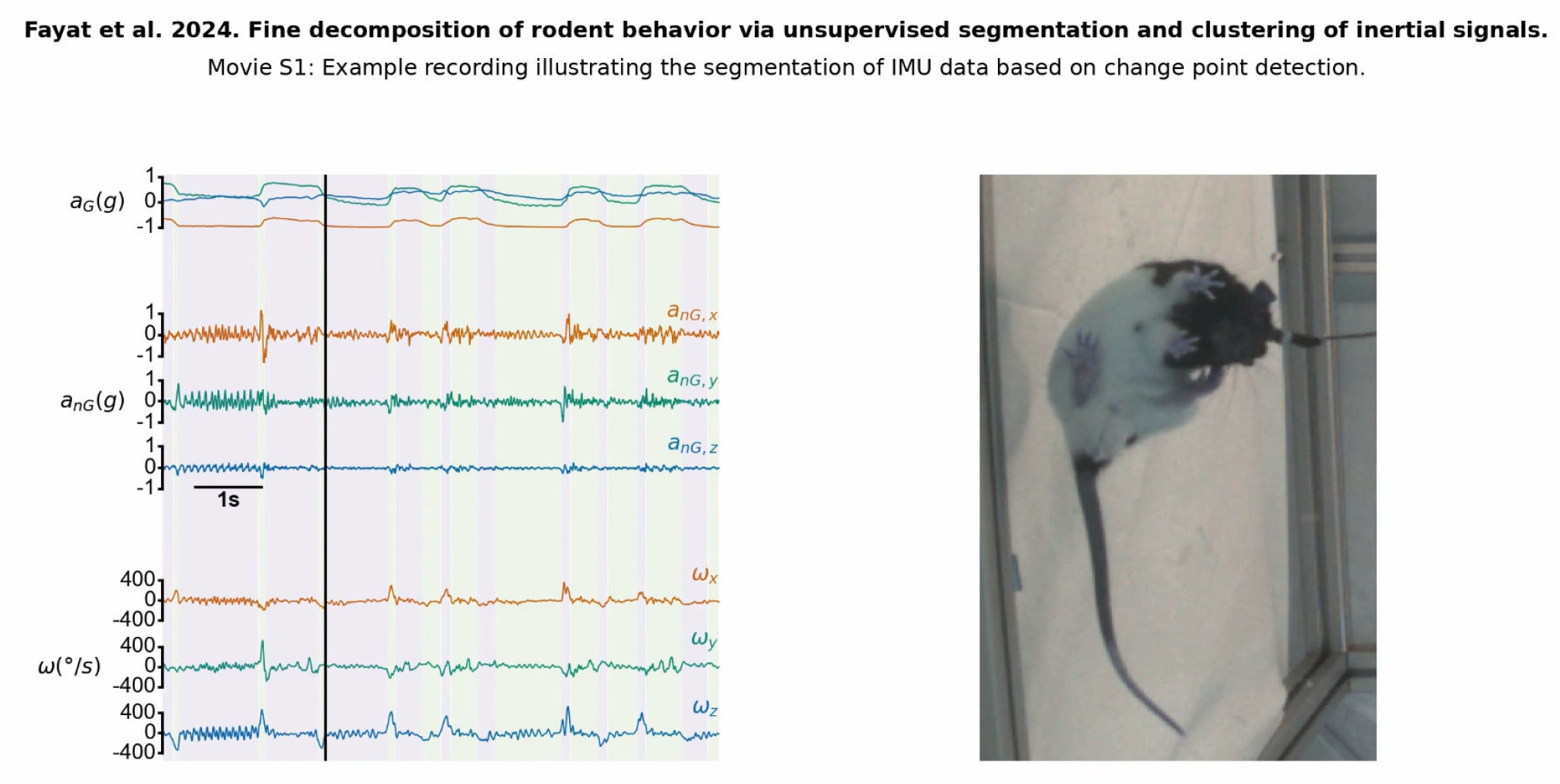
Example recording illustrating IMU data segmentation using change-point detection. Left: Gravitational (***a***_***G***_) and non-gravitational (***a***_***nG***_) components of head acceleration, alongside head angular velocity (***ω***), for an 8-second self-grooming episode. Alternating color bands mark segment boundaries. Notably, rapid changes in head orientation appear as peaks in the angular velocity trace and are accurately identified as distinct segments by the change-point detection algorithm. Right: Simultaneous video recording from beneath a glass floor. IMU and video recordings are synchronized and displayed at half speed.

**Movie S2:**
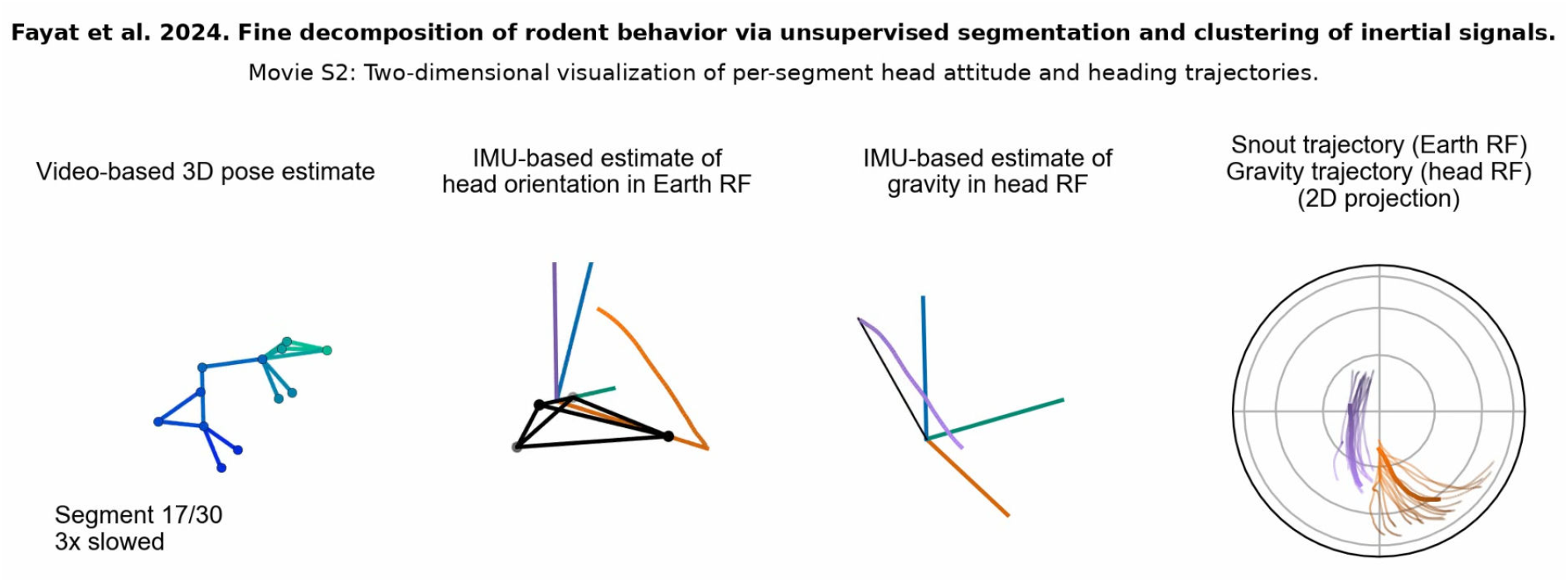
Two-dimensional visualization of per-segment head attitude and heading trajectories. This video details the method used to generate a joint visualization of the trajectories of ***a***_***G***_ in the head reference frame (purple) and of the animal’s heading in the Earth reference frame (orange) for a series of segments (here belonging to the same cluster). See Methods.

**Movie S3:**
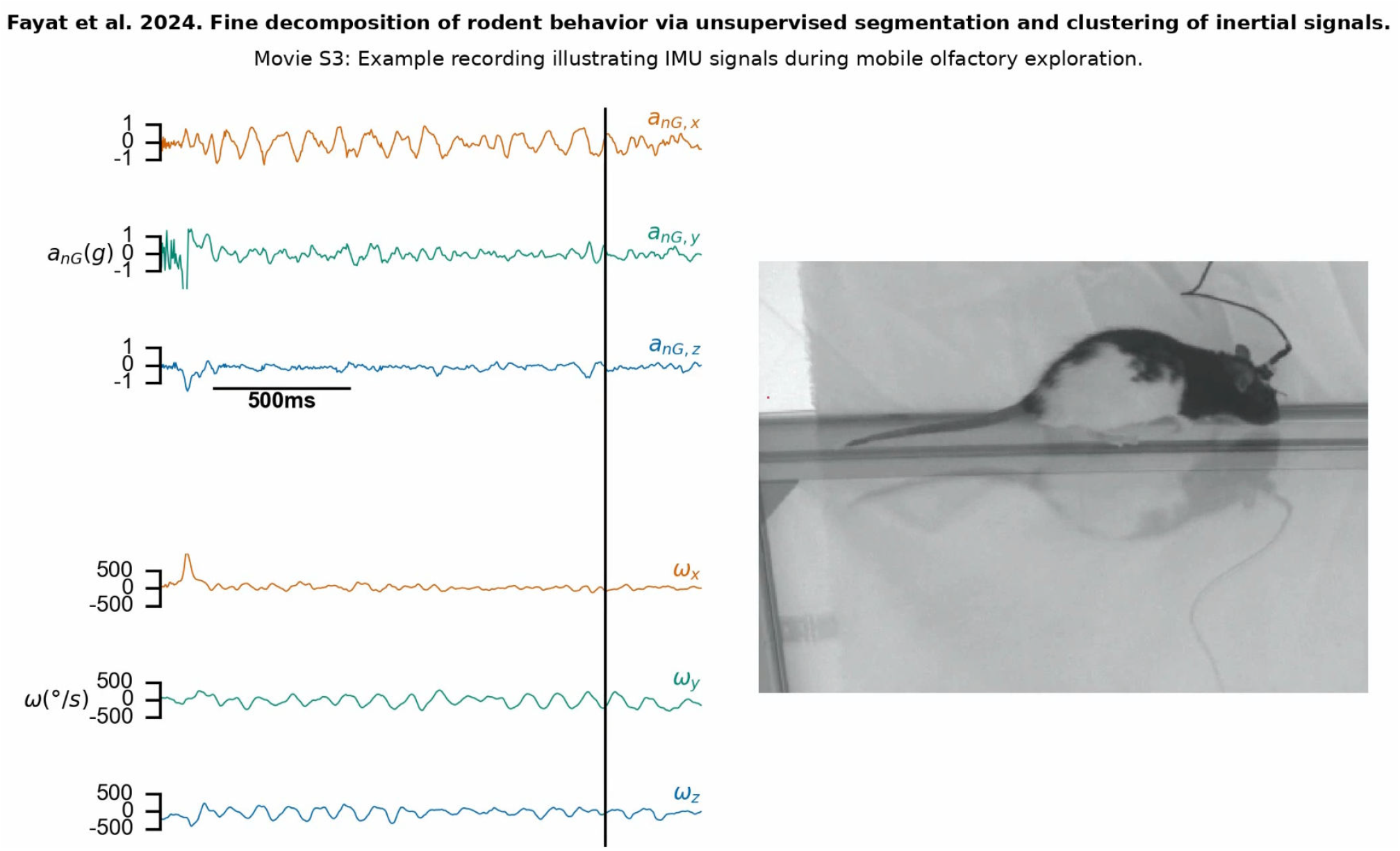
Example recording illustrating IMU signals during mobile olfactory exploration. Left: Non-gravitational component of head acceleration (***a***_***nG***_) and head angular velocity (***ω***) for a 2-second episode of mobile olfactory exploration (see also Fig. 4). Right: Simultaneous video recording. IMU and video recordings are synchronized and displayed at half speed.

**Movie S4:**
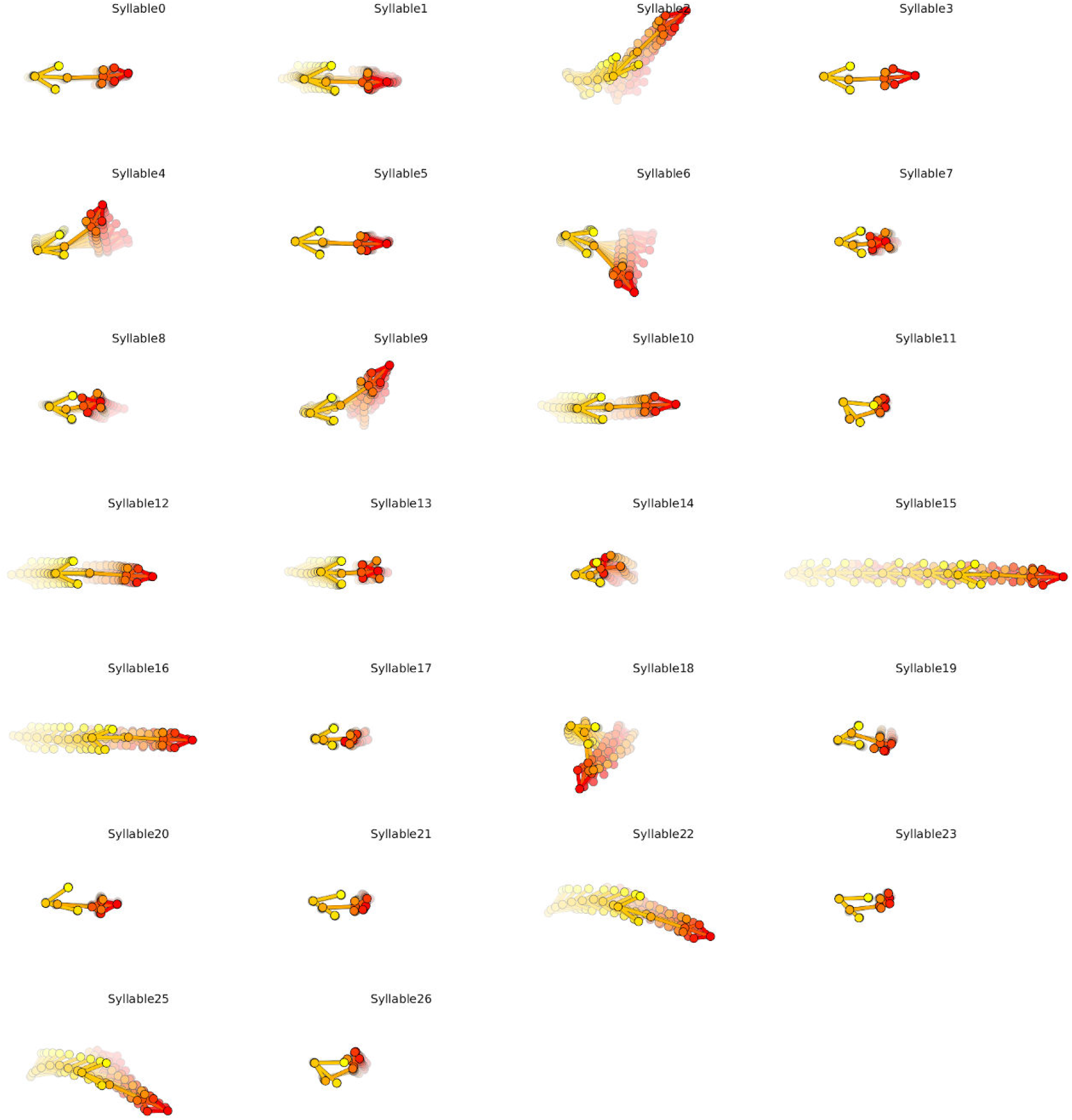
Animated gif showing the representative keypoint trajectories for each syllable projected onto the horizontal (*x, y*) plane, generated by the get_typical_trajectories function of Keypoint-MoSeq.

**Movie S5:**
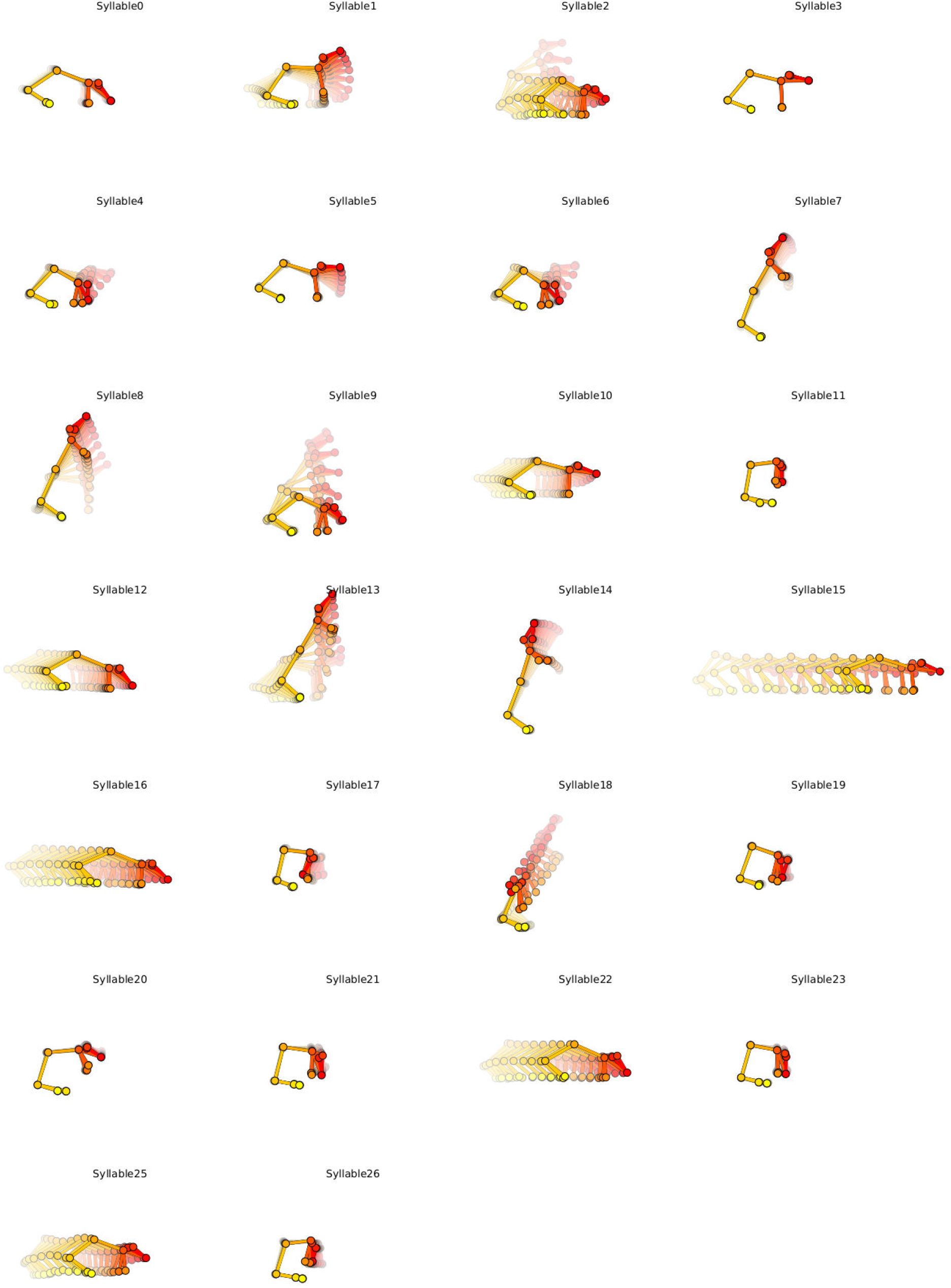
Animated gif showing the representative keypoint trajectories for each syllable projected onto the sagittal (*x, y*) plane, generated by the get_typical_trajectories function of Keypoint-MoSeq.

## References

1. Sousa N, Almeida OF, and Wotjak CT. A hitchhiker’s guide to behavioral analysis in laboratory rodents. Genes, Brain and Behavior 2006;5:5–24.

2. Datta SR, Anderson DJ, Branson K, Perona P, and Leifer A. Computational Neuroethology: A Call to Action. Neuron 2019;104:11–24.

3. Kennedy A. The what, how, and why of naturalistic behavior. Current Opinion in Neurobiology 2022;74:102549.

4. Berman GJ. Measuring behavior across scales. BMC Biology 2018;16:23–34.

5. Ziegler L von, Sturman O, and Bohacek J. Big behavior: challenges and opportunities in a new era of deep behavior profiling. Neuropsychopharmacology 2021;46:33–44.

6. Mathis A, Mamidanna P, Cury KM, et al. DeepLabCut: Markerless Pose Estimation of User-Defined Body Parts with Deep Learning. Nature Neuroscience 2018;21:1281–9.

7. Pereira TD, Aldarondo DE, Willmore L, et al. Fast Animal Pose Estimation Using Deep Neural Networks. Nature methods 2019;16:117–25.

8. Graving JM, Chae D, Naik H, et al. DeepPoseKit, a software toolkit for fast and robust animal pose estimation using deep learning. Elife 2019;8:e47994.

9. Nath T, Mathis A, Chen AC, Patel A, Bethge M, and Mathis MW. Using DeepLabCut for 3D markerless pose estimation across species and behaviors. Nature Protocols 2019;14:2152–76.

10. Deepfly3D, a deep learning-based approach for 3D limb and appendage tracking in tethered, adult Drosophila. eLife 2019;8:1–23.

11. Automated markerless pose estimation in freely moving macaques with OpenMonkeyStudio. Nature Communications 2020;11:1–12.

12. Karashchuk P, Rupp KL, Dickinson ES, et al. Anipose: A toolkit for robust markerless 3D pose estimation. Cell Reports. 36:109730.

13. Dunn TW, Marshall JD, Severson KS, et al. Geometric Deep Learning Enables 3D Kinematic Profiling across Species and Environments. Nature Methods 2021:1–10.

14. Schneider A, Zimmermann C, Alyahyay M, Steenbergen F, Brox T, and Diester I. 3D pose estimation enables virtual head fixation in freely moving rats. Neuron 2022;110:2080–2093.e10.

15. Marshall JD, Aldarondo DE, Dunn TW, Wang WL, Berman GJ, and Ölveczky BP. Continuous Whole-Body 3D Kinematic Recordings across the Rodent Behavioral Repertoire. Neuron 2021;109:420–437.e8.

16. Mimica B, Tombaz T, Battistin C, Fuglstad JG, Dunn BA, and Whitlock JR. Behavioral decomposition reveals rich encoding structure employed across neocortex in rats. Nature Communications 2023;14:3947.

17. Hardcastle K, Marshall JD, Gellis A, et al. Differential kinematic coding in sensorimotor striatum across species-typical and learned behaviors reflects a difference in control. bioRxiv 2023:2023.10.13.562282.

18. Weinreb C, Abdal M, Osman M, et al. Keypoint-MoSeq: parsing behavior by linking point tracking to pose dynamics. Nature Methods 2024;21:1329–39.

19. Nourizonoz A, Zimmermann R, Ho CLA, et al. EthoLoop: automated closed-loop neuroethology in naturalistic environments. Nature methods 2020;17:1052–9.

20. Brown DD, Kays R, Wikelski M, Wilson R, and Klimley A. Observing the unwatchable through acceleration logging of animal behavior. Animal Biotelemetry 2013;1:20.

21. Kays R, Davidson SC, Berger M, et al. The Movebank system for studying global animal movement and demography. Methods in Ecology and Evolution 2022;13:419–31.

22. Chapa JM, Maschat K, Iwersen M, Baumgartner J, and Drillich M. Accelerometer systems as tools for health and welfare assessment in cattle and pigs – A review. Behavioural Processes 2020;181:104262.

23. Garcia-De-Villa S, Casillas-Perez D, Jimenez-Martin A, and Garcia-Dominguez JJ. Inertial Sensors for Human Motion Analysis: A Comprehensive Review. IEEE Transactions on Instrumentation and Measurement 2023;72.

24. Onnela JP. Opportunities and challenges in the collection and analysis of digital phenotyping data. Neuropsychopharmacology 2021;46:45–54.

25. Gu C, Lin W, He X, Zhang L, and Zhang M. IMU-based motion capture system for rehabilitation applications: A systematic review. Biomimetic Intelligence and Robotics 2023;3:100097.

26. Venkatraman S, Jin X, Costa RM, and Carmena JM. Investigating Neural Correlates of Behavior in Freely Behaving Rodents Using Inertial Sensors. Journal of Neurophysiology 2010;104:569–75.

27. Chen CY, Yang CC, Lin YY, and Kuo TB. Locomotion-induced hippocampal theta is independent of visual information in rats during movement through a pipe. Behavioural Brain Research 2011;216:699– 704.

28. Ledberg A and Robbe D. Locomotion-related oscillatory body movements at 6–12 Hz modulate the hippocampal theta rhythm. PloS one 2011;6:e27575.

29. Structured Variability in Purkinje Cell Activity during Locomotion. Neuron 2015;87:840–52.

30. Pasquet MO, Tihy M, Gourgeon A, et al. Wireless inertial measurement of head kinematics in freely-moving rats. Scientific reports 2016;6:35689.

31. Kurnikova A, Moore JD, Liao SM, Deschênes M, and Kleinfeld D. Coordination of orofacial motor actions into exploratory behavior by rat. Current Biology 2017;27:688–96.

32. Dugué GP, Tihy M, Gourévitch B, and Léna C. Cerebellar re-encoding of self-generated head movements. Elife 2017;6:e26179.

33. Klaus A, Martins GJ, Paixao VB, Zhou P, Paninski L, and Costa RM. The Spatiotemporal Organization of the Striatum Encodes Action Space. Neuron 2017;95:1171–1180.e7.

34. Dhawale AK, Poddar R, Wolff SB, Normand VA, Kopelowitz E, and Ölveczky BP. Automated long-Term recording and analysis of neural activity in behaving animals. eLife 2017;6:1–40.

35. Wilson JJ, Alexandre N, Trentin C, and Tripodi M. Three-dimensional representation of motor space in the mouse superior colliculus. Current Biology 2018;28:1744–55.

36. Menardy F, Varani AP, Combes A, Léna C, and Popa D. Functional alteration of cerebello–cerebral coupling in an experimental mouse model of Parkinson’s disease. Cerebral Cortex 2019;29:1752–66.

37. Meyer AF, O’Keefe J, and Poort J. Two distinct types of eye-head coupling in freely moving mice. Current Biology 2020;30:2116–30.

38. Bouvier G, Senzai Y, and Scanziani M. Head movements control the activity of primary visual cortex in a luminance-dependent manner. Neuron 2020;108:500–11.

39. Guitchounts G, Masis J, Wolff SB, and Cox D. Encoding of 3D head orienting movements in the primary visual cortex. Neuron 2020;108:512–25.

40. Angelaki DE, Ng J, Abrego AM, et al. A gravity-based three-dimensional compass in the mouse brain. Nature Communications 2020;11.

41. Mallory CS, Hardcastle K, Campbell MG, et al. Mouse entorhinal cortex encodes a diverse repertoire of self-motion signals. Nature communications 2021;12:671.

42. Fayat R, Delgado Betancourt V, Goyallon T, et al. Inertial Measurement of Head Tilt in Rodents: Principles and Applications to Vestibular Research. Sensors 2021;21.

43. Demars F, Todorova R, Makdah G, et al. Post-Trauma Behavioral Phenotype Predicts the Degree of Vulnerability to Fear Relapse after Extinction in Male Rats. Current Biology 2022;32:3180–3188.e4.

44. Younk R and Widge A. Quantifying defensive behavior and threat response through integrated headstage accelerometry. Journal of Neuroscience Methods 2022;382:109725.

45. Parker PR, Abe ET, Leonard ES, Martins DM, and Niell CM. Joint coding of visual input and eye/head position in V1 of freely moving mice. Neuron 2022;110:3897–3906.e5.

46. Silasi G, Boyd JD, Ledue J, and Murphy TH. Improved methods for chronic light-based motor mapping in mice: automated movement tracking with accelerometers, and chronic EEG recording in a bilateral thin-skull preparation. Frontiers in neural circuits 2013;7:123.

47. Monsees A, Voit KM, Wallace DJ, et al. Estimation of skeletal kinematics in freely moving rodents. Nature Methods 2022;19:1500–9.

48. Shih YH and Young MS. Integrated digital image and accelerometer measurements of rat locomotor and vibratory behaviour. Journal of Neuroscience Methods 2007;166:81–8.

49. Lu J, Zhang L, Zhang D, et al. Development of implantable wireless sensor nodes for animal husbandry and MedTech innovation. Sensors 2018;18:1–12.

50. Wright JP, Mughrabi IT, Wong J, et al. A fully implantable wireless bidirectional neuromodulation system for mice. Biosensors and Bioelectronics 2022;200:113886.

51. Chen M, Liu Y, Tam JC, et al. Wireless AI-Powered IoT Sensors for Laboratory Mice Behavior Recognition. IEEE Internet of Things Journal 2022;9:1899–912.

52. Ouyang W, Kilner KJ, Xavier RM, et al. An implantable device for wireless monitoring of diverse physio-behavioral characteristics in freely behaving small animals and interacting groups. Neuron 2024;112:1764–1777.e5.

53. Massot B, Arthaud S, Barrillot B, et al. ONEIROS, a new miniature standalone device for recording sleep electrophysiology, physiology, temperatures and behavior in the lab and field. Journal of Neuroscience Methods 2019;316:103–16.

54. Groot A de, Boom BJ van den, Genderen RM van, et al. Ninscope, a versatile miniscope for multi-region circuit investigations. eLife 2020;9:1–24.

55. Jonathan P. Newman, Jie Zhang, Aarón Cuevas-López, et al. A unified open-source platform for multimodal neural recording and perturbation during naturalistic behavior. bioRxiv 2023;44.

56. Gomaa W and Khamis MA. A perspective on human activity recognition from inertial motion data. Vol. 35. 28. Springer London, 2023:20463–568.

57. Kleanthous N, Hussain AJ, Khan W, Sneddon J, Al-Shamma’a A, and Liatsis P. A survey of machine learning approaches in animal behaviour. Neurocomputing 2022;491:442–63.

58. Wang G. Machine learning for inferring animal behavior from location and movement data. Ecological Informatics 2019;49:69–76.

59. Bermudez Contreras E, Sutherland RJ, Mohajerani MH, and Whishaw IQ. Challenges of a small world analysis for the continuous monitoring of behavior in mice. Neuroscience and Biobehavioral Reviews 2022;136:104621.

60. Parker PR, Abe ET, Beatie NT, et al. Distance estimation from monocular cues in an ethological visuomotor task. eLife 2022;11.

61. Meyer AF, Poort J, O’Keefe J, Sahani M, and Linden JF. A Head-Mounted Camera System Integrates Detailed Behavioral Monitoring with Multichannel Electrophysiology in Freely Moving Mice. Neuron 2018;100:46–60.e7.

62. Wiltschko AB, Johnson MJ, Iurilli G, et al. Mapping Sub-Second Structure in Mouse Behavior. Neuron 2015;88:1121–35.

63. Luxem K, Mocellin P, Fuhrmann F, et al. Identifying behavioral structure from deep variational embeddings of animal motion. Communications Biology 2022;5.

64. Arlot S, Celisse A, and Harchaoui Z. A kernel multiple change-point algorithm via model selection. Journal of machine learning research 2019;20:1–56.

65. Killick R, Fearnhead P, and Eckley IA. Optimal Detection of Changepoints With a Linear Computational Cost. Journal of the American Statistical Association 2012;107:1590–8.

66. Truong C, Oudre L, and Vayatis N. ruptures: change point detection in Python. 1801.00826 2018:1– 5.

67. Lebarbier É. Detecting multiple change-points in the mean of Gaussian process by model selection. Signal processing 2005;85:717–36.

68. Cappello L and Padilla OHM. Bayesian variance change point detection with credible sets. 2211.14097 2022.

69. Celisse A, Marot G, Pierre-Jean M, and Rigaill G. New efficient algorithms for multiple change-point detection with reproducing kernels. Computational Statistics & Data Analysis 2018;128:200–20.

70. Berman GJ, Choi DM, Bialek W, and Shaevitz JW. Mapping the stereotyped behaviour of freely moving fruit flies. Journal of The Royal Society Interface 2014;11:20140672.

71. Todd JG, Kain JS, and De Bivort BL. Systematic exploration of unsupervised methods for mapping behavior. Physical Biology 2017;14.

72. Lloyd S. Least squares quantization in PCM. IEEE transactions on information theory 1982;28:129–37.

73. Bishop CM. Pattern recognition and machine learning. Springer, 2006.

74. McLachlan GJ, Lee SX, and Rathnayake SI. Finite mixture models. Annual review of statistics and its application 2019;6:355–78.

75. Bouveyron C, Celeux G, Murphy TB, and Raftery AE. Model-based clustering and classification for data science: with applications in R. Vol. 50. Cambridge University Press, 2019.

76. Dempster AP, Laird NM, and Rubin DB. Maximum likelihood from incomplete data via the EM algorithm. Journal of the royal statistical society: series B (methodological) 1977;39:1–22.

77. Hsu AI and Yttri EA. B-SOiD, an open-source unsupervised algorithm for identification and fast prediction of behaviors. Nature Communications 2021;12:5188.

78. Cavagna GA. Storage and utilization of elastic energy in skeletal muscle. Exercise and sport sciences reviews 1977;5:89–130.

79. Skinner R and Garcia-Rill E. The mesencephalic locomotor region (MLR) in the rat. Brain research 1984;323:385–9.

80. Findley TM, Wyrick DG, Cramer JL, et al. Sniff-synchronized, gradient-guided olfactory search by freely moving mice. Elife 2021;10:e58523.

81. Uchida N and Mainen ZF. Speed and accuracy of olfactory discrimination in the rat. Nature neuroscience 2003;6:1224–9.

82. Khan AG, Sarangi M, and Bhalla US. Rats track odour trails accurately using a multi-layered strategy with near-optimal sampling. Nature communications 2012;3:703.

83. Berridge KC, Fentress JC, and Parr H. Natural syntax rules control action sequence of rats. Behavioural Brain Research 1987;23:59–68.

84. Lundblad M, Picconi B, Lindgren H, and Cenci M. A model of L-DOPA-induced dyskinesia in 6-hydroxydopamine lesioned mice: relation to motor and cellular parameters of nigrostriatal function. Neurobiology of disease 2004;16:110–23.

85. Patin R, Etienne MP, Lebarbier E, Chamaillé-Jammes S, and Benhamou S. Identifying stationary phases in multivariate time series for highlighting behavioural modes and home range settlements. Journal of Animal Ecology 2020;89:44–56.

86. Sur M, Hall JC, Brandt J, Astell M, Poessel SA, and Katzner TE. Supervised versus unsupervised approaches to classification of accelerometry data. Ecology and Evolution 2023;13:1–11.

87. Liu J, Bailey DW, Cao H, Cao Son T, and Tobin CT. Development of a Novel Classification Approach for Cow Behavior Analysis Using Tracking Data and Unsupervised Machine Learning Techniques. Sensors 2024;24:4067.

88. Perochon S and Oudre L. Unsupervised Action Segmentation of Untrimmed Egocentric Videos. In: ICASSP 2023 - 2023 IEEE International Conference on Acoustics, Speech and Signal Processing (ICASSP). 2023:1–5.

89. Venkatraman S, Long JD, Pister KSJ, and Carmena JM. Wireless inertial sensors for monitoring animal behavior. Conference proceedings : … Annual International Conference of the IEEE Engineering in Medicine and Biology Society. IEEE Engineering in Medicine and Biology Society. Annual Conference 2007;2007:378–81.

90. Alcacer C, Klaus A, Mendonça M, Abalde SF, Cenci MA, and Costa RM. Abnormal hyperactivity of specific striatal ensembles encodes distinct dyskinetic behaviors revealed by high-resolution clustering. bioRxiv 2024:2024.09.06.611664.

91. Zerveas G, Jayaraman S, Patel D, Bhamidipaty A, and Eickhoff C. A Transformer-based Framework for Multivariate Time Series Representation Learning. Proceedings of the ACM SIGKDD International Conference on Knowledge Discovery and Data Mining 2021:2114–24.

92. Kalueff AV, Stewart AM, Song C, Berridge KC, Graybiel AM, and Fentress JC. Neurobiology of rodent self-grooming and its value for translational neuroscience. Nature Reviews Neuroscience 2016;17:45–59.

93. Lévesque M and Avoli M. The kainic acid model of temporal lobe epilepsy. Neuroscience and Biobehav- ioral Reviews 2013;37:2887–99.

94. De Cordé A, Krzascik P, Wolinska R, Kleczkowska P, Filip M, and Bujalska-Zadrozny M. Disulfiram attenuates morphine or methadone withdrawal syndrome in mice. Behavioural Pharmacology 2018;29:393–9.

95. Suemaru K, Araki H, Kitamura Y, Yasuda K, and Gomita Y. Cessation of chronic nicotine administration enhances wet-dog shake responses to 5-HT2 receptor stimulation in rats. Psychopharmacology 2001;159:38–41.

96. Gomez-Marin A, Paton JJ, Kampff AR, Costa RM, and Mainen ZF. Big behavioral data: Psychology, ethology and the foundations of neuroscience. Nature Neuroscience 2014;17:1455–62.

97. Sebastianutto I, Maslava N, Hopkins CR, and Cenci MA. Validation of an improved scale for rating l-DOPA-induced dyskinesia in the mouse and effects of specific dopamine receptor antagonists. Neurobiology of disease 2016;96:156–70.

98. Coutant B, Frontera JL, Perrin E, et al. Cerebellar stimulation prevents Levodopa-induced dyskinesia in mice and normalizes activity in a motor network. Nature Communications 2022;13:1–17.

99. Sarkar I, Maji I, Omprakash C, Stober S, Mikulovic S, and Bauer P. Evaluation of deep lift pose models for 3D rodent pose estimation based on geometrically triangulated data. 2106.12993 2021.

100. Lopes G, Bonacchi N, Frazão J, et al. Bonsai: An event-based framework for processing and controlling data streams. Frontiers in Neuroinformatics 2015;9:1–14.

101. Zhang L, Dunn T, Marshall J, Olveczky B, and Linderman S. Animal pose estimation from video data with a hierarchical von Mises-Fisher-Gaussian model. In: Proceedings of The 24th International Conference on Artificial Intelligence and Statistics. Ed. by Banerjee A and Fukumizu K. Vol. 130. Proceedings of Machine Learning Research. PMLR, 2021:2800–8.

102. Sabatini AM. Kalman-filter-based orientation determination using inertial/magnetic sensors: Observability analysis and performance evaluation. Sensors 2011;11:9182–206.

103. Harchaoui Z and Cappé O. Retrospective mutiple change-point estimation with kernels. In: 2007 IEEE/SP 14th Workshop on Statistical Signal Processing. IEEE. 2007:768–72.

104. Truong C, Oudre L, and Vayatis N. Selective review of offline change point detection methods. Signal Processing 2020;167:107299.

105. Dickerson AK, Mills ZG, and Hu DL. Wet mammals shake at tuned frequencies to dry. Journal of the Royal Society, Interface 2012;9:3208–18.

106. Josse J and Husson F. Selecting the number of components in principal component analysis using cross-validation approximations. Computational Statistics & Data Analysis 2012;56:1869–79.

107. Schwarz G. Estimating the dimension of a model. The annals of statistics 1978:461–4.

108. McInnes L, Healy J, and Melville J. Umap: Uniform manifold approximation and projection for dimension reduction. arXiv preprint 1802.03426 2018.

109. Fraley C and Raftery AE. Model-based clustering, discriminant analysis, and density estimation. Journal of the American statistical Association 2002;97:611–31.

110. Paxinos G and Franklin KBJ. The Mouse Brain in Stereotaxic Coordinates, Compact, Third Edition: The coronal plates and diagrams. 3rd ed. Academic Press, 2008:256.

111. Hofmann T, Schölkopf B, and Smola AJ. Kernel methods in machine learning. 2008.

112. Sra S. A short note on parameter approximation for von Mises-Fisher distributions: and a fast implementation of I s (x). Computational Statistics 2012;27:177–90.

113. Jolliffe IT and Cadima J. Principal component analysis: a review and recent developments. Philosophical transactions of the royal society A: Mathematical, Physical and Engineering Sciences 2016;374:20150202.

114. Pearson K. LIII. On lines and planes of closest fit to systems of points in space. The London, Edinburgh, and Dublin philosophical magazine and journal of science 1901;2:559–72.

115. Hotelling H. Analysis of a complex of statistical variables into principal components. Journal of educational psychology 1933;24:417.

116. Boyd S and Vandenberghe L. Convex optimization. Cambridge university press, 2004.

117. Cattell RB. The scree test for the number of factors. Multivariate behavioral research 1966;1:245–76.

118. Bouveyron C, Latouche P, and Mattei PA. Exact dimensionality selection for Bayesian PCA. Scandinavian Journal of Statistics 2020;47:196–211.

119. Tipping ME and Bishop CM. Probabilistic principal component analysis. Journal of the Royal Statistical Society Series B: Statistical Methodology 1999;61:611–22.

120. Bouveyron C, Latouche P, and Mattei PA. Bayesian variable selection for globally sparse probabilistic PCA. Electronic Journal of Statistics 2018;12:3036–70.

121. Peel D and MacLahlan G. Finite mixture models. John & Sons 2000.

122. MacQueen J et al. Some methods for classification and analysis of multivariate observations. In: Proceedings of the fifth Berkeley symposium on mathematical statistics and probability. Vol. 1. 14. Oakland, CA, USA. 1967:281–97.

123. Fruhwirth-Schnatter S, Celeux G, and Robert CP. Handbook of mixture analysis. CRC press, 2019.

124. He K, Zhang X, Ren S, and Sun J. Deep Residual Learning for Image Recognition. In: 2016 IEEE Conference on Computer Vision and Pattern Recognition (CVPR). 2016 IEEE Conference on Computer Vision and Pattern Recognition (CVPR). Las Vegas, NV, USA: IEEE, 2016:770–8.

